# Reliable inference of phylogenomic relationship via assembly-based strategy accommodating raw reads and proteins

**DOI:** 10.1101/2024.07.24.604968

**Authors:** Yunlong Li, Xu Liu, Chong Chen, Jian-Wen Qiu, Kevin Kocot, Jin Sun

## Abstract

Phylogenomics has emerged as a transformative approach in systematics, conservation biology, and biomedicine, enabling the inference of evolutionary relationships by leveraging hundreds to thousands of genes from genomic or transcriptomic data. However, acquiring high-quality genomes and transcriptomes necessitates samples with intact DNA and RNA, substantial sequencing investments, and extensive bioinformatic processing, such as genome/transcriptome assembly and annotation. This challenge is particularly pronounced for rare or difficult-to-collect species, such as those inhabiting the deep sea, where only fragmented DNA reads are often available due to environmental degradation or suboptimal preservation conditions. To address these limitations, we introduce VEHoP (Versatile, Easy-to-use Homology-based Phylogenomic pipeline), a tool designed to infer protein-coding regions from diverse inputs, including raw reads (short and long), draft genomes, transcriptomes, and annotated genomes. VEHoP automates the generation of orthologous sequence alignments, concatenated matrices, and phylogenetic trees, streamlining phylogenomic analyses for researchers across disciplines. The tool aims to (1) expand taxonomic sampling by accommodating a wide range of input data types and (2) simplify phylogenomic workflows, making them accessible to researchers with varying levels of bioinformatic expertise. We evaluated VEHoP’s performance using datasets from oysters, catfish, and insects, demonstrating its ability to produce robust phylogenetic trees with strong bootstrap support, outperforming assembly-free methods. Additionally, we applied VEHoP to reconstruct the phylogeny of the enigmatic deep-sea gastropod order Neomphalida, successfully resolving a well-supported phylogenetic backbone for this poorly understood group. VEHoP is freely available on GitHub (https://github.com/ylify/VEHoP), with dependencies easily installable via Bioconda.

## Background

Phylogenetics is now the most fundamental method in evolutionary biology research to understand and illuminate the relationships between organisms. Multiple types of data can be used to infer phylogenetic relationships, including phenotypic and genotypic characteristics. Among them, biological molecules (i.e., nucleic acids and amino acids) are widely used for reconstructing phylogenetic trees. At the early stages of molecular phylogeny, one or a few gene markers were used, such as the mitochondrial cytochrome *c* oxidase subunit I (COI), NADH dehydrogenase subunit 4 (NAD4), nuclear ribosomal RNA genes, or the combination of them (Hao et al. 2015; Ibáñez et al. 2019). With the improvement of sequencing techniques, this was followed by mitogenome-based reconstructions (Donath et al. 2019; Irisarri et al. 2020; Ghiselli et al. 2021). However, these gene trees sometimes failed to reveal the true relationships among taxa due to introgression, different gene evolutional rates between groups, and long-branch attraction (Doolittle and Logsdon Jr 1998; Huynen and Bork 1998; Doolittle 1999; Degnan and Rosenberg 2006). This called for more sophisticated methods for phylogenetics that can address all such issues. Recently, with the development of next-generation sequencers, phylogenetics based on genome-level data (i.e., phylogenomics) has become a focus in many fields (Dunn et al. 2008; Young and Gillung 2020).

It has been shown that taxon sampling is key in reducing errors in phylogenetic inferences (Powell and Battistuzzi 2022). Despite this, in most cases, it is unrealistic to gather sufficient data on all target species to answer the phylogenetic questions. For one, some species inhabit inaccessible environments, such as the deep sea and polar regions, or maybe extremely rare that only one or few specimens are available as long-preserved samples in natural history museums. Also, species distribution in certain groups can be skewed, which leads to biased sampling. In these cases, researchers would have to perform a phylogenetic reconstruction using a dataset lacking some species. If those species happen to represent an important node, the tree topology may be changed based on such an imbalanced taxon-sampling dataset. In addition, most extinct fossil species cannot be sequenced, thus rendering it impossible for molecular phylogenies to include all taxa on the tree of life across evolutionary history (Marshall 2017).

There is no doubt that genome-based phylogeny contains much more information than single or few gene makers (Chang et al. 2011). As next-generation sequencing (NGS) technology advanced, more and more sequenced genomes and transcriptomes have been released to the public at an elevated rate year after year (Turnbull et al. 2023). Nevertheless, many of these datasets were sequenced initially for organelle genome assembly, genome survey, genome annotation, gene expression level analysis, and so on. These are all potential sources for phylogenetics, yet they often remain buried deep in the public database.

The best datasets for phylogenomic analysis are whole genome data from different species (Cheon et al. 2020; Fleming et al. 2023). Yet, the situation is often complicated in practical use. In many groups, only a few well-annotated genomes are available while the rest are transcriptomes and raw Illumina DNA reads. To obtain a genome dataset for phylogenomic analyses from these, multiple steps of bioinformatics analyses must be performed (Liu et al. 2023), which always include quality control of the raw data, draft genome assembly and annotation (Simão et al. 2015). Apart from these, ortholog inference must be performed to identify sequences whose evolutionary history reflects that of the species, which may be the most important step for reliable phylogenomic reconstructions (Yang and Smith 2014; Mongiardino Koch 2021; Lozano-Fernandez 2022). Finally, matrix assembly must be performed, which involves further steps such as alignment, trimming of ambiguously aligned positions, concatenation, and tree reconstruction. The whole workflow is time-consuming and can be confusing for those researchers not from a bioinformatics background (Dylus et al. 2023).

Some tools for phylogenomic analysis can use raw sequencing reads to generate phylogenetic trees, such as Read2Tree (Dylus et al. 2023). However, the reference OMA (“Orthologous MAtrix”) database designated in Read2Tree is not fully customized, and the current procedure for the phylogenetic reconstruction is sophisticated with many manual curation steps. MIKE (Wang et al. 2024) is a MinHash-based and *k*-mer phylogenetic algorithm developed for large-scale next-generation sequencing data. GeneMiner (Xie et al. 2024) is a toolkit developed for phylogenetic marker mining, which extracts markers from transcriptomic, genomic, or other next-generation sequenceing (NGS) or third-generation sequencing (TGS) data. It could be used for multiple gene phylogeny, yet it is still inefficient in phylogenomic analysis due to vague instructions and low numbers of single-copy orthologs extracted.

To address these problems, we here developed a new pipeline which we name ‘VEHoP’ (Versatile, Easy-to-use Homology-based Phylogenomic pipeline). The VEHoP workflow allows different types of datasets as input, including raw reads, genomic DNA assemblies, transcriptomes, well-annotated genomes, or any combinations thereof. After providing these files as the input, users only need to provide a prefix for the run, a path to the database (required if DNA assemblies or transcriptomes are provided), and the optional adjustment of quality control in matrix assembly (e.g., occupancy and alignment threshold, 2/3 and 100 AAs by default, respectively). Alternative analyses can be specified if needed, such as PhyloBayes, ASTRAL, set up occupancy. The output files include single-gene alignments, single-gene trees, a concatenated supermatrix, and results of phylogenetic analyses using the supertree and supermatrix-based approaches.

To assess and benchmark the performance of the VEHoP, we tested it in three benchmarking groups with well-annotated references. Ostreida (the ‘oyster’ order) is a well-studied group of animals in phylum Mollusca with 10 high-quality and well-annotated genomes plus a range of transcriptome datasets, making it an ideal clade for benchmarking the performance and reliability of VEHoP. The other two groups of fish and insects were also selected to verify the feasibility of VEHoP. To further test the applicability of VEHoP in resolving phylogenetic issues, we also used it to analyze a dataset of the gastropod order Neomphalida which is a deep-sea clade of typically small-sized animals. Previously phylogenetic analyses did not fully resolve the internal relationships within this order, due to the lack of high-quality genomes and transcriptomes required by traditional phylogenomic pipelines, and thus the evolutionary relationships among the neomphalidan taxa remained highly contentious. Our results lend support to the VEHoP as a user-friendly, efficient, and accurate workflow.

## Description of VEHoP

### Input files and parameters

The VEHoP pipeline accepts raw reads, draft genome, transcriptome sequencing data, and well-annotated genomes, or any combination of these data types. Raw reads can be NGS or TGS, which could be configured in input with the tab-delimited text (prefix in output; supporting type: NGS, HiFi, ONT, or RNA; read path; read path). It also allows the SRA accession number instead of the local path, which is compiled to download data from NCBI automatically. The raw data will go through a simple, coarse assembly using a *de novo* assembler, such as MEGAHIT (Li et al. 2015) for genomic data (i.e., NGS), Trinity (Haas et al. 2013) for transcriptome data (i.e., RNA), hifiasm (Cheng et al., 2021) for HiFi and Shasta (Shafin et al., 2020) for nanopore reads (i.e., ONT), after quality control and trimming procedures. Other inputs should be in .*fasta* format, but with different suffixes: .*pep.fasta* for proteomes from quality datasets, *.transcript.fasta* for transcriptomes, and *.genomic.fasta* for DNA genomic assemblies. All these assembling procedures can be customized in a VEHoP *.config* file, instead of sophisticated manual operations one by one. In addition, the user also needs to prepare a database for homolog extraction, if genomic or transcriptomic reads or assemblies are provided. The reference database could be a concatenation of protein files suggested from close relatives with well-annotated genomes. By default, VEHoP uses 40 threads (-t 40) throughout, including *de novo* assembly, homolog-inference using miniprot, OrthoFinder processing, matrix assembly and tree construction. During the matrix assembly, VEHoP keeps the quality single-gene alignments with the threshold of alignment length (-l 100) and taxonomy occupancy (2/3, users could adjust manually via setting the minimum samples, -m #s).

### Workflow

The pipeline was coded in Python. All dependencies can be easily installed via Anaconda (Fig. 1) and implemented as follows, except HmmCleaner (Di Franco et al. 2019) which the user can install optionally by the instructions provided in the GitHub repository.

**Fig. 1.**
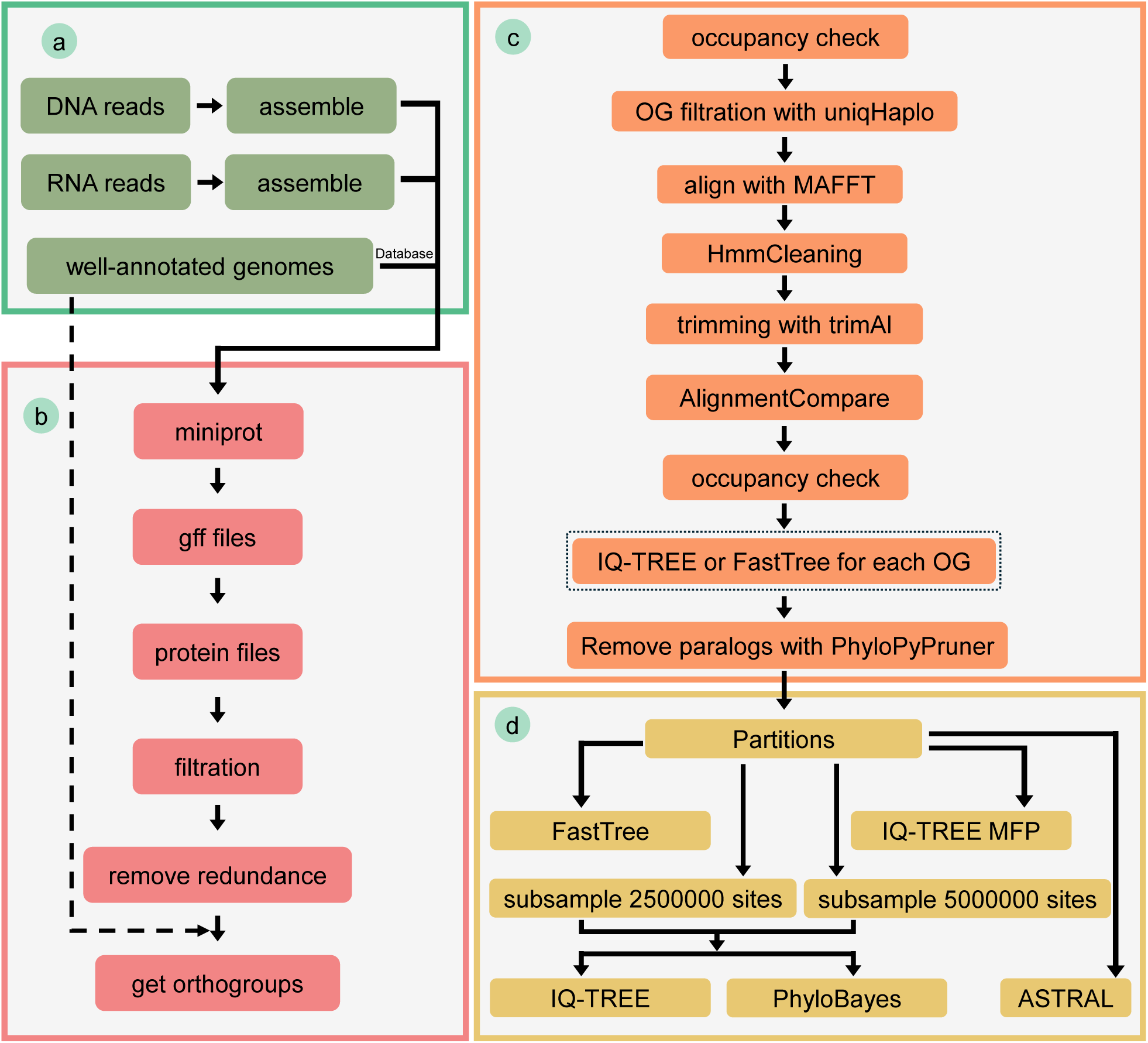
The workflow of the VEHoP pipeline. a) supported input data; b) homolog extraction; c) ortholog inference; d) phylogenetic analyses.

The workflow consists of the following steps (Fig. 1), which can be implemented using a single command:

1) Draft assembly from reads based on the content of a configured text, with SRA download (if applicable) and *de novo* assembly, including Trinity for RNA-seq (NGS in paired-end or single-end mode), Megahit for metagenome from NGS, hifiasm for HiFi reads, and Shasta for nanopore reads; 2) miniprot (Li 2023) is used to map protein sequences from the reference database to the coarsely assembled genomic or transcriptomic data to predict gene models; 3) TransDecoder (Douglas 2018) and embedded Python are used to extract quality proteins based on the predicted gene models (no stop codon in the sequences except for the last one and length above threshold); 4) cd-hit (Fu et al. 2012) is performed to remove redundant sequences with the threshold of 85% similarity; 5) the filtered protein sequences are submitted to OrthoFinder (Emms and Kelly 2019) to identify orthogroups (OGs), with the occupancy assigned by the user (default 2/3, and only orthologs matching the standard will be kept); 6) redundant sequences are removed with uniqHaplo while the remaining sequences are aligned with MAFFT (Katoh and Standley 2013) with default settings; 7) the misaligned regions are removed with HmmCleaner and the aligned files are trimmed with BMGE (Criscuolo and Gribaldo 2010) and trimAL (Capella-Gutiérrez et al. 2009); 8) AlignmentCompare (https://github.com/DamienWaits/Alignment_Compare) is then used to remove sequences shorter than 20 amino acids (AAs), followed by a second occupancy check to make sure all sequences overlap, which is necessary for single-gene tree reconstructions; 9) IQ-TREE or FastTree (default being FastTree) is used to build trees for each filtered OGs. 10) PhyloPyPruner is used to remove paralogs in the filtered alignments; 11) The generated supermatrix is used to reconstruct phylogenetic trees, using IQ-TREE (Minh et al. 2020), FastTree (Price et al. 2010), and PhyloBayes (Lartillot et al. 2013); 12) A random subsample of the initial matrix to 2,500,000 and 5,000,000 sites can also performed for the reconstruction of phylogenetic relationships using IQ-TREE and PhyloBayes. Apart from concatenation-based phylogeny, the pipeline provides a coalescent phylogenetic approach (default: off) implemented via ASTRAL (Mirarab et al. 2014).

### Output files

The output files of the workflow include an initial data matrix in .fasta format, an IQ-TREE tree file, and a FastTree tree file. Apart from the above-mentioned default outputs, the results of ASTRAL and PhyloBayes can also be found in the final output directory if related settings are specified in the commands. If users want to attempt more phylogenetic analyses, they can perform additional custom analyses using the initial data matrix.

## Results

### Benchmark test 1: Oyster dataset

To benchmark the usability and efficiency of the workflow, we collected data from representatives of Ostreida as an example. The datasets include 10 species from Ostreida including *Pinctada fucata*, *Crassostrea hongkongensis*, *C. angulata*, *C. ariakensis*, *C. nippona*, *Ostrea edulis*, *O. denselamellosa*, *C. virginica*, *C. gigas*, *Saccostrea glomerata*, and two species from the closely related order Pectinida (as the outgroup), *Pecten maximus* and *Mizuhopecten yessoensis*. The data included well-annotated genomes, draft genomes from NGS reads, and *de novo* transcriptomes from RNA-seq. The sources of these data are included in the Supplementary Table S1.

We tested our workflow with different datasets, including the following. Dataset 1: well-annotated genomes, whose output was labelled as “reference topology” in Fig. 2; dataset 2: NGS raw reads; dataset 3: transcriptome reads, assembled with Trinity; and dataset 4: a dataset combination including all three types of abovementioned data. For each dataset, the occupancy was set to 2/3, and phylogenetic analyses were performed with two efficient algorithms IQ-TREE (MFP) and FastTree, based on maximum likelihood estimation. The analyses resulted in the same branching order between the reference topology from well-annotated genomes and that from NGS raw reads (Fig. 2a). All bootstrap values reached 100 in these two trees, except for two nodes in NGS reads dataset, with a bootstrap value of 68 within the genus *Crassostrea*. However, the position of *C. nippona* was different from reference topology when using dataset 3 (transcriptomes), though the bootstrap of all nodes reached 100 (Fig. 2a). Furthermore, the same phylogenetic methods were performed on the matrix of 2973 orthologs generated from genome-wide proteins, genome sequences, and transcriptomes, which showed that most of the terminals were clustered by species, except that *C. gigas* was mixed with its most closely related species *C. angulata* (Supplementary Fig. 1).

**Fig. 2.**
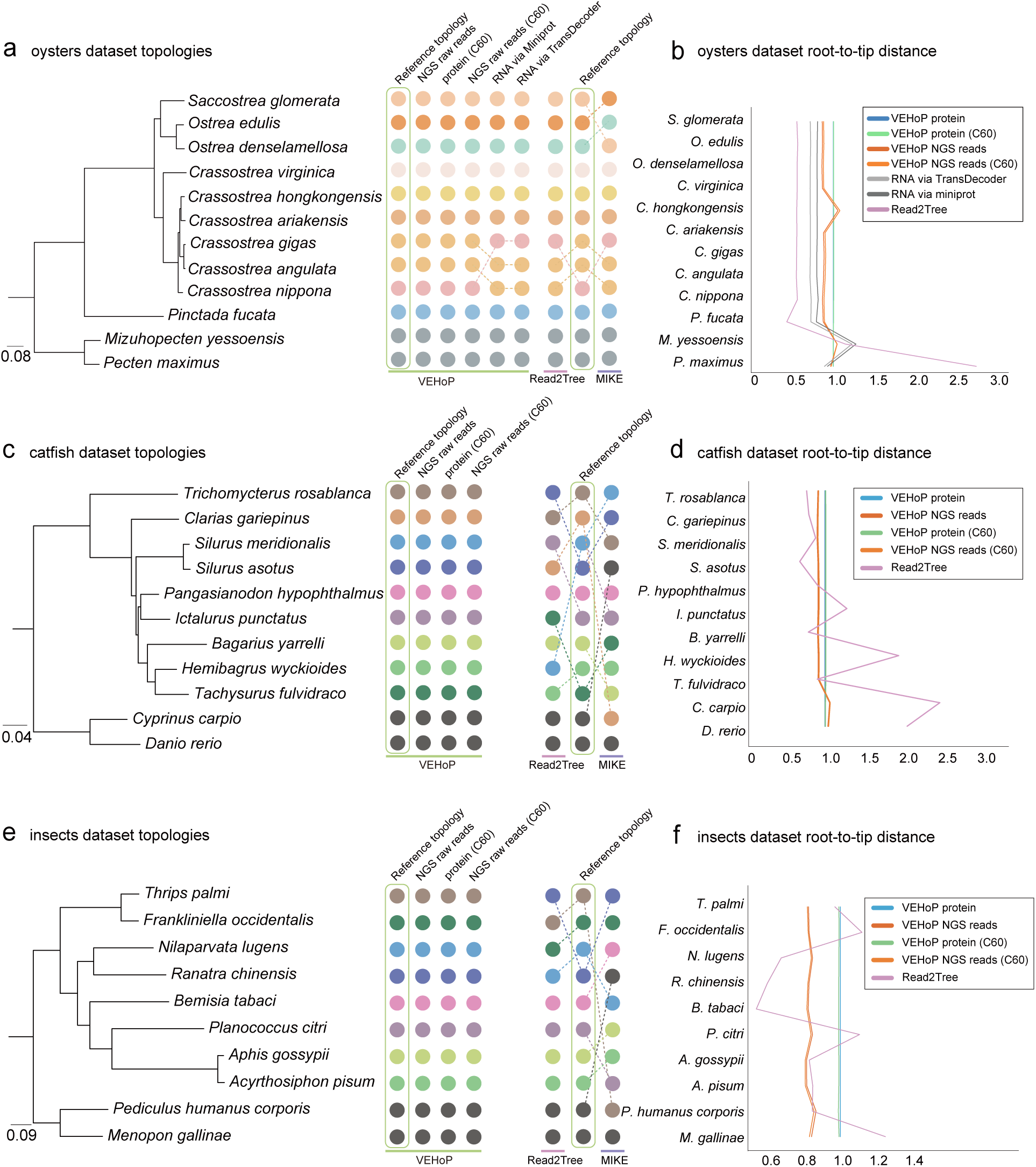
Results of phylogenomic analyses with different datasets, including Ostreida, fish and insects. a) Ostreida dataset topology comparison between different methods and the reference topologies. b) Ostreida dataset root-to-tip distance analysis. c) fish dataset topology comparison between different methods and the reference topologies. d) fish dataset root-ot-tip distance analysis. e) insect dataset topology comparison between different methods and the reference topologies. f) insect dataset root-to-tip distance analysis.

To better understand how much data is sufficient to reconstruct reliable phylogenetic relationships, we subsampled the *C. hongkongensis* data into 2X, 4X, 6X, and 8X of its genome size. Based on these datasets, we performed phylogenetic analyses using IQ-TREE (MFP) and FastTree. The results showed that the pipeline worked well with all the datasets: the branching order of the trees was identical to reference topology, and all node supports were 100% (Supplementary Fig.2). Reduced datasets for every species (1 Gb, 2 Gb, 4 Gb, 6 Gb, and 8 Gb) were also made and phylogenetic analyses conducted (see Supplementary Table S2 for details). The results showed that branch order became unstable for the 1 Gb and 2 Gb datasets, resulting in paraphyly within *Crassostrea*. For datasets larger than 2 Gb, the VEHoP was able to recover phylogenetic relationships from well-annotated genomes, at least at the genus level (Supplementary Fig. 1). The total run time was also recorded for these different datasets to test the performance. For the reduced datasets, it took 4.24, 10.22, 18.60, 25.46, and 27.32 hours to obtain the two tree files, one generated by FastTree and another one generated by IQ-TREE, respectively. As for the full-size mixed dataset, it took VEHoP 54.38 hours to obtain the results, showing the clusters from same species (except the NGS data from *C. angulata*) and the consistent branching order with reference (Supplementary Table S3).

Read2Tree (Dylus et al. 2023) was also performed on the reduced datasets and full-size genomic datasets. Marker genes of the only three mollusc species available on the OMA database, including the oyster *C. gigas*, the octopus *Octopus bimaculoides,* and the true limpet *Lottia gigantea*, were downloaded from the OMA Orthology database as mapping references. For the 1G dataset, Read2Tree took 7.79 hours to get a *.nwk* format tree file, with *Pecten maximus* incorrectly grouped with two reference species from the OMA database (Supplementary Fig. 3). As for the 2G dataset, 19.56 hours were used to generate the tree, yet the position of *C. nippona* was inconsistent with the genome-based tree, though the bootstrap of this node was 100%. In the 4G dataset, a total of 18.5 hours was used, resulting in the same branching order as that in the 2G dataset. For 6G, 8G, and full-size datasets, 21.75, 27.55, and 43.83 hours were used for each dataset, respectively, and they all shared the same branching order as that of the 2G dataset. The total run time for each dataset can also be found in Supplementary Table S3.

MIKE was also performed to benchmark the performance of the VEHoP pipeline with different sizes of datasets, in addition to Read2Tree. In 1G, 2G, 4G, 8G, and full-size datasets, *Saccostrea glomerata* nested with *Crassostrea* or within *C. virginica*, causing paraphyly. Only in the 6G dataset, the topology was well-resolved and consistent with the current understanding of oyster phylogeny at the genus level (Li et al. 2021) (Supplementary Fig. 4). The run time of MIKE for different sizes of datasets can be found in Supplementary Table S4.

The root-to-tip distances for each species were calculated using various tree files to assess tree quality. The distances of each tip in reference topology (well-annotated genomes) were employed to normalize the corresponding distances in other trees (Supplementary Table S5). The findings showed the similar root-to-tip distances to reference topology, whereas the trees produced by Read2Tree displayed significant variance compared to the other results (Fig. 2b). The root-to-tip distances in MIKE is not applicable for quantification.

### Benchmark test 2: Catfish dataset

The fish datasets include 9 catfishes from the order Siluriformes: *Bagarius yarrelli*, *Clarias gariepinus*, *Hemibagrus wyckioides*, *Ictalurus punctatus*, *Pangasianodon hypophthalmus*, *Silurus asotus*, *Silurus meridionalis*, *Tachysurus fulvidraco* and *Trichomycterus rosablanca*. The common carp *Cyprinus carpio* and the zebrafish *Danio rerio* were used as outgroups. The datasets include well-annotated genomes, NGS raw reads and public draft assemblies from NCBI. The source of these data can be found at Supplementary Table S1.

Three datasets were used in this benchmark test, including dataset 1: well-annotated genomes dataset 2: NGS raw reads. VEHoP, Read2Tree and MIKE were applied on dataset 2, whose topologies can be found in Fig. 2c. Other than that, an additional IQ-TREE (MFP) procedure was also performed on the matrix generated by Read2Tree. IQ-TREE (MFP) and IQ-TREE (C60) were performed based on the matrix generated by VEHoP in both two datasets. And for comparing, the topology generated by IQ-TREE (MFP) based on well-annotated genomes, in this case, dataset 1, was chosen to be the reference topology. VEHoP showed a great consistence and stability in NGS reads (dataset 2), while Read2Tree and MIKE both resulted in rather different topologies compared to the reference topology (Fig. 2c). The topology generated by Read2Tree showed that *H. wychioides* grouped together with the outgroup species, and *Sasotus* came to the basal position of the ingroup instead of *T. rosablanca*. The genus *Silurus* was recovered as paraphyletic. In MIKE, the outgroup species *C. carpio* nested within catfishes. And the genus *Silurus* became basal instead of *T. rosablanca*. *C. gariepinus* nested deep inside of the ingroup catfish species, while in the reference topology, it was positioned at the location of the secondary basal node. Root-to-tip distance was also calculated and normalized for each tree file generated (Supplementary Table S5); all results generated by VEHoP demonstrated a high level of consistency (Fig. 2d). All original tree topologies can be found in Supplementary Fig. 5.

### Benchmark test 3: Insect dataset

The insect datasets include 8 species from the superorder Condylognatha, which comprises of two orders: Thysanoptera (thrips) and Hemiptera (true bugs). These species are *Acyrthosiphon pisum*, *Aphis gossypiii*, *Bemisia tabaci*, *Frankliniella occidentalis*, *Nilaparvata lugens*, *Planococcus citri*, *Ranatra chinensis* and *Thrips palmi*. And two species from Psocodea were selected as outgroups: *Pediculus humanus corporis* and *Menopon gallinae*. The datasets used in this study also include well-annotated genomes, NGS raw reads and public genome assemblies from NCBI, whose data source can also be found at Supplementary Table S1.

The dataset composition in this benchmark test was the same as that in benchmark test 2: including dataset 1: well-annotated genomes; dataset 2: NGS raw reads. The methodology used was the same as for the catfish case study. The results generated by VEHoP shared the same branch order as the reference topology (i.e., inferred from well-annotated genomes), who successfully recovered the monophyletic relationship of Thysanoptera and Hemiptera. But in

Read2Tree and MIKE, they both resulted in inconsistent branch orders compared to reference topology (Fig. 2e) In Read2Tree, Hemiptera was successfully recovered as monophyletic, yet *N. lugens* was assigned to Thysanoptera incorrectly. And in MIKE, *M. gallinae* nested within Hemiptera as an outgroup species. Besides, both Hemiptera and Thysanoptera were not analyzed as monophyletic groups. The inconsistency can also be found in the root-to-tip distance results in Fig. 2f. All original tree topologies can be found in Supplementary Fig. 5.

### Case study: Neomphalidan snails

The molecular phylogeny of deep-sea endemic neomphalidan gastropods has long been contentious, partially due to insufficient sampling, small body size and tissue quantity, and lacking many sequences. Here, we applied VEHoP on the original Illumina sequencing dataset (see Supplementary Table S6 for details) used to assemble the mitochondrial genomes from a previous study (see Zhang et al., 2024), which generated a matrix consisting of 1899 orthologs with an occupancy of 2/3. In addition, to improve taxon sampling, we newly sequenced a specimen of *Neomphalus fretterae* (collected from Tempus Fugit vent field, Galápagos Rift, 0°46.1954’N / 85°54.6869’W, 2561 m deep, R/V *Falkor (too)* cruise FKt231024, remotely operated vehicle (ROV) *SuBastian* dive #609, 2023/Nov/02) following the same methods as Zhang et al. (2024). Four species of Cocculinida (*Cocculina enigmadonta, C. tenuitesta, C. japonica, C. subcompressa*), the sister-order of Neomphalida were used as outgroups, as well as the more distantly related vetigastropod snails *Tristichotrochus unicus* and *Steromphala cineraria*. The data of *C. enigmadonta*, *C. tenuitesta*, *Lamellomphalus manusensis*, *Lirapex politus*, *Symmetriapelta wreni*, *Melanodrymia laurelin*, *Melanodrymia telperion*, Neomphalidae gen *et* sp. Hatoma *sensu* Zhong et al., 2022, *Nodopelta heminoda*, and *Symmetriapelta becki* were gathered from previous studies, which were used to assemble mitochondrial genomes for phylogeny (Zhong et al. 2022; Zhang et al. 2024).

We first attempted to reconstruct the molecular phylogeny of Neomphalida using mitochondrial genomes with multiple models in IQ-TREE, including MFP, C20, C40, and C60 based on the matrices from Zhang et al. (2024). This revealed two distinct tree branching orders with nearly equal support from different sequencing matrices (see Supplementary Fig. 6, confirming the same situation encountered also in a previous study (Zhang et al., 2024). We then conducted multiple phylogenetic analyses through VEHoP based on the assemblies of the abovementioned datasets, including IQ-TREE with the MFP model, Site-specific frequency models (including C20, C40, and C60), and FastTree. All these analyses resulted in the same tree branching order with maximum support in each node, except for one node in Peltospiridae which had the bootstrap value of 85 in the C20 model (Fig. 3). Apart from VEHoP, Read2Tree and MIKE were also performed on the same dataset of Neomphalida. However, these two methods were unable to resolve a consistent topology, even at the order level (Supplementary Fig. 7).

**Fig. 3.**
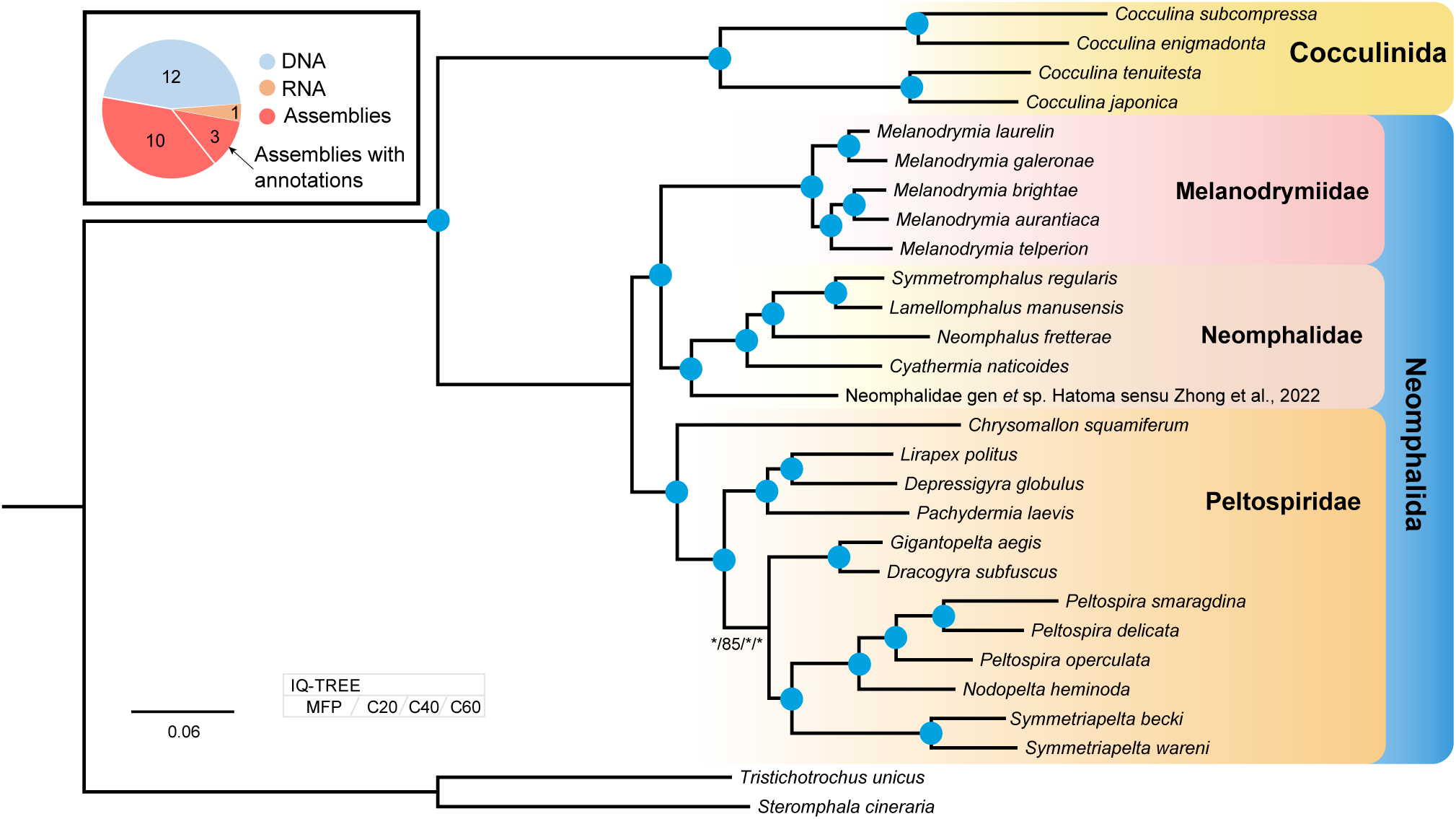
Results of phylogenomic analysis using VEHoP on short Illumina sequencing data from Neomphalida. Nodes with blue dots indicate maximal support in all analyses using different methods. *Neomphalus fretterae* was newly sequenced in this study.

## Discussion

We present VEHoP, a new pipeline for phylogenomic analyses with the flexibility of using publically genomic assemblies, well-annotated genomes, NGS raw reads, RNA-seq raw reads, or a combination of these data. This workflow allows users to reconstruct phylogenetic trees with one single command, significantly lowering the technical hurdle for researchers to carry out phylogenomic inferences. VEHoP is able to reconstruct congruent and robust relationships among taxa using fragmented draft genomes that were rapidly assembled from NGS reads, with results comparable with trees generated from well-annotated genome datasets.

Currently, most available phylogenomic pipelines are based on protein datasets (Kocot et al. 2011; Sun et al. 2021), which require cumbersome steps and are time-consuming to prepare. To obtain high-quality protein files, high-quality DNA sequencing data is inevitably needed. Furthermore, it is necessary to conduct genome assembly to get a contig- or scaffold-level draft genome, followed by gene model prediction. This workflow usually takes several days just for one single species even with ample computing resources.

There is a vast amount of data in public databases, including unannotated genomes and raw NGS reads (genome skimming projects previously used in organelle assemblies or genome surveys), which have been underutilized in phylogenomic studies. Understandably, these data vary in quality and coverage, and thus it has been challenging to use them for phylogenetic analysis. With VEHoP, however, researchers can extract homologs from these genomic data at ease, with the potential to greatly enhance taxon sampling and produce a more robust and consistent tree topology in phylogenetic analyses. As an example, we generated pie charts for major lophotrochozoan animal phyla to show the potential of these ‘buried’ data in phylogenetics based on NCBI data (Fig. 4 and Supplementary Table S7, data up to May 2024). Among Mollusca, for example, there are only 286 species with genome assemblies (only a small fraction of these is annotated) while an additional 896 species have transcriptomic data. These two data types are mostly commonly used source data for phylogenomic analysis. With VEHoP, we can further include 325 species which lack both genome and transcriptome data but with DNA genomic data, greatly expanding the taxon coverage.

**Fig. 4.**
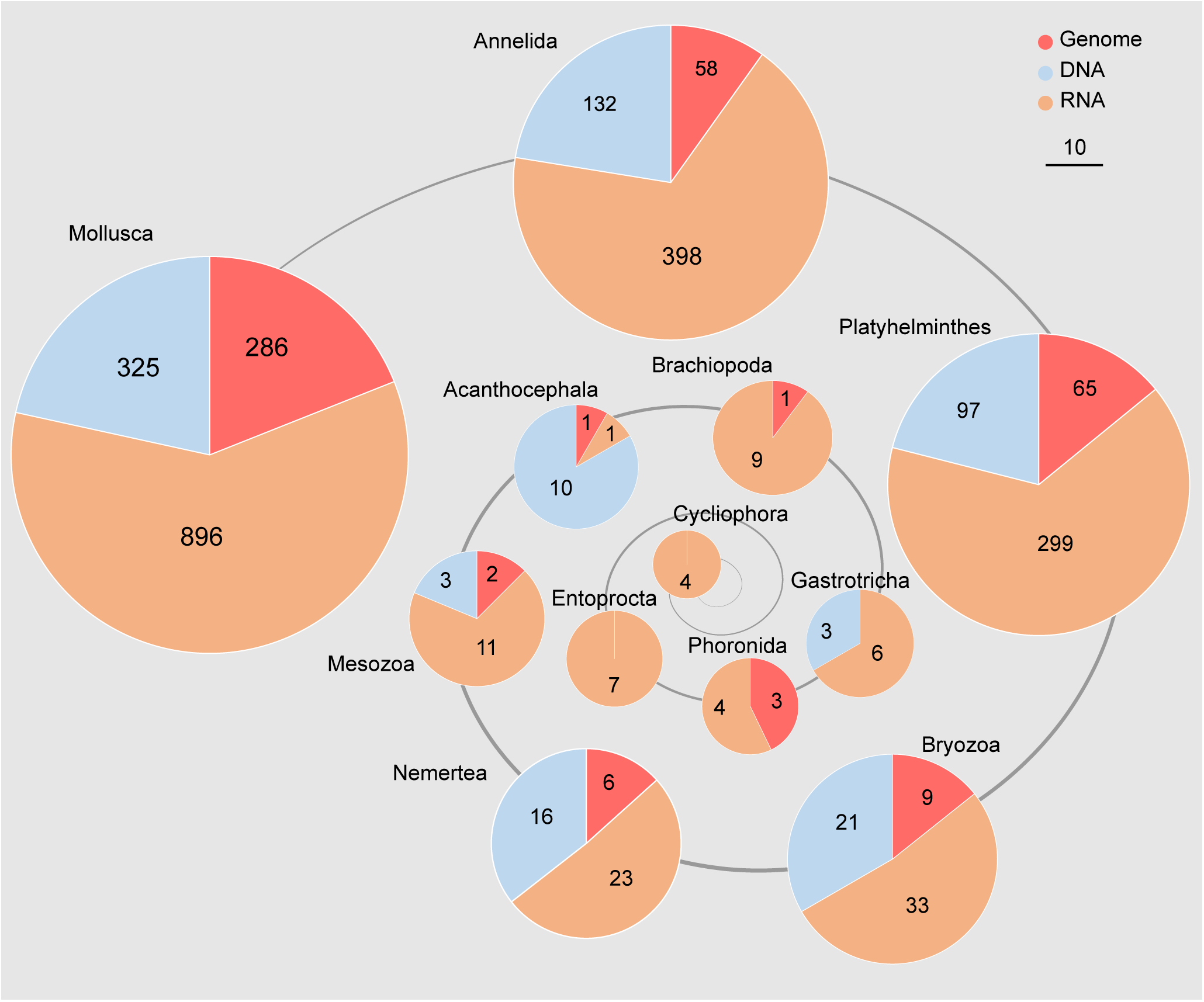
Available phylogenomic resources for major phyla in the major animal clade Lophotrochozoa enumerated in terms of the number of taxa with published genomes (red), RNA-seq datasets (orange), and DNA genomic assemblies (blue). Sizes of the circles are proportional to the number of species in each phylum.

In our benchmarking study using various data types from oysters (benchmark test 1), VEHoP showed a high speed and accuracy in inferring phylogeny. The branching order inferred based on unannotated genomic data was the same as that based on well-annotated genomes, though not all node support reached 100%. For the RNA data, we attempted two strategies: 1) extracting homologs directly from assembled transcripts with miniprot; 2) predicting proteins with TransDecoder. Those two strategies resulted in the same branching order, and each node reached 100% support. However, the branching order from this analysis differed from those based on well-annotated genomes. This discrepancy was probably due to the presence of isoforms in the transcriptomes, which made it difficult to distinguish homologs from paralogs, leading to the different branching orders in the transcriptome-based trees (Cheon et al. 2020). Thus, genomic data is still recommended when available. Nonetheless, the miniprot-based strategy in transcripts could be a more accurate way compared with TransDecoder strategy in tree construction and still highly robust at the genus level, since the transcripts were obtained by blasting with closely relatives, which in some cases, would reduce the impact of contamination.

We also tested Read2Tree (Dylus et al. 2023) with the same datasets and made a comparison with VEHoP. Read2Tree only accepts marker genes from the OMA database, where only three mollusc species are currently available. We used marker genes of these abovementioned species as a reference to reconstruct phylogenetic trees with Read2Tree. Both Read2Tree and VEHoP were not able to reveal the same branching order as that of the high-quality genome dataset. The position of *Crassostrea nippona* was unstable. However, VEHoP successfully recovered the same branching order as the same as reference topology inferred from well-annotated genomes, while Read2Tree retained the branching order with low-coverage datasets. As for run time comparison, VEHoP performed much quicker with dataset less than 4G. After 4G, Read2Tree took less time than VEHoP, since it reconstructed trees directly from raw sequencing reads, and VEHoP needed to assemble the reads first before proceeding with phylogenetic reconstruction. Apart from Read2Tree, MIKE was also tested with the same datasets mentioned above. Though the total run time of MIKE was much less than both Read2Tree and VEHoP, the branching orders generated by MIKE were unstable in most datasets. *Saccostrea glomerata* grouped within *Crassostrea* in most cases (Supplementary Fig. 3), with the sole exception of the 6G dataset, where *S. glomerata* grouped with *Ostrea*. Besides, none of the branch order were the same as reference topology (Supplementary Fig. 4). Compared with Read2Tree and MIKE, VEHoP accepts all three types of input data, including proteins from well-annotated genomes, transcriptomes and DNA genomic data, as well as raw Illumina reads, which highly improved the taxon sampling in the phylogenetic analysis.

To test the universality of VEHoP across diverse taxa, we employed catfish and insect datasets for testing, comparing the results with those of Read2Tree and MIKE. In these two benchmark tests, the tree generated based on well-annotated genomes was chosen as the reference topology. VEHoP successfully reproduced the same branch orders as that of the reference topology in both cases. In the catfish datasets, all results generated by VEHoP exhibited a high degree of consistency with 100% support for all nodes. In contrast, Read2Tree and MIKE misclassified outgroup species within the ingroup catfishes. Regarding the insect datasets, VEHoP effectively recovered both Hemiptera and Thysanoptera as monophyletic groups with high bootstrap support value. But in Read2Tree, Thysanoptera was found to be paraphyletic. Moreover, in MIKE, none of the orders were recovered as monophyletic. These findings indicate that assemble-free phylogenomic methods still have certain limitations. The inconsistency was also shown in the root-to-tip distance analysis.

We applied VEHoP to resolve the evolutionary history of the deep-sea gastropod order Neomphalida, which mostly lacks high-quality genome assemblies (unlike the three benchmarking tests). The topology shown on Fig. 3 obtained by VEHoP is identical to ‘topology 1’ in a former study using mitochondrial genomes (Zhang et al. 2024), which lends further support to the hypothesis of multiple habitat transitions from non-chemosynthetic deep sea to various chemosynthetic habitats, i.e., hot vent, sunken wood, or even inactive vent, over the evolutionary history of Neomphalida (Chen et al. 2024). These results indicate that phylogenomic analyses using VEHoP are more robust than phylogenetic analyses using mitochondrial genomes and the other two published software (i.e., MIKE and Read2Tree).

We acknowledge that VEHoP currently has several limitations: 1) In some uncommon cases (not shown in this work), HmmCleaner.pl or BMGE appeared to get ‘stuck’ on a single OG, taking up to thousands of minutes on a single OG. 2) The data size imbalance of raw reads may result in unstable topology through VEHoP, such as data from organisms with extremely low read coverage (< 2X). This might also lead to the expurgation of some taxa, if the strict occupancy criteria (e.g. >80%) is applied. Therefore, adjustment of occupancy and length thresholds are recommended when processing low-coverage sequenced samples. 3) So far, VEHoP is only compiled for use in the Linux system. We are improving the pipeline to make it more widely accessible (e.g., on Windows system).

With VEHoP, users can define a highly customizable dataset for reference, and it can be a concatenation of high-quality genomes of related species, not limited by an online orthology database, which might result in much more homologs for ortholog inference. The ortholog inference procedure used in VEHoP has been shown to work well in metazoan (Kocot et al. 2019; Sun et al. 2020; Sun et al. 2021) and bacterial (Li et al. 2023) datasets. With VEHoP, every ortholog that passes the filtering steps is kept, and the user can determine which ones to eliminate based on other criteria if desired, after the process has been completed. In the output folder, the orthologs, concatenated matrix, as well as related partition file will be available for further deep-phylogeny analyses if necessary. Overall, VEHoP shows many advantages, including fast, accurate, and user-friendly. Importantly, VEHoP makes it possible to utilize and combine genomic DNA and transcriptome data widely available in SRAs. We foresee that a wide application of VEHoP would alleviate the problem of low taxon sampling in the phylogenetic analysis of many groups of organisms.

## Author Contributions

JS and YL conceived the project. YL coded the pipeline. CC collected the samples. YL and XL carried out the phylogenetic analyses (i.e., draft genome assembly, benchmarks, reanalysis of public data) and manuscript preparation. All authors contributed to the revision of the manuscript.

## Acknowledgements

This research was financially supported by the National Key Research and Development Program of China (2024YFC2816100), Science and Technology Innovation Project of Laoshan Laboratory (LSKJ202203104), Natural Science Foundation of Shandong Province (ZR2023JQ014), Fundamental Research Funds for the Central Universities (202172002 and 202241002), and the Young Taishan Scholars Program of Shandong Province (tsqn202103036). The *Neomphalus fretterae* specimen used herein was collected during R/V *Falkor (too)* cruise FKt231024 (Project Zombie: Bringing dead vents to life – Ultra fine-scale seafloor mapping”) funded by the Schmidt Ocean Institute. We thank the captain and crew of R/V *Falkor (too)* as well as the ROV *SuBastian* team for their immense support of our science. John W. Jamieson (Memorial University of Newfoundland), the chief scientist of cruise FKt231024, is gratefully acknowledged for his diligent execution of the research cruise.

## Data Availability

The raw reads from the newly sequenced Neomphalida are deposited in NCBI BioProject (accession number: PRJNA1129887). All the raw inputs (draft genomes, transcripts, and proteins) used, and matrixes generated in this work are available at Figshare (https://doi.org/10.6084/m9.figshare.26370955.v1 including oyster dataset and https://doi.org/10.6084/m9.figshare.28189616.v1 for fish and insect datasets.). For further enquiries on how to use the VEHoP pipeline, please feel free to contact the corresponding authors.

## Code Availability

The package of VEHoP is available at https://github.com/ylify/VEHoP/.

---

**Supplementary Fig. 1.** Ostreida phylogeny by VEHoP of full-size dataset and reduced datasets, including 1G, 2G, 4G, 6G and 8G.

**Supplementary Fig. 2.** Ostreida phylogeny by VEHoP of subsampled *Crassostrea hongkongensis* data size test.

**Supplementary Fig. 3.** Ostreida phylogeny by ReadTree of full-size dataset and reduced datasets, including 1G, 2G, 4G, 6G and 8G.

**Supplementary Fig. 4.** Ostreida phylogeny by MIKE of full-size dataset and reduced datasets, including 1G, 2G, 4G, 6G and 8G.

**Supplementary Fig. 5.** Tree topologies of Fish and Insect datasets.

**Supplementary Fig. 6.** Mitochondrial genome-based phylogeny of Neomphalida.

**Supplementary Fig. 7.** Neomphalida phylogeny based on NGS data, including VEHoP (multiple models), MIKE and Read2Tree.

**Supplementary Fig. 8.** Occupancy of matrix generated by VEHoP.

