## Supplementary figures and images for "Reliable inference of phylogenomic relationship via assembly-based strategy accommodating raw reads and proteins"

### Supplementary Fig. 1

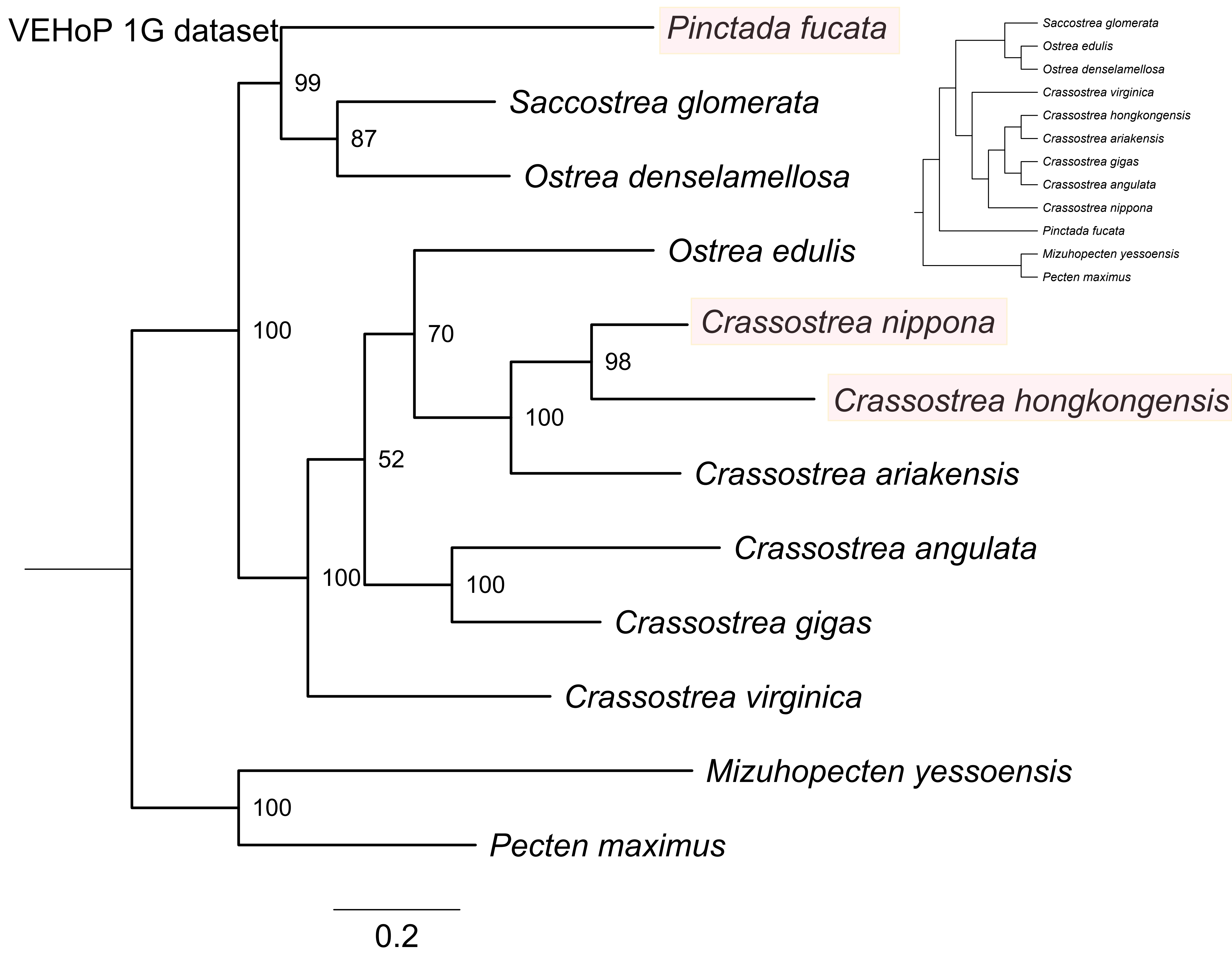

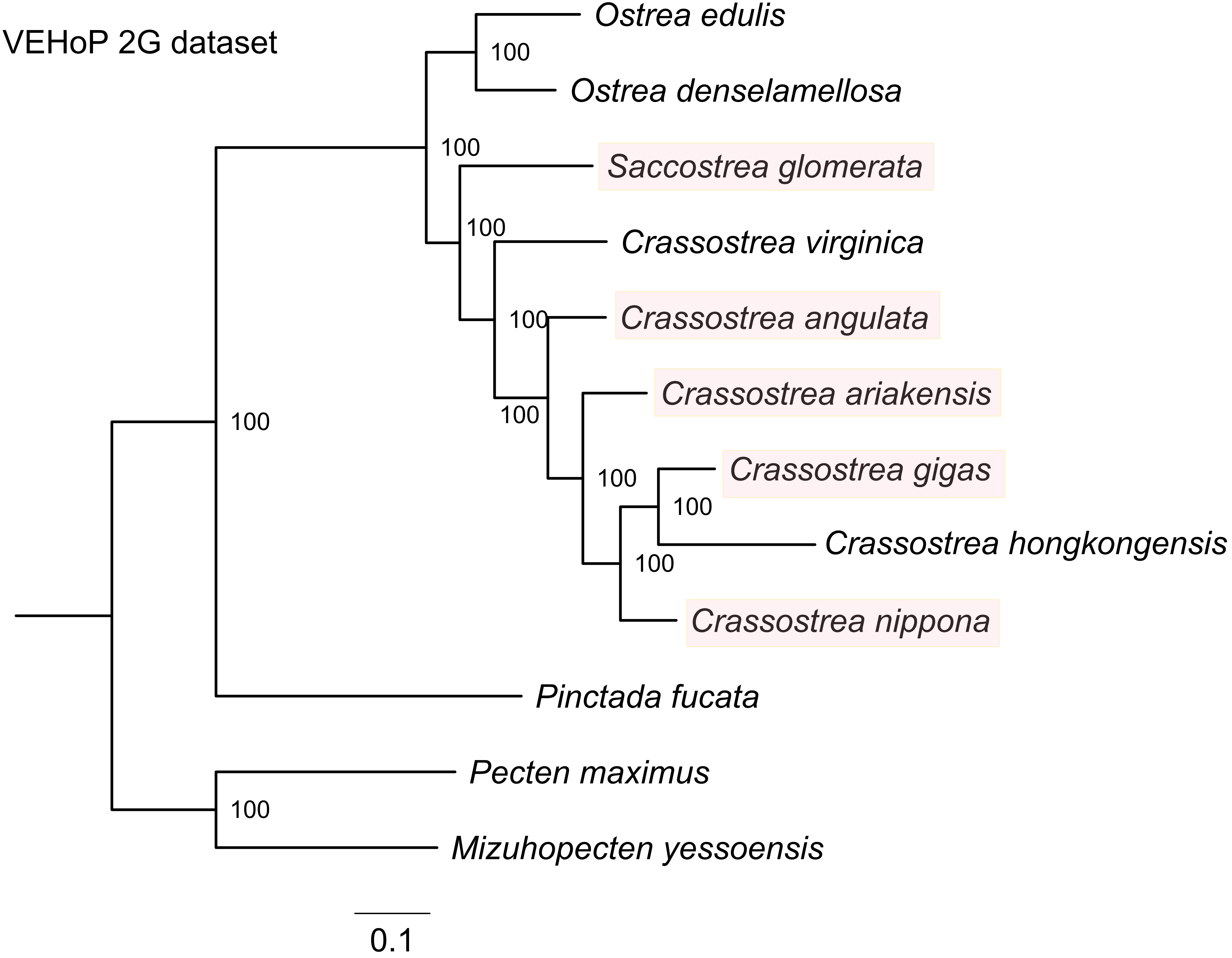

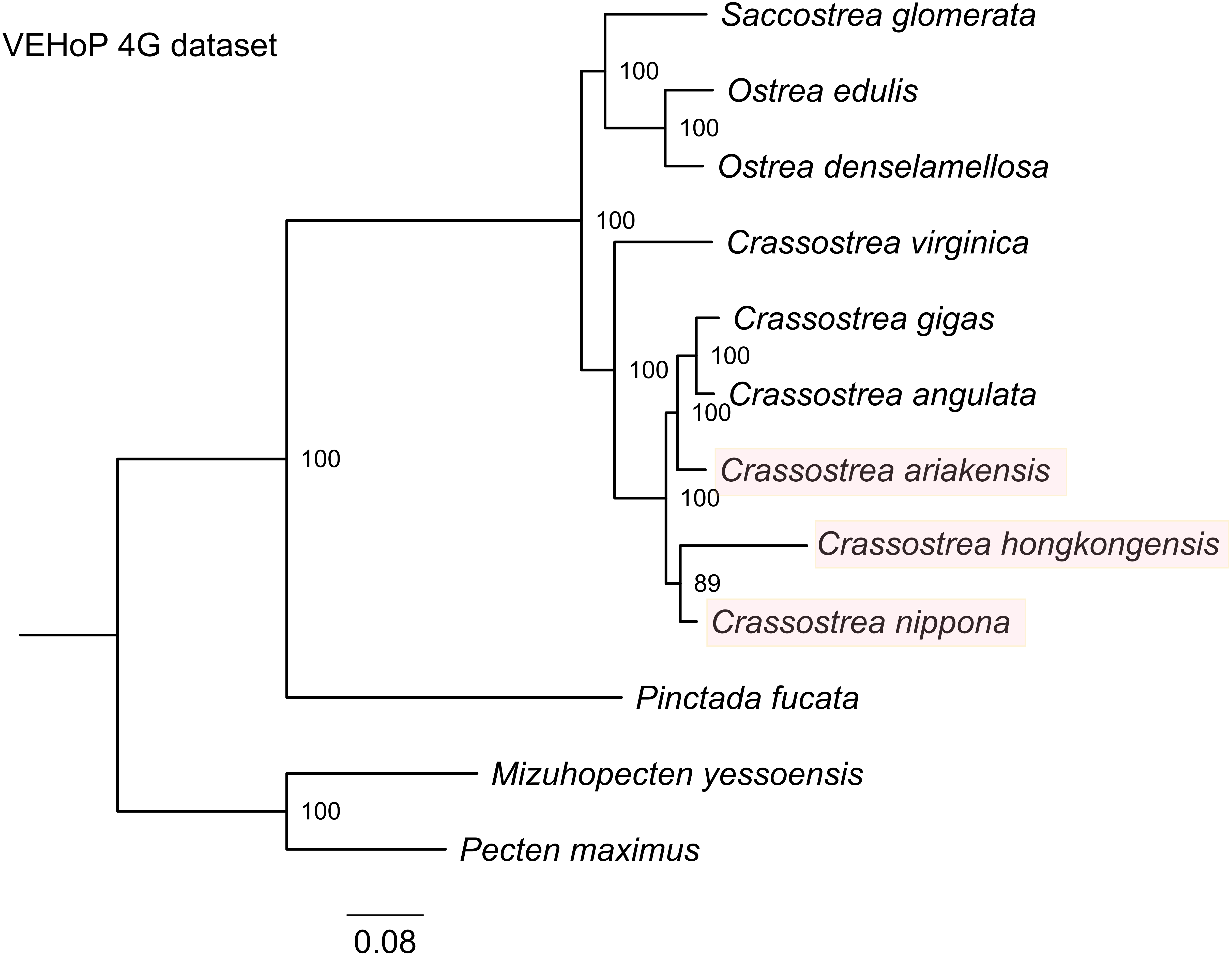

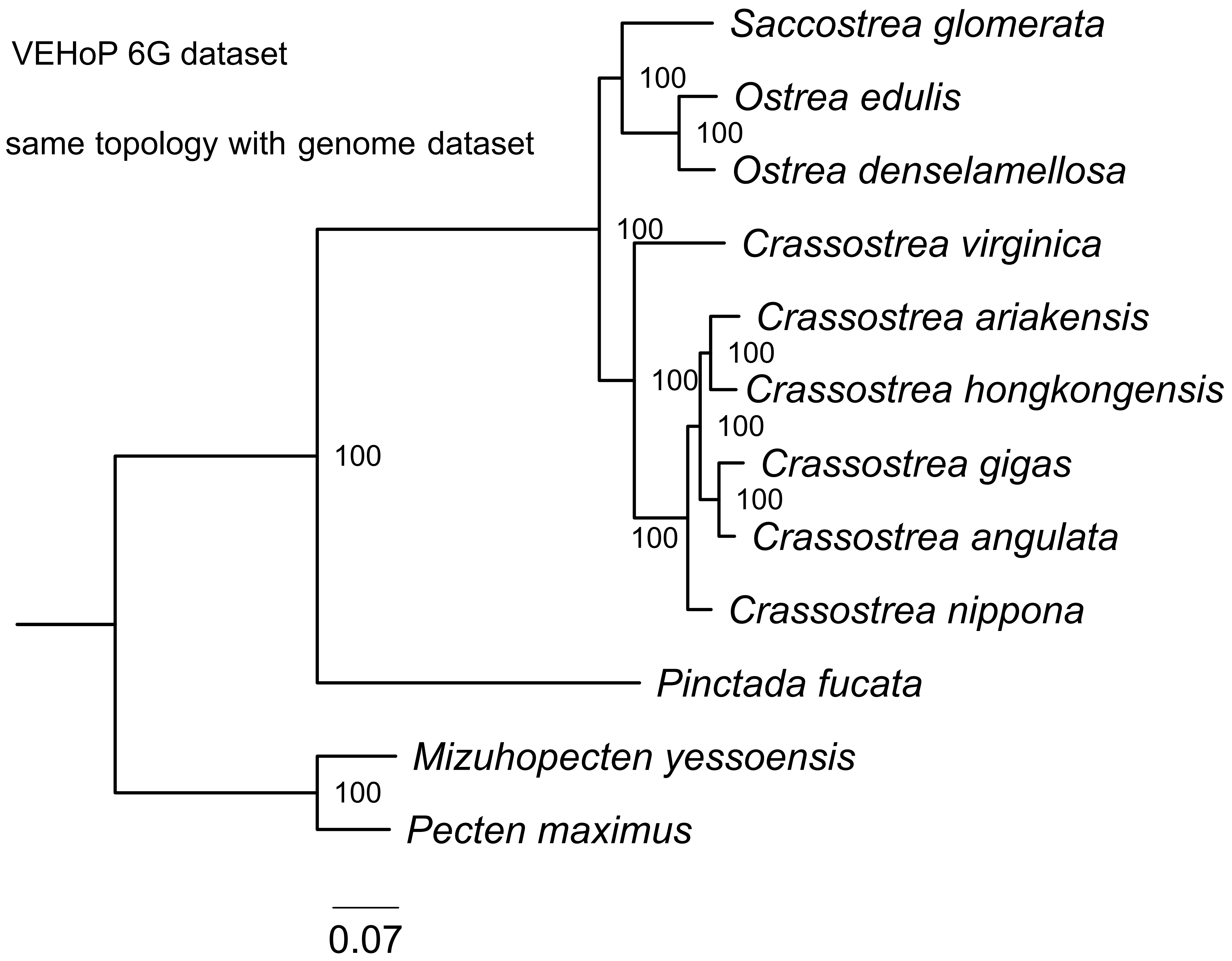

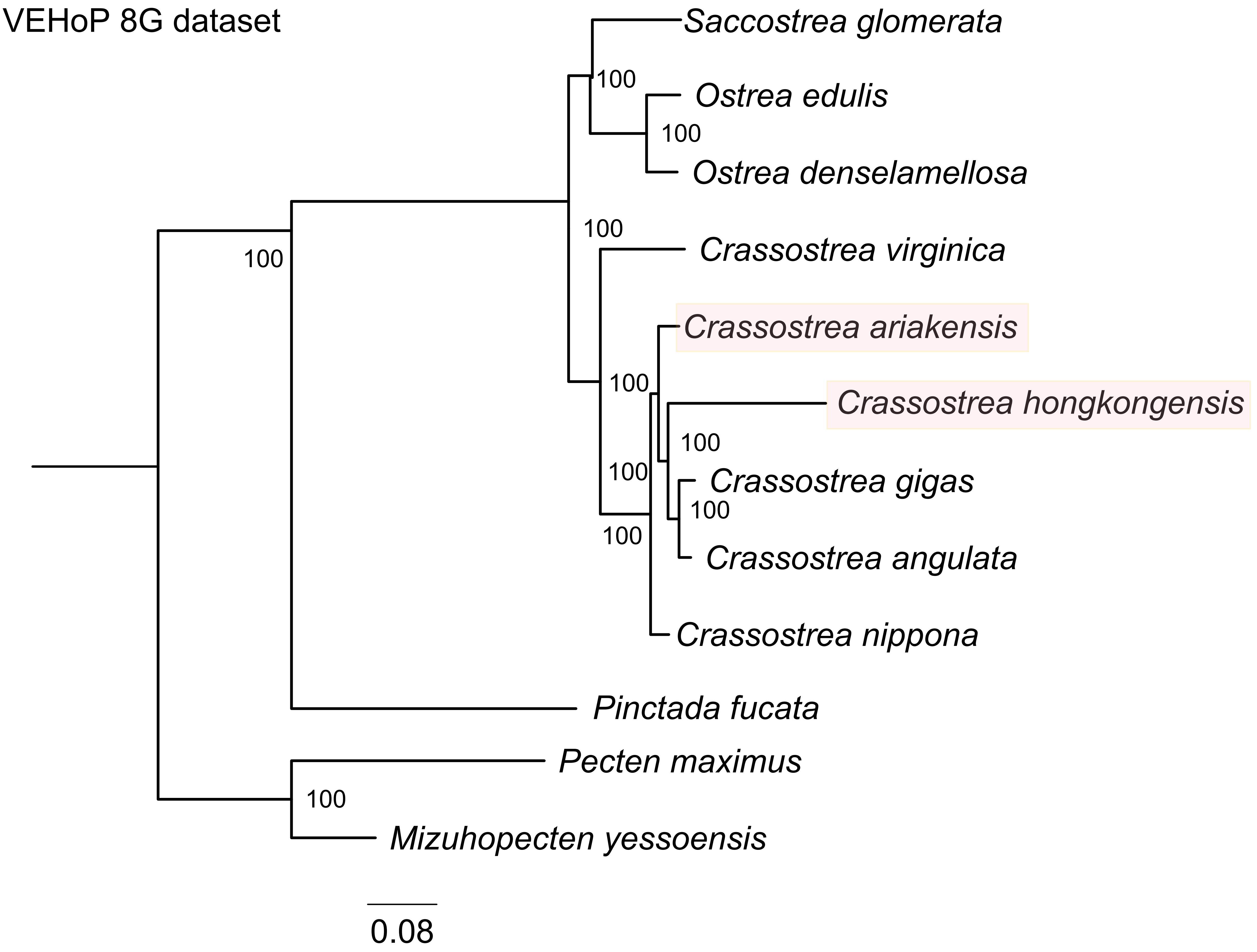

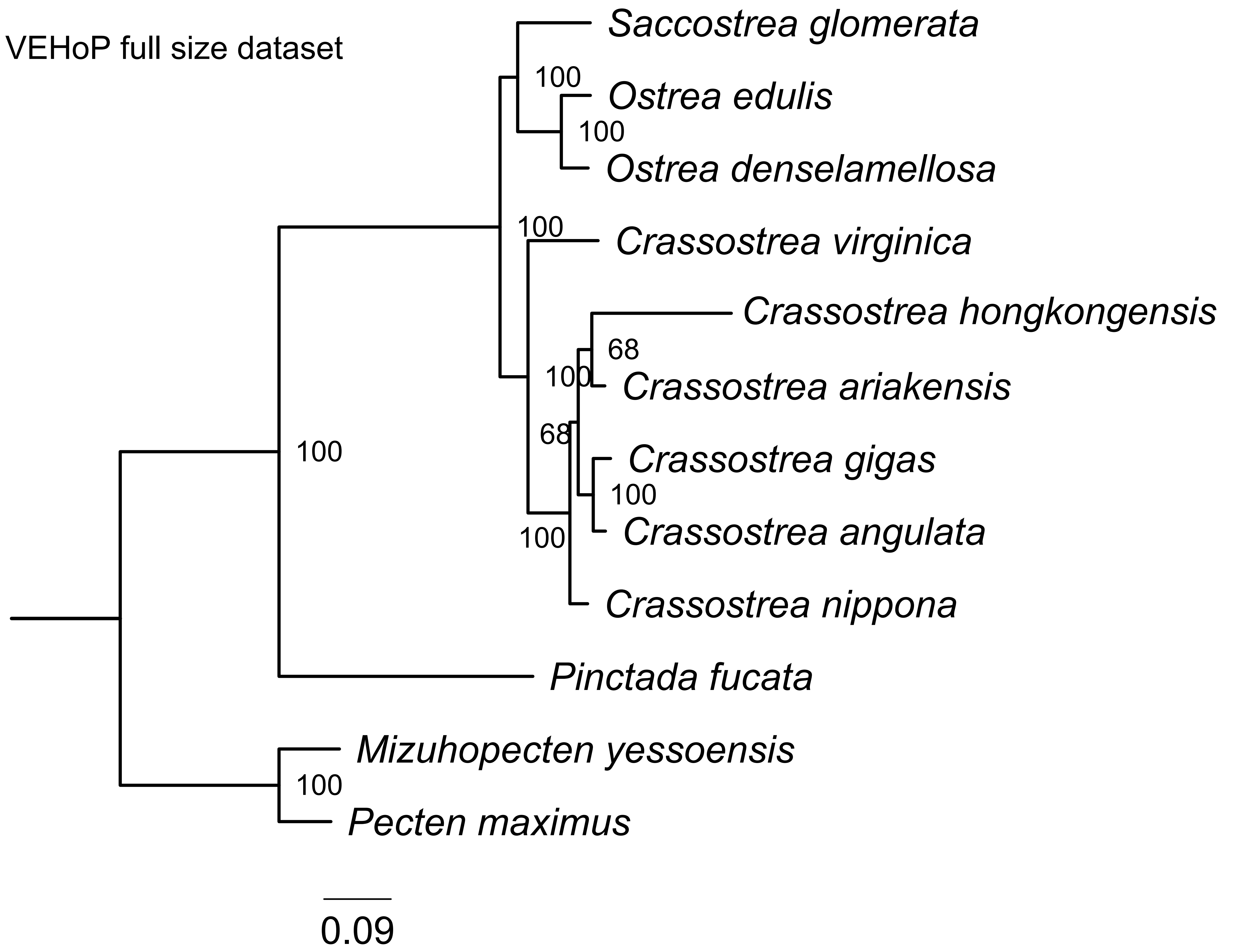

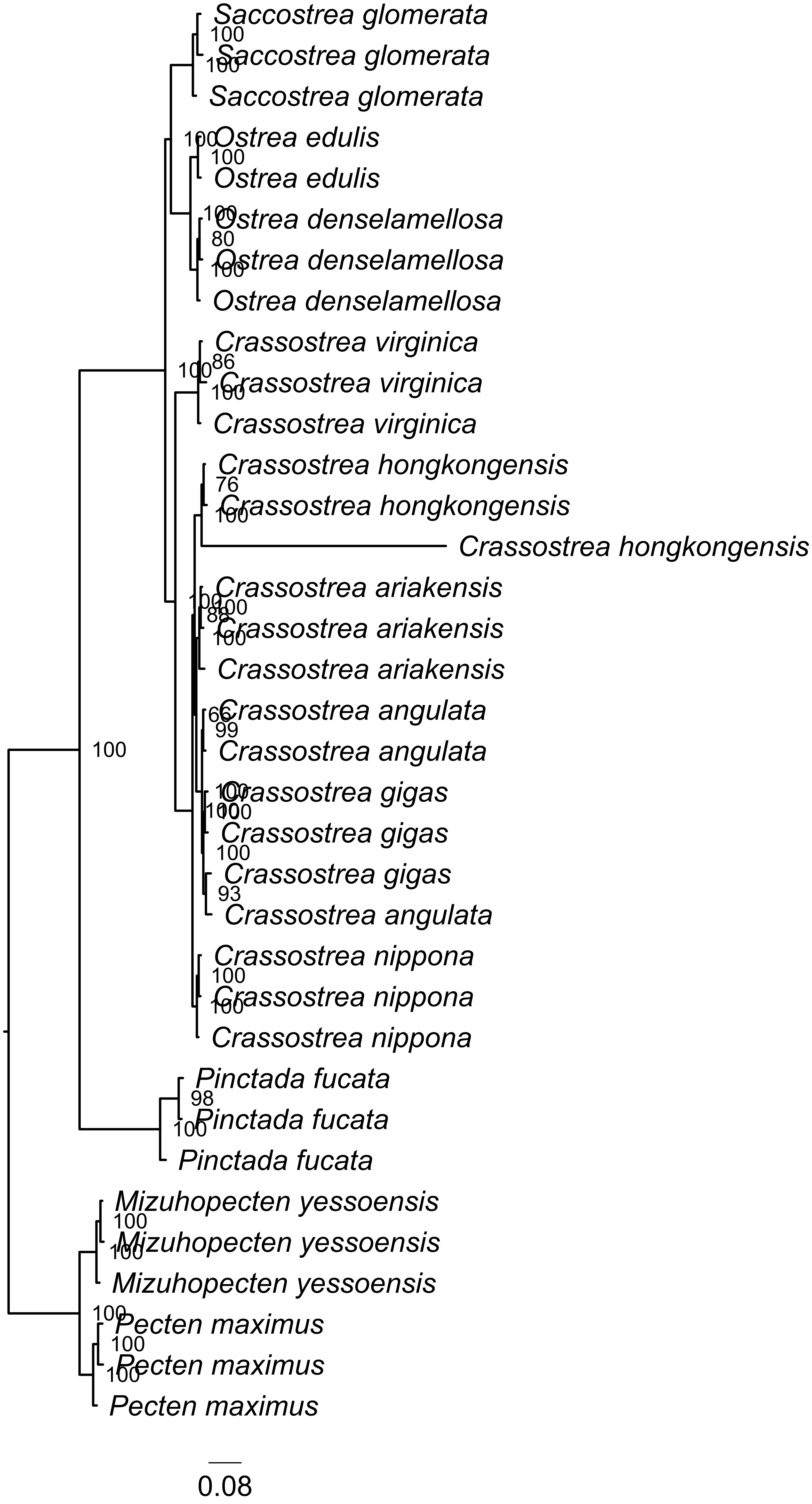

### Supplementary Fig. 2

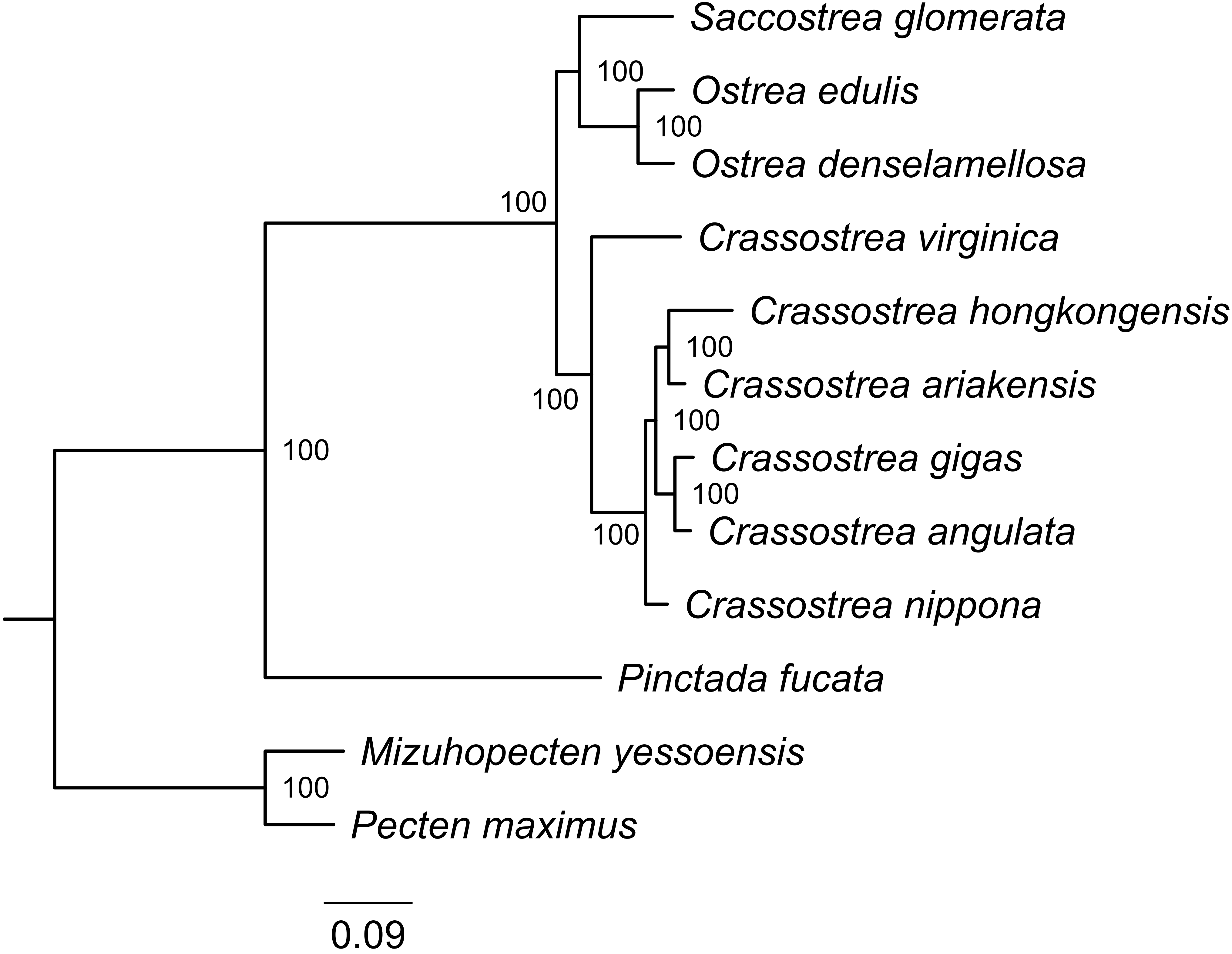

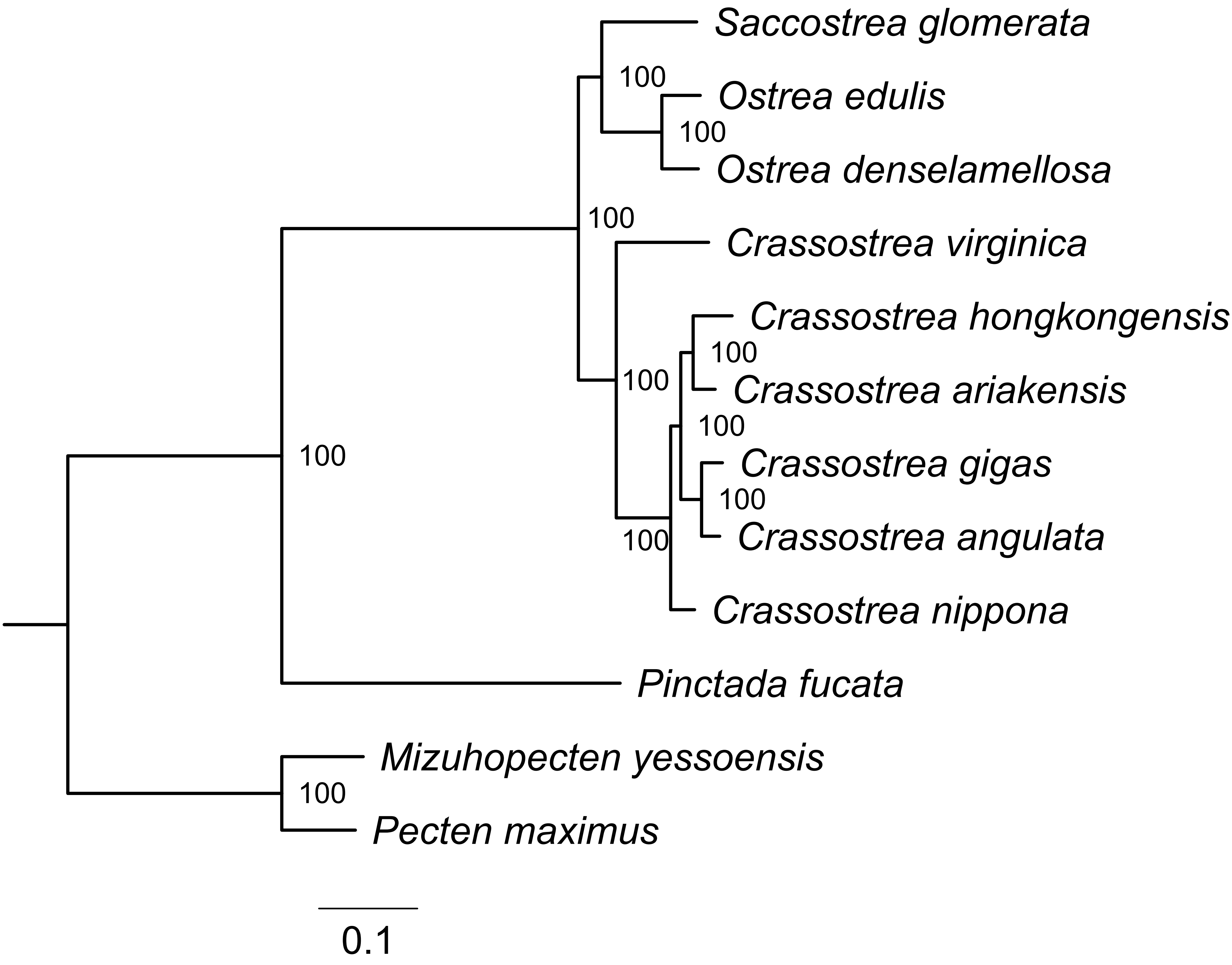

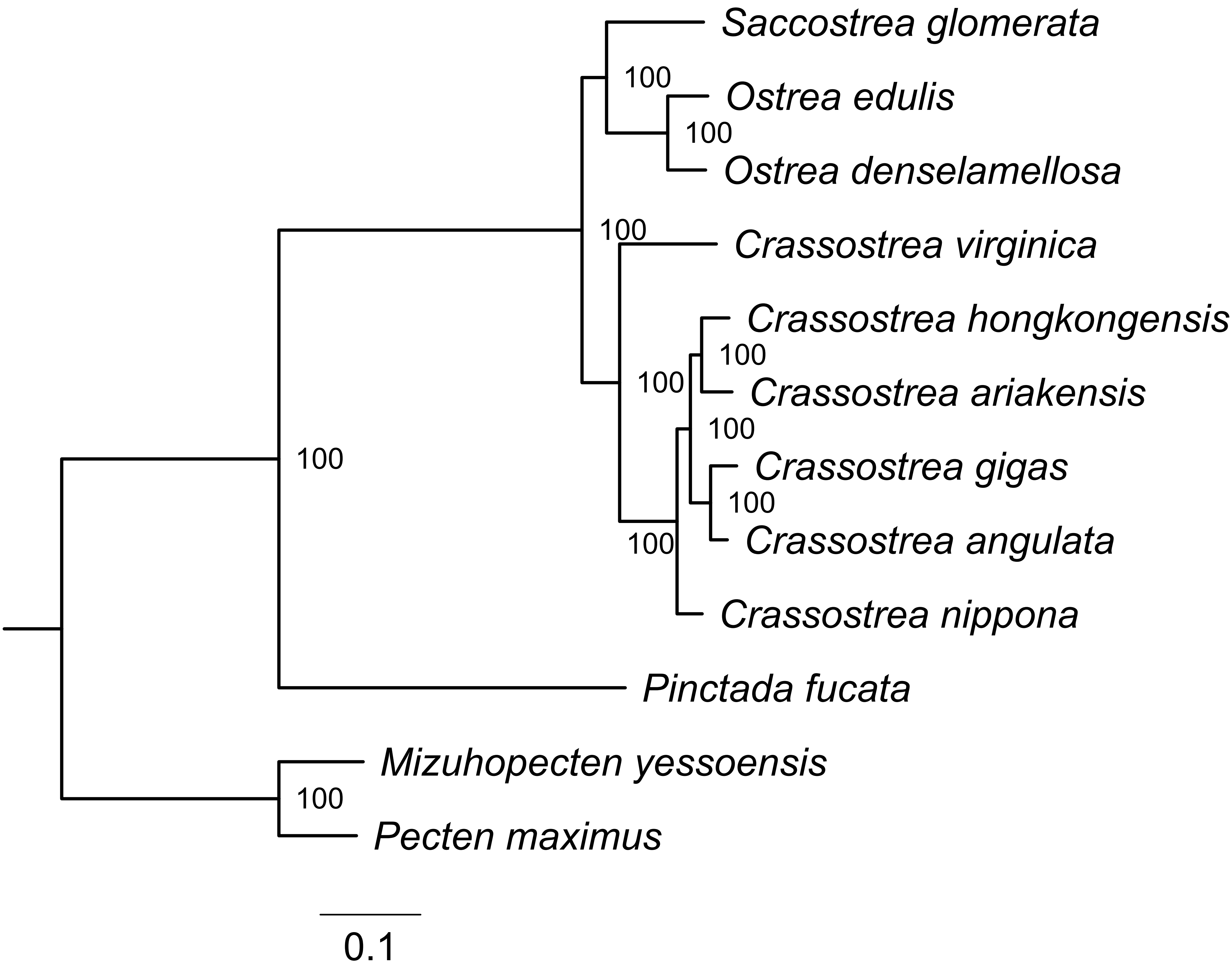

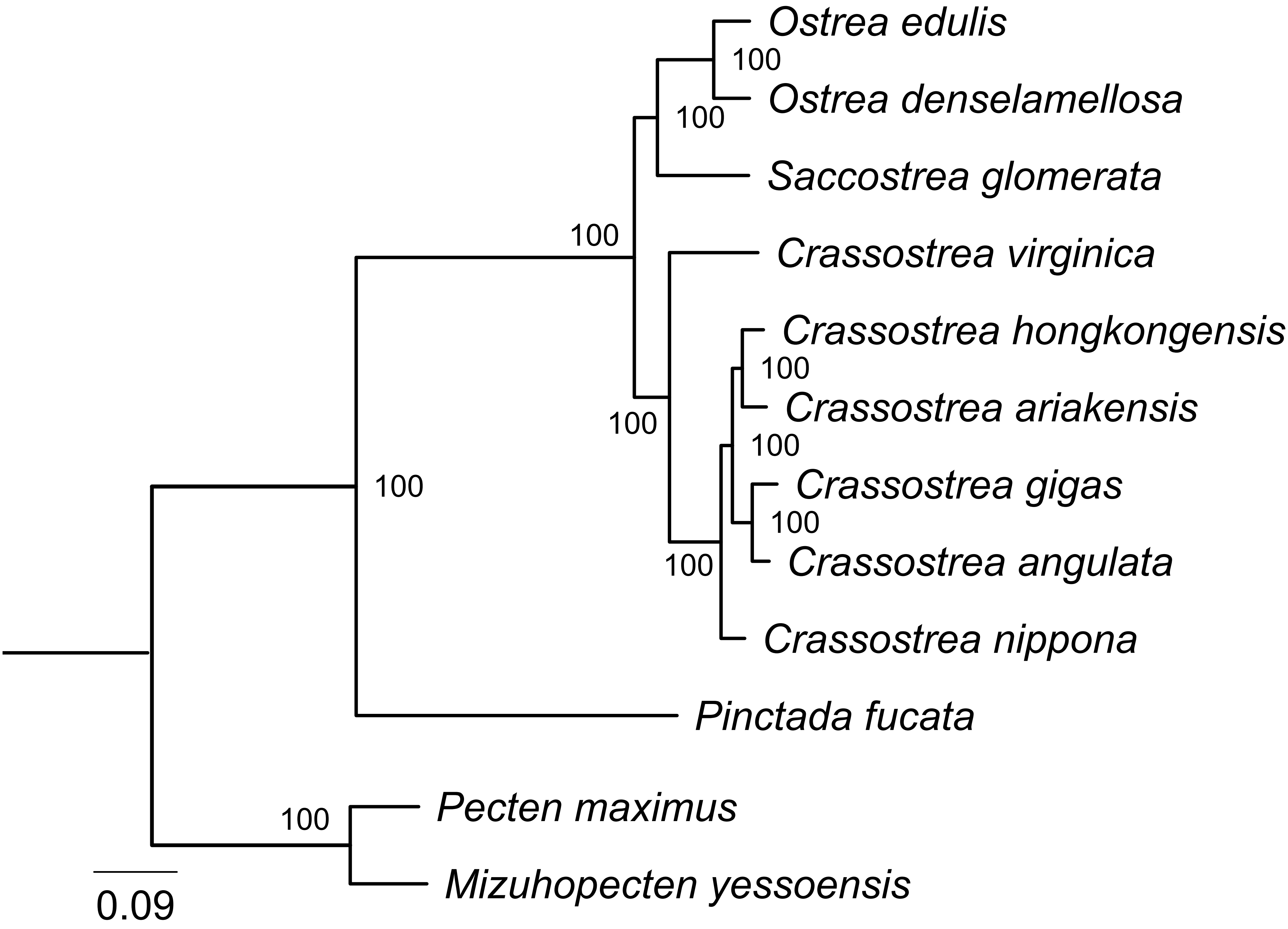

### Supplementary Fig. 3

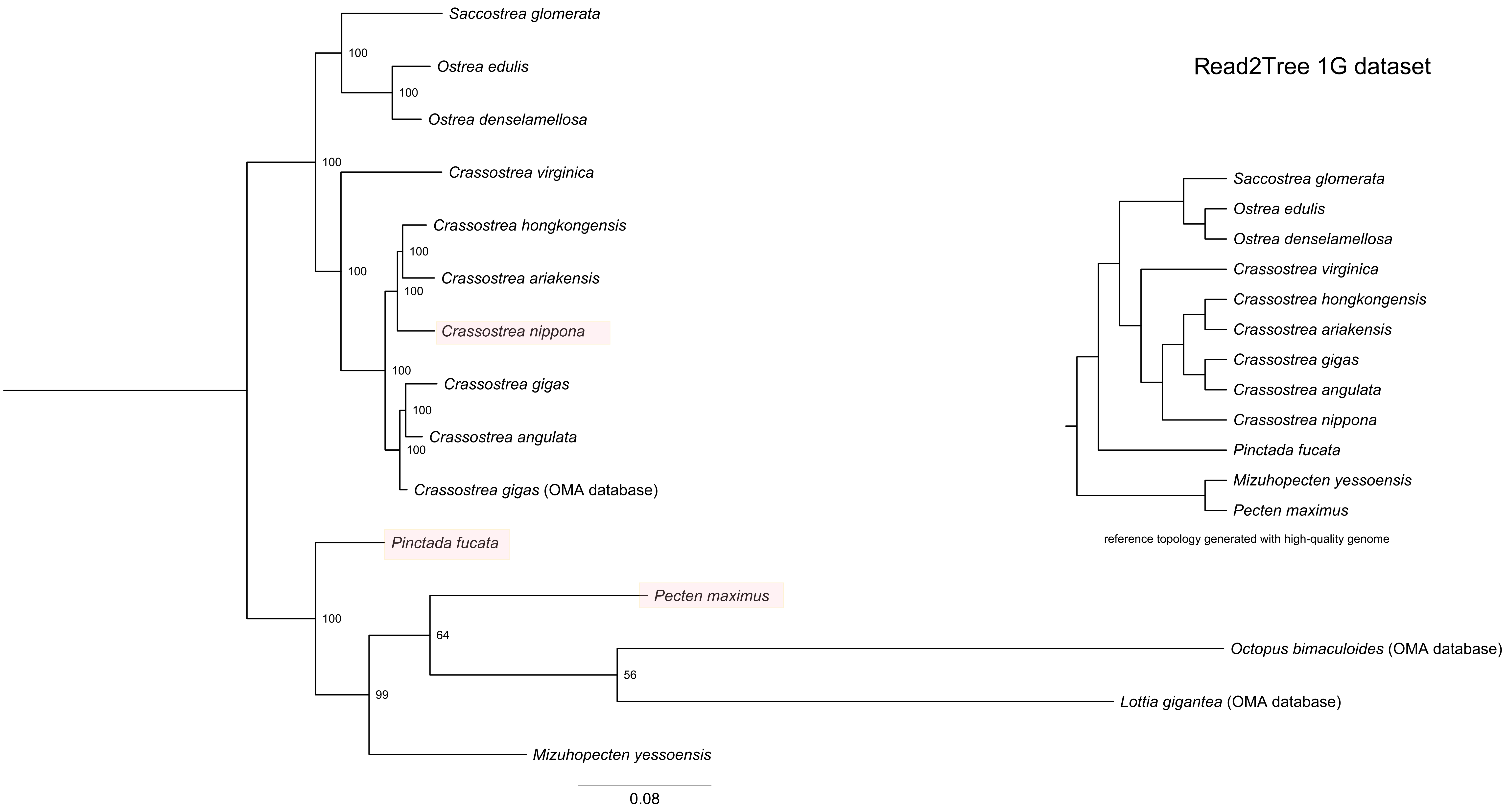

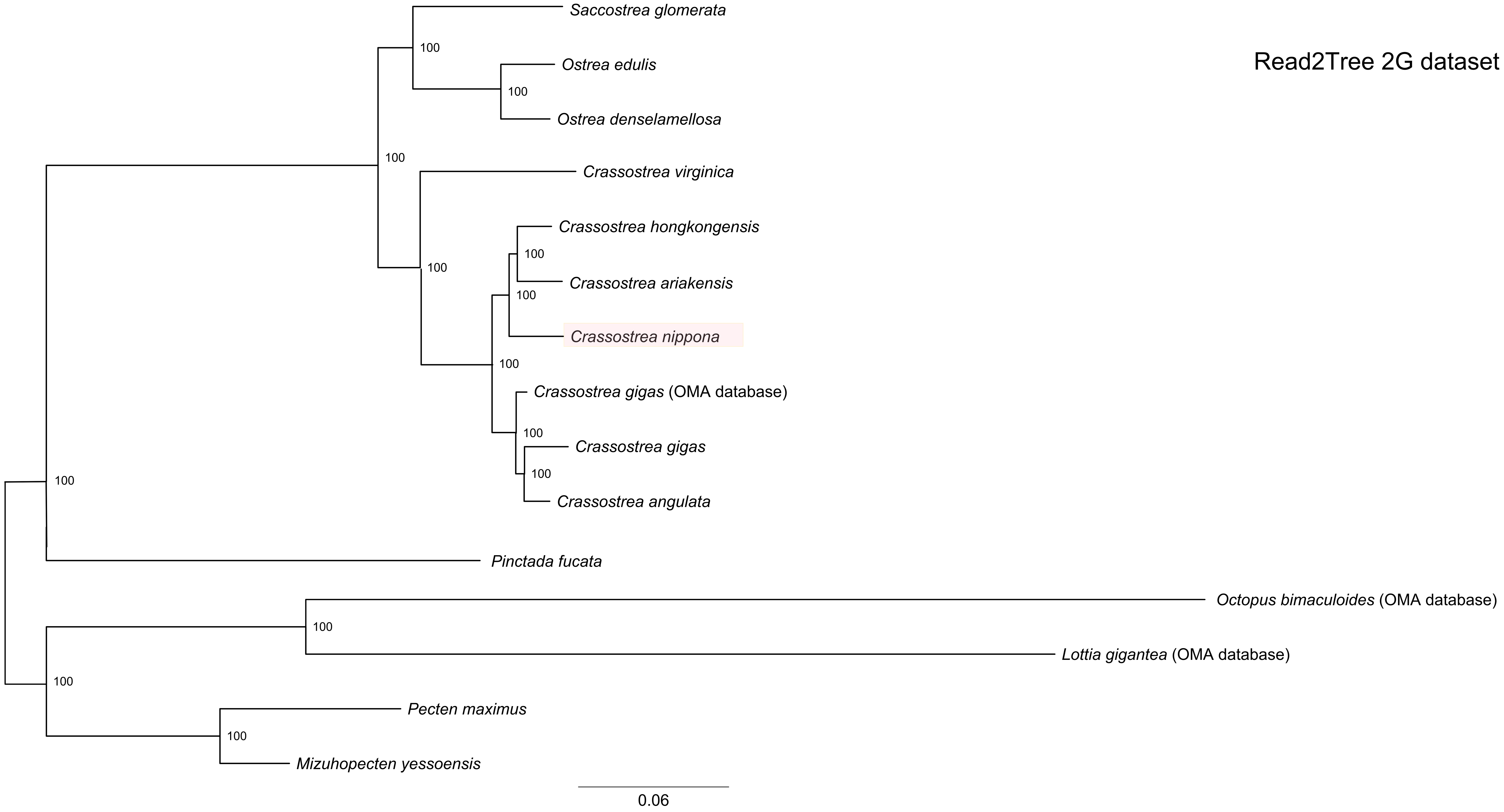

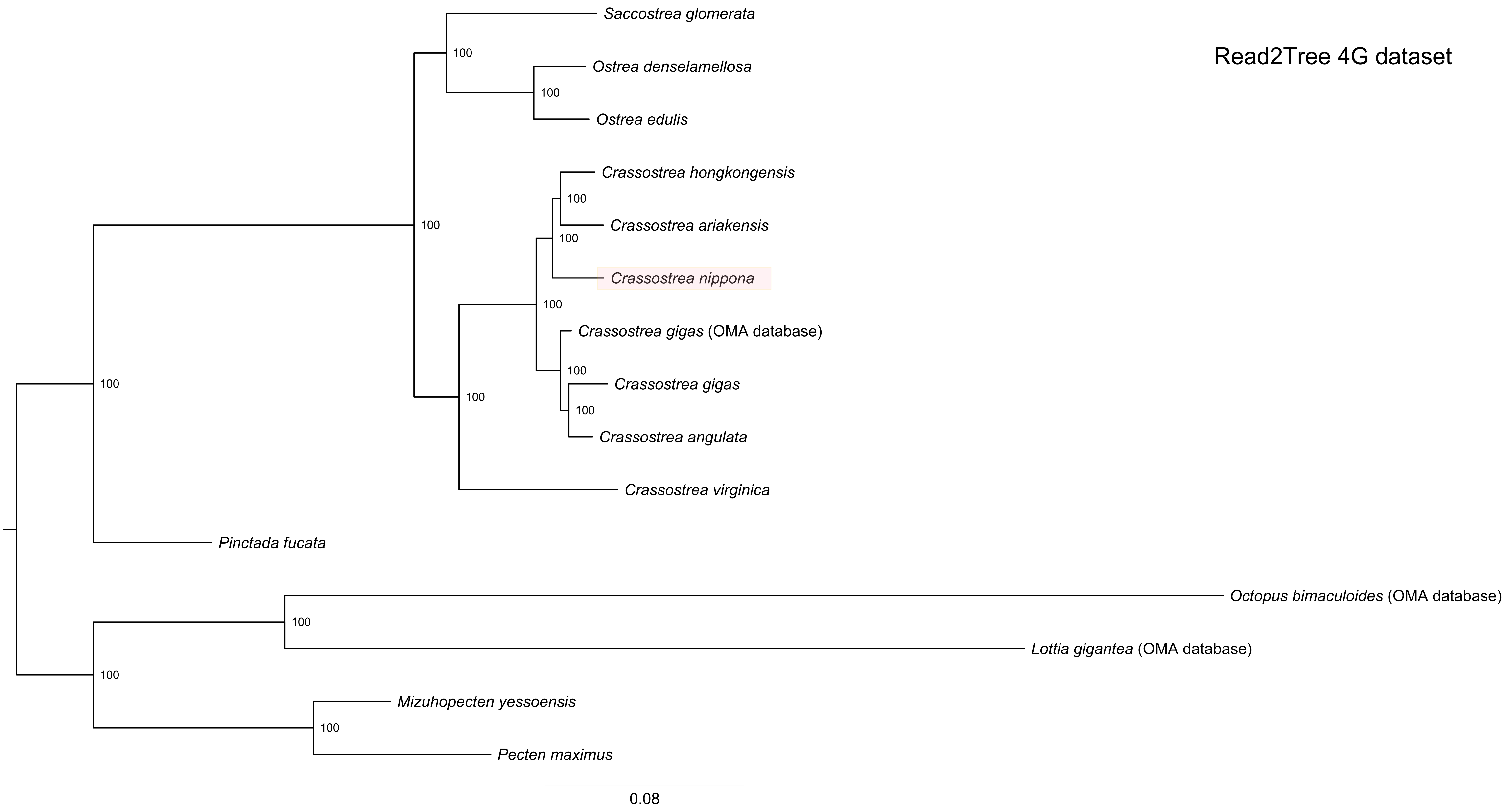

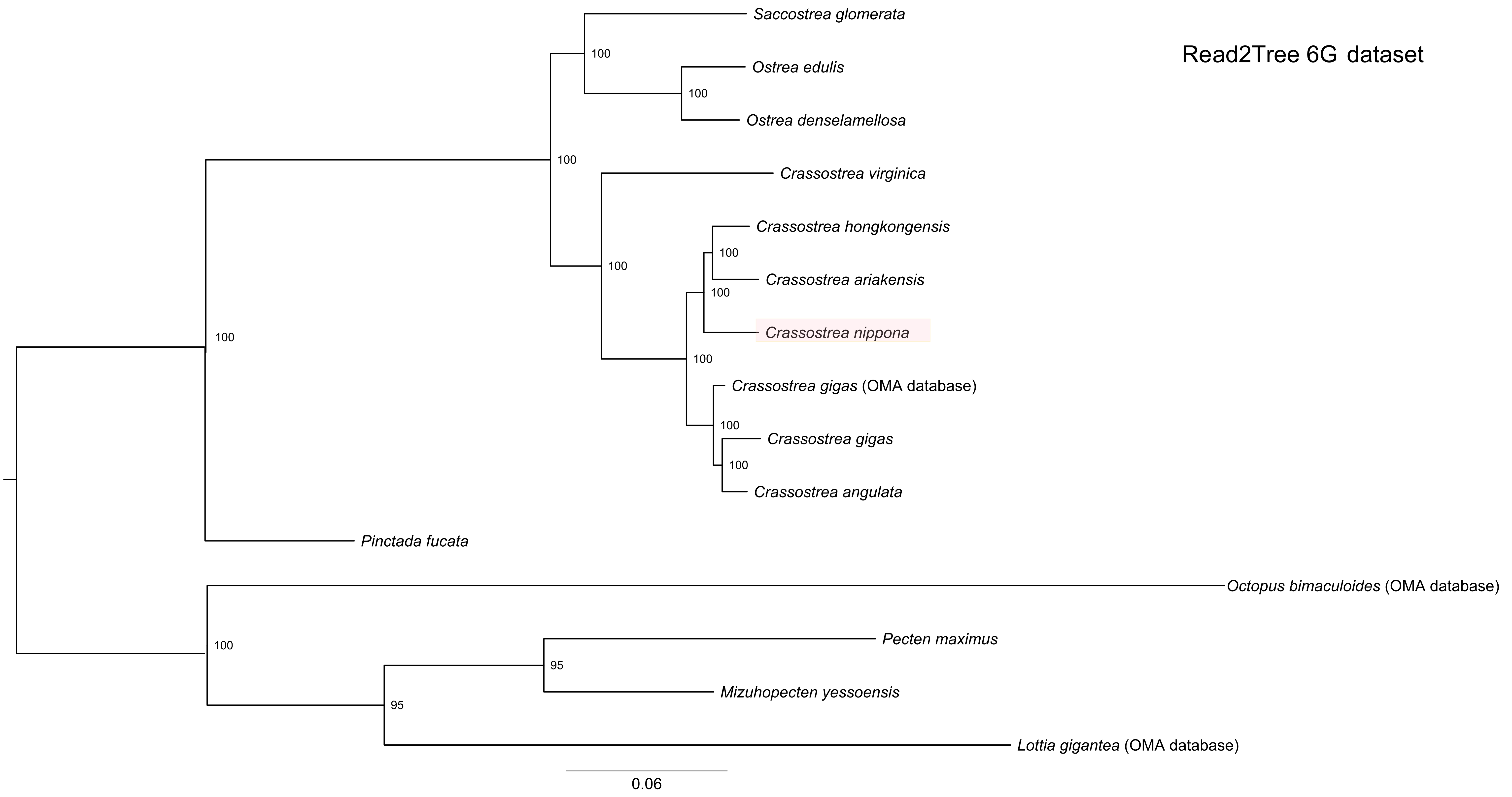

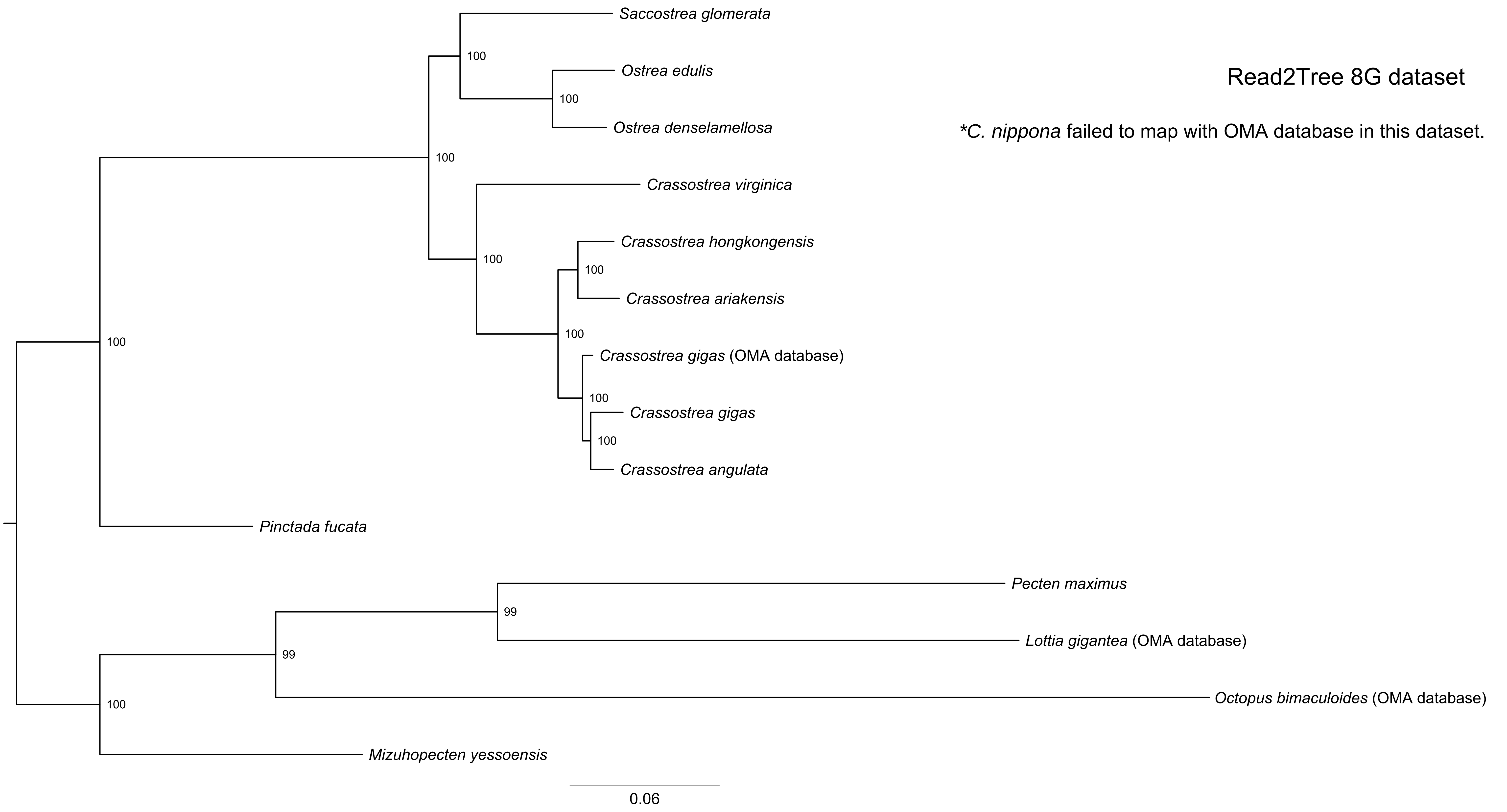

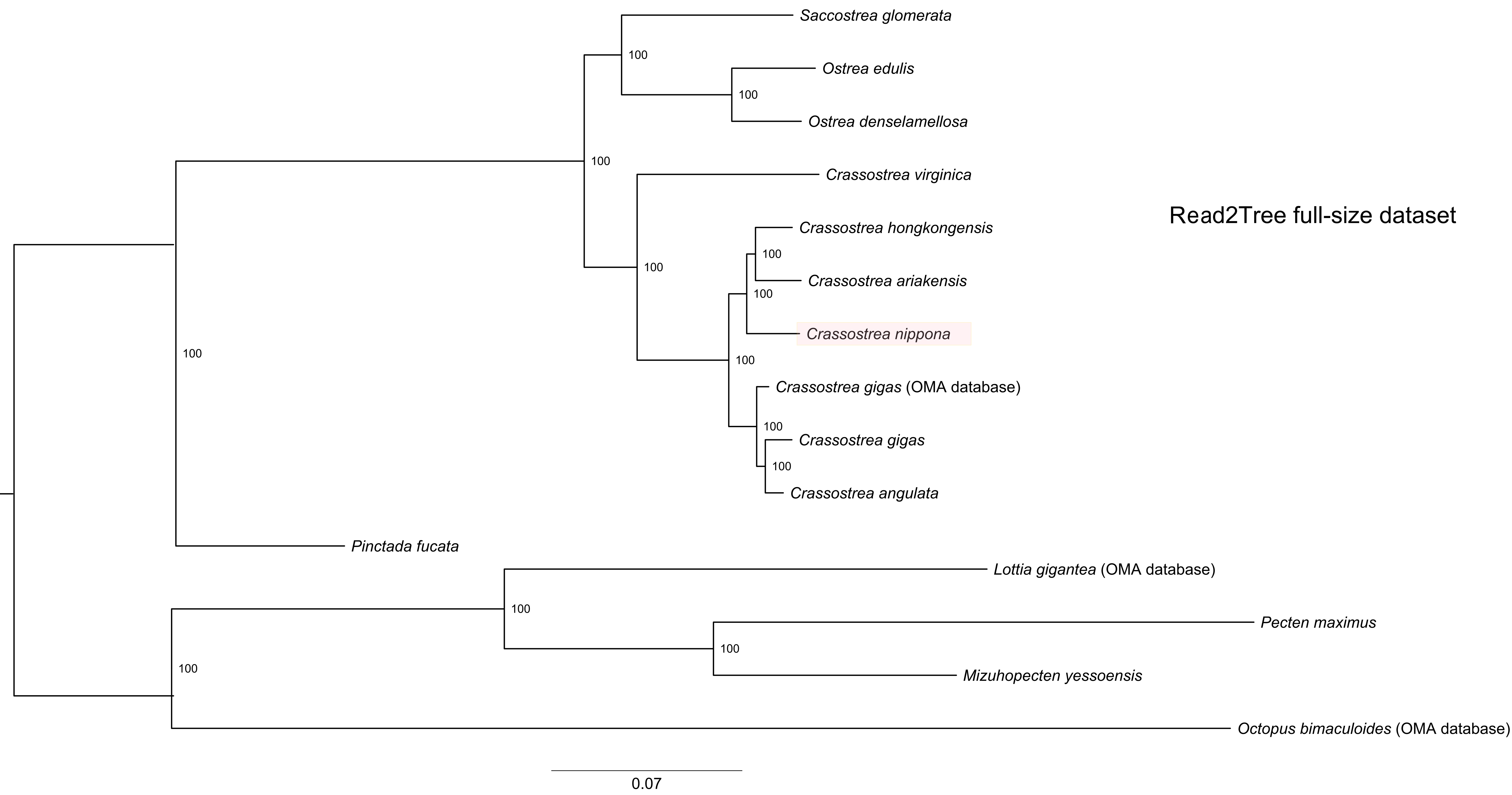

### Supplementary Fig. 4

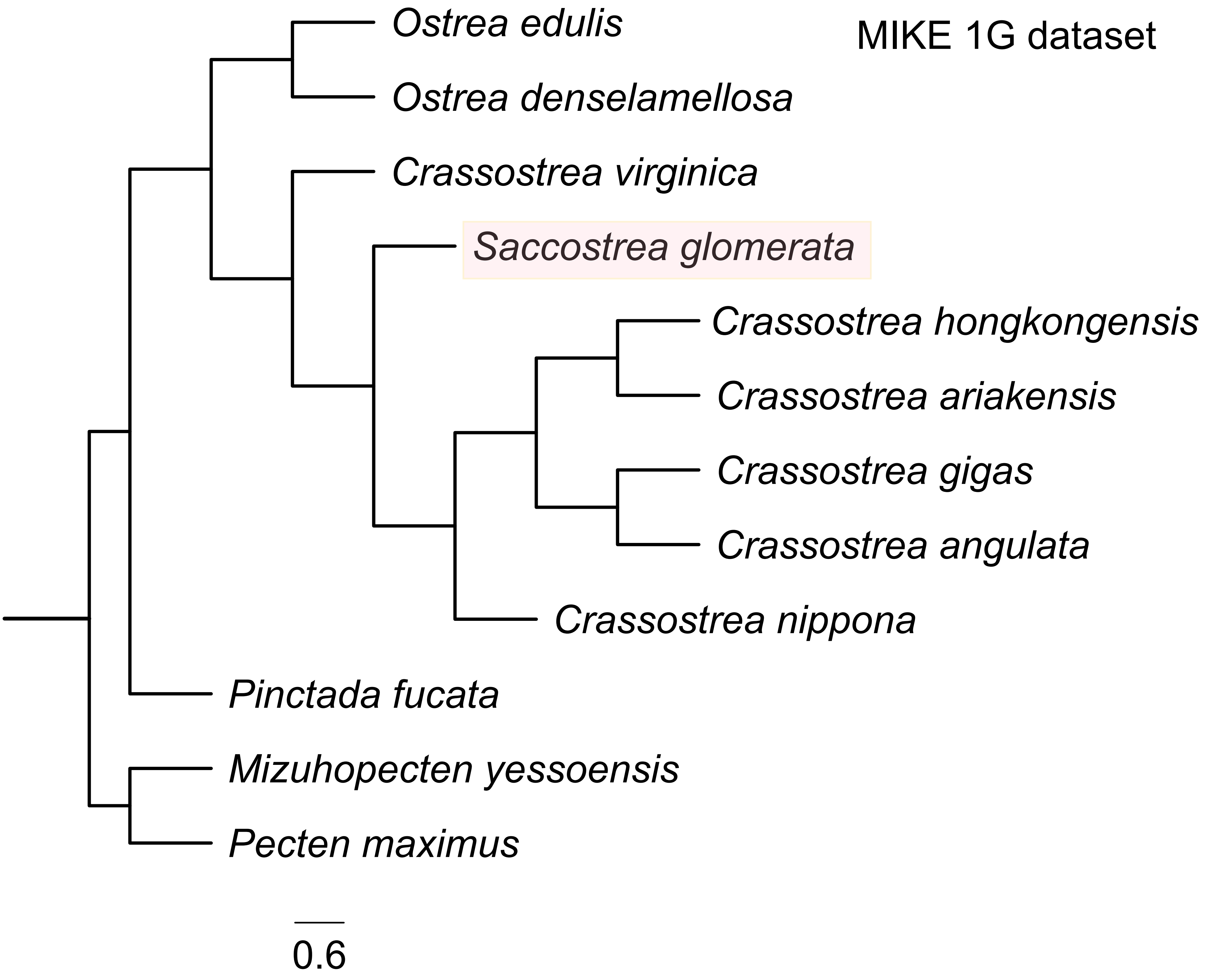

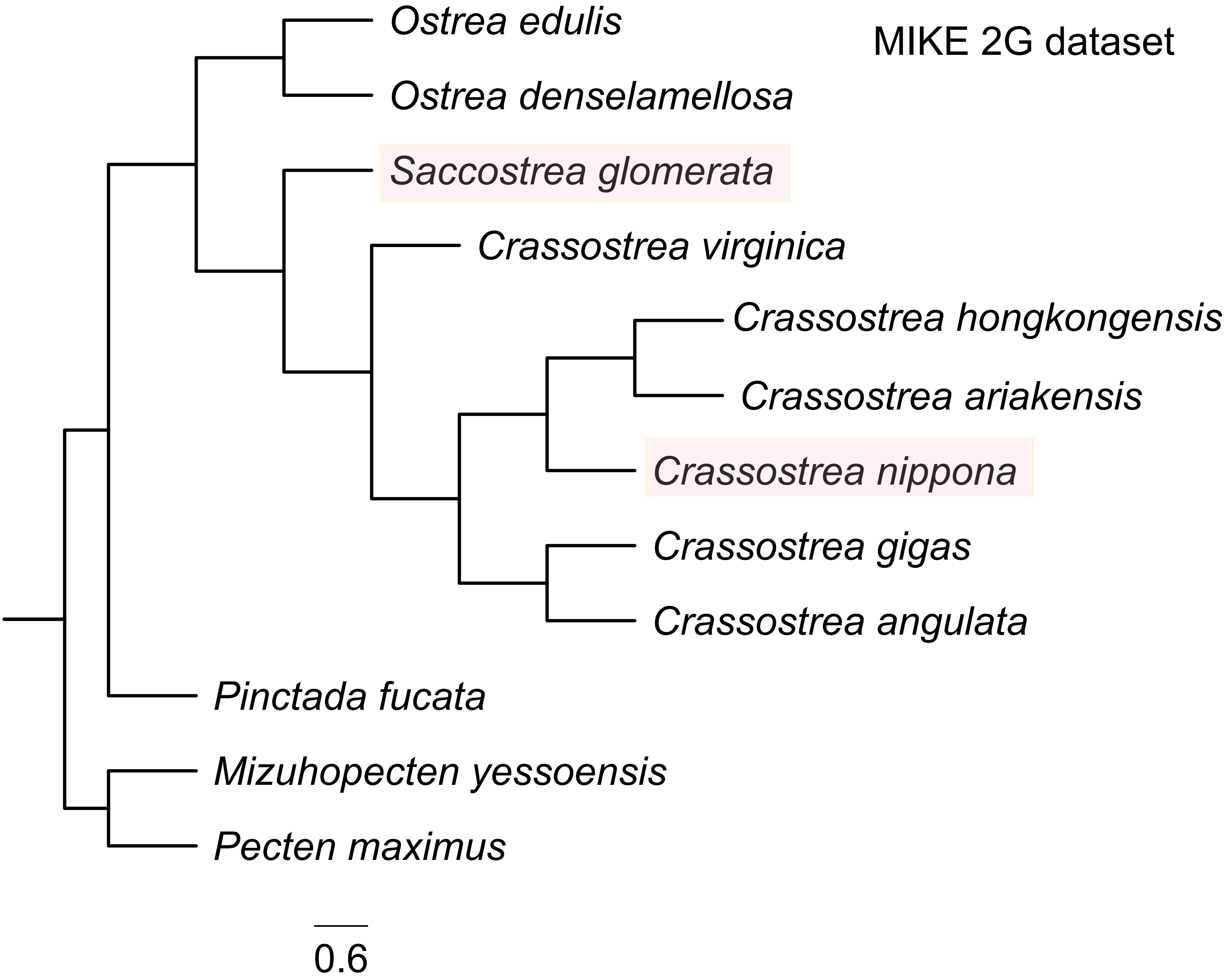

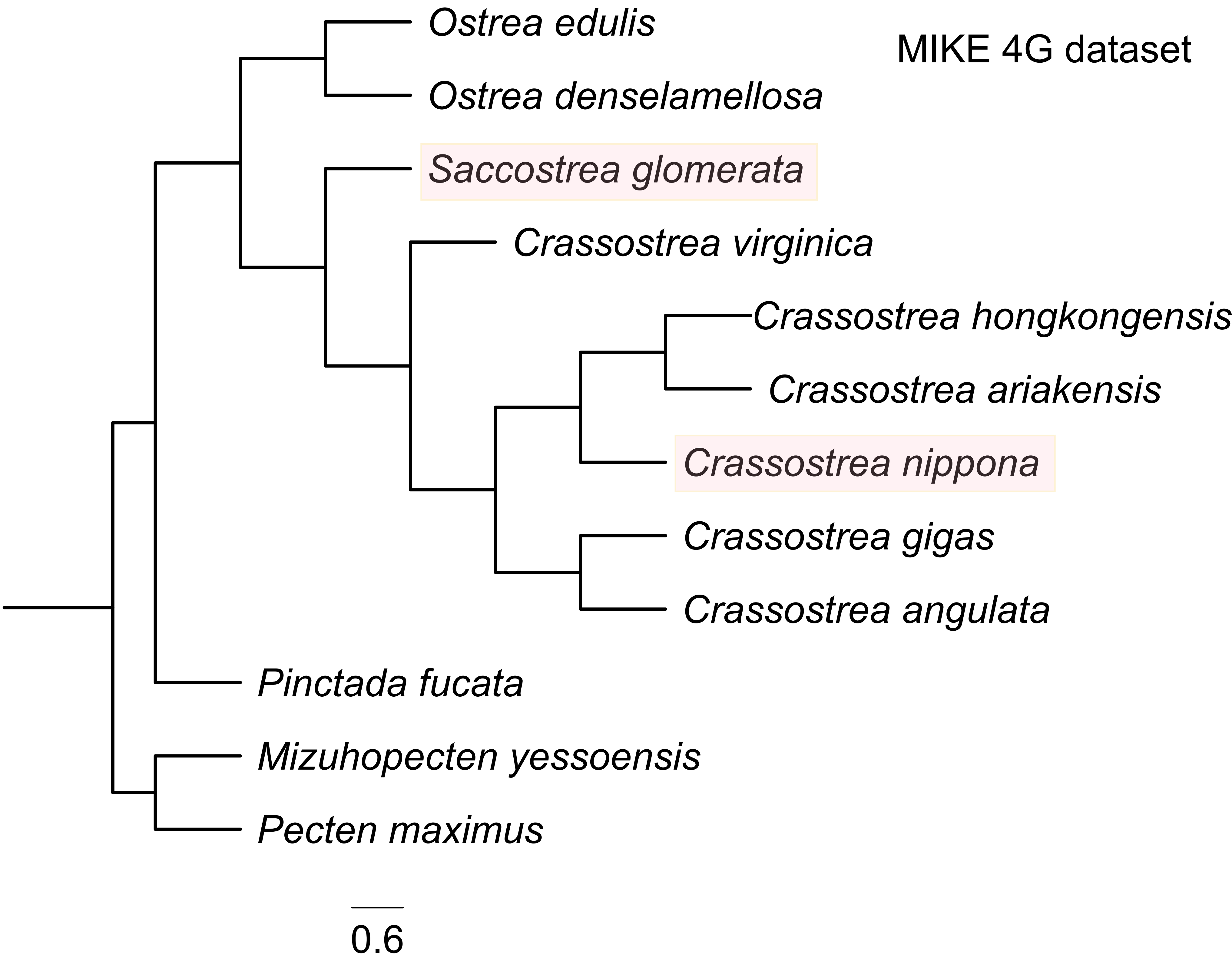

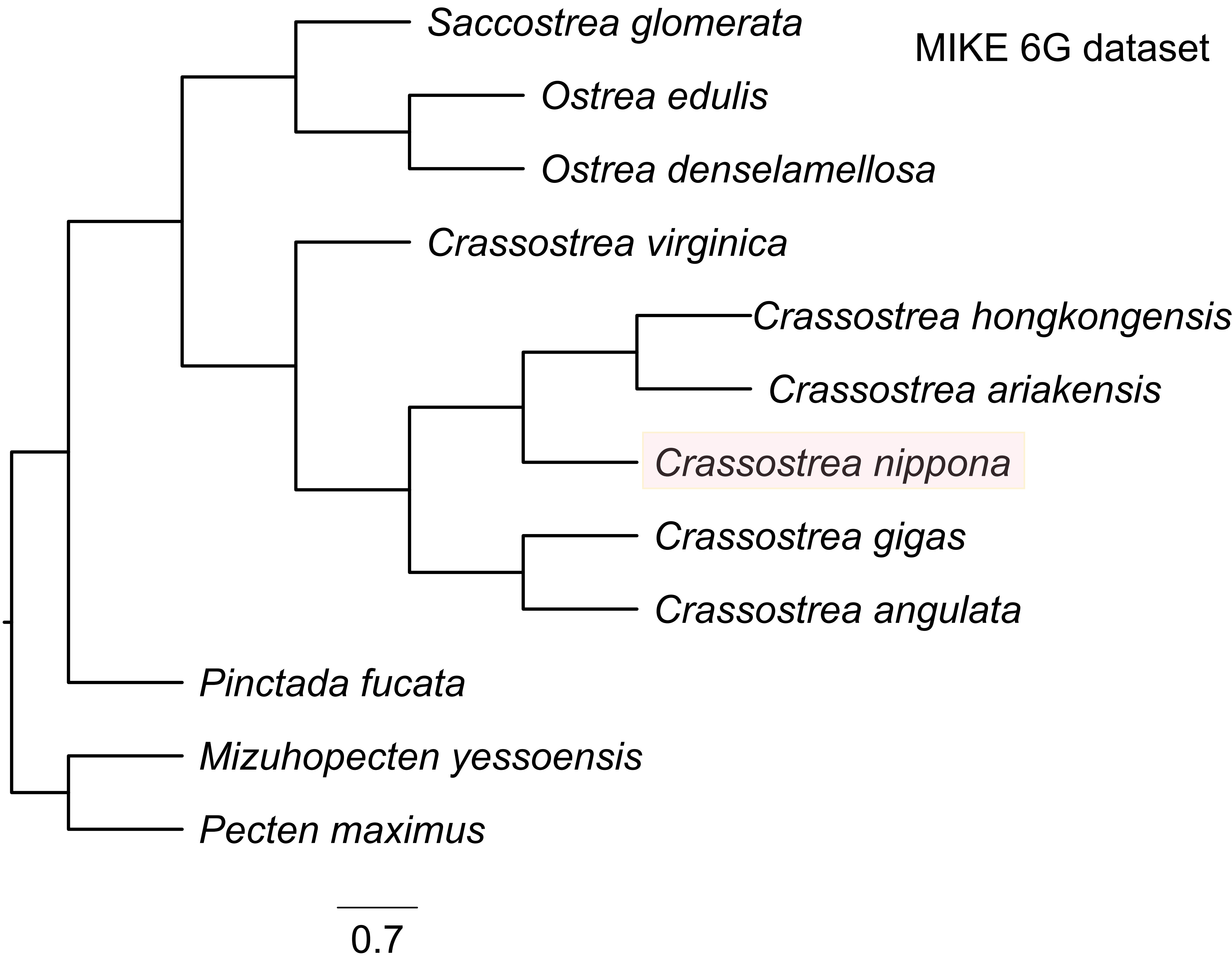

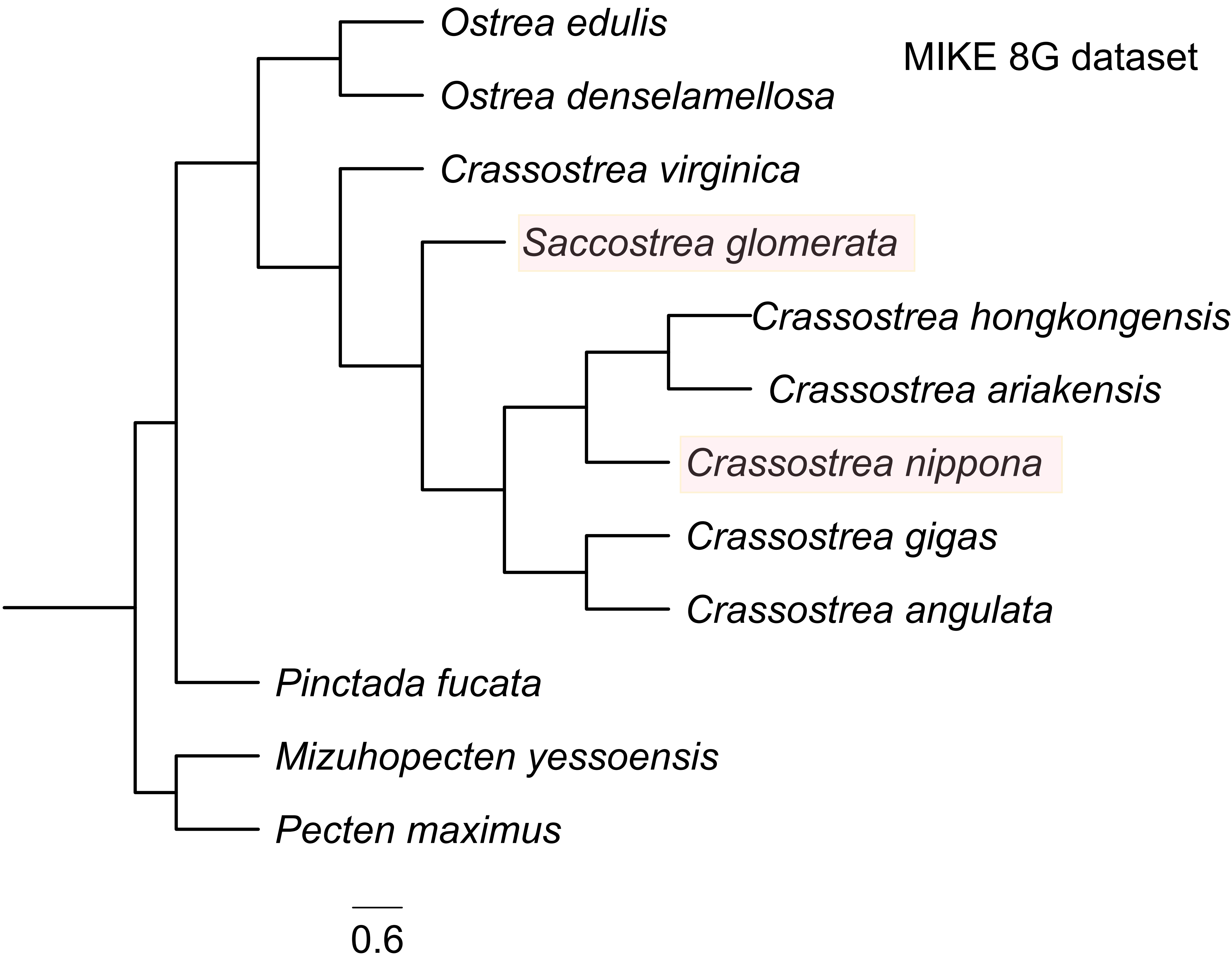

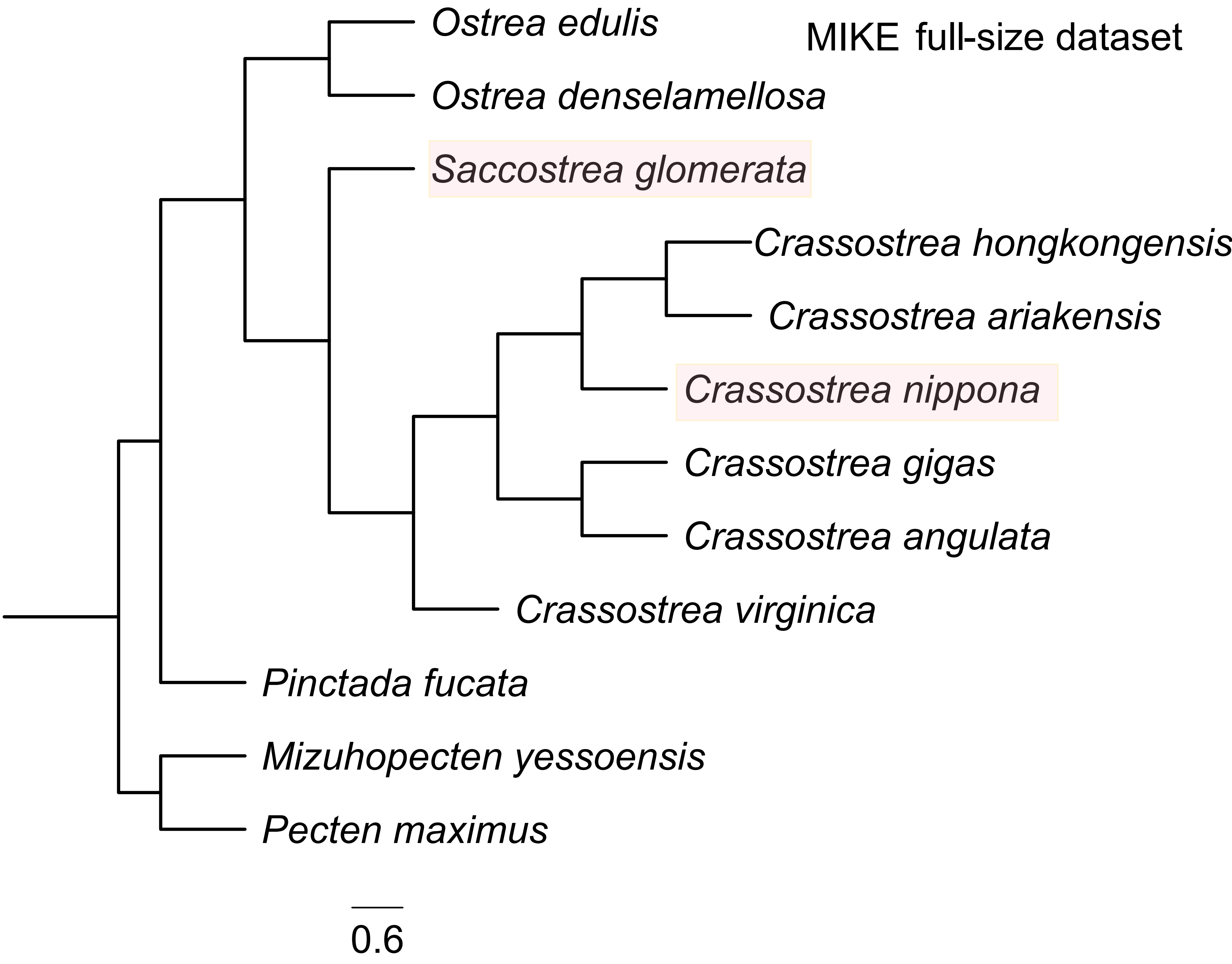

### Supplementary Fig. 6

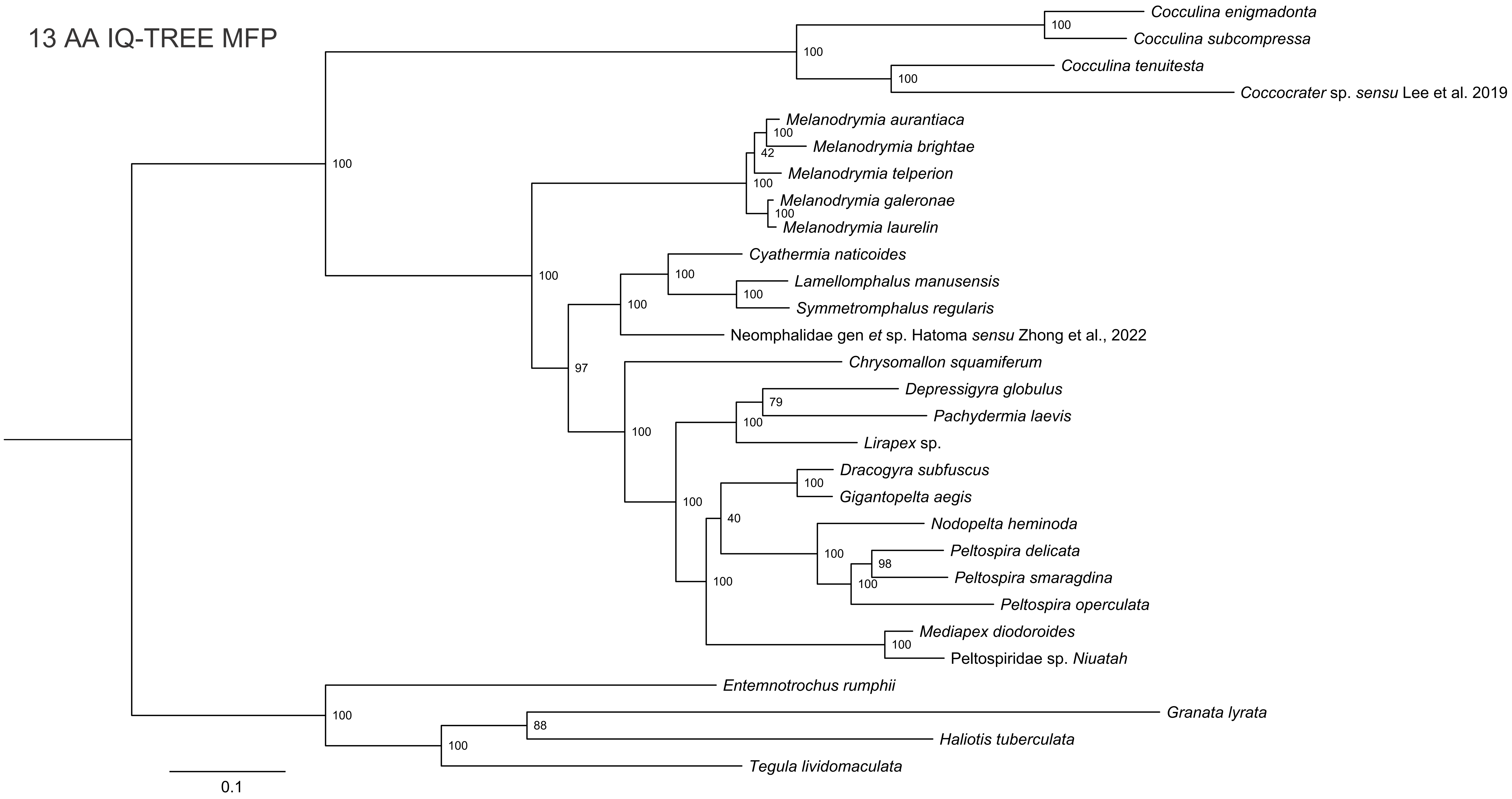

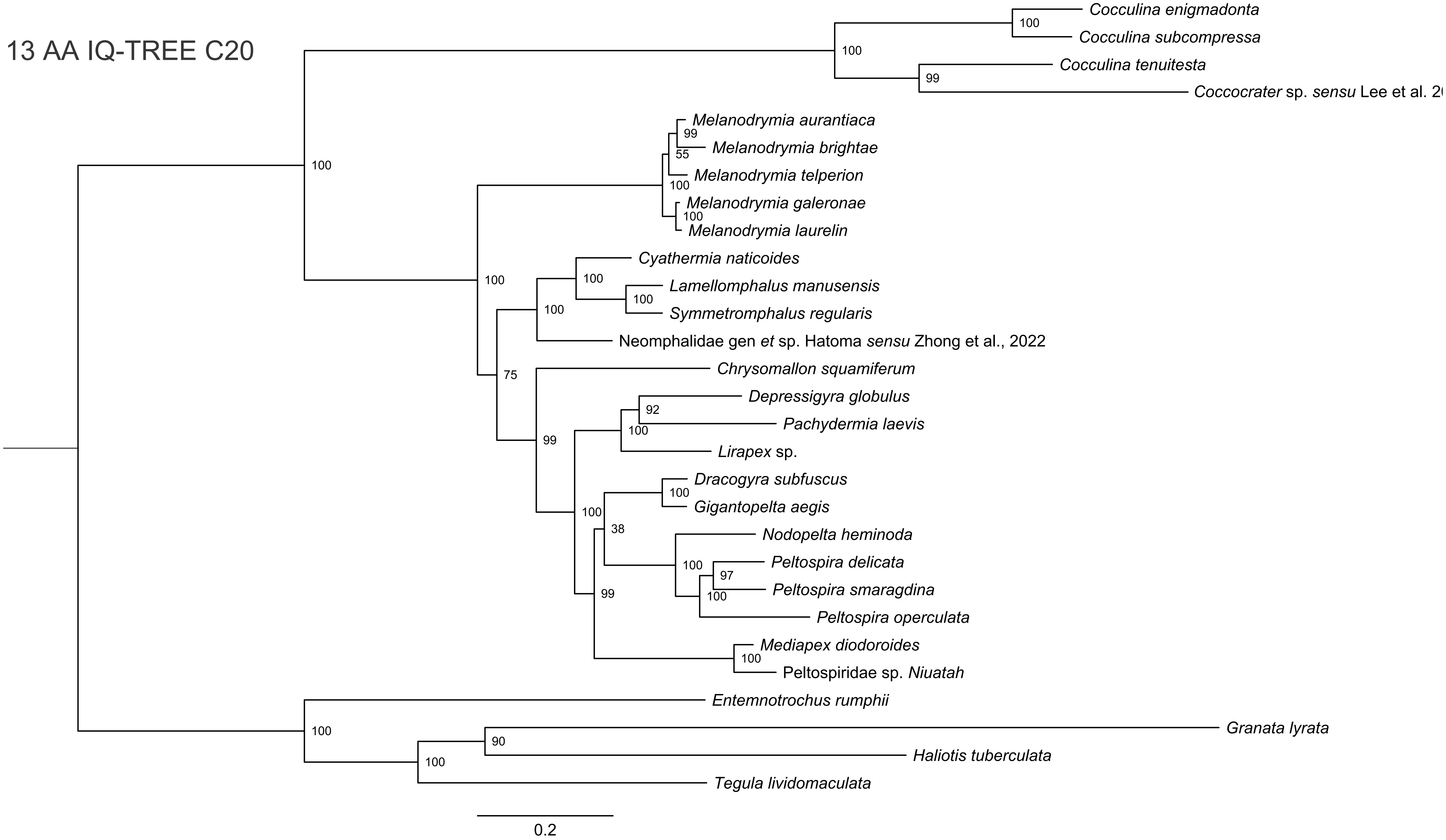

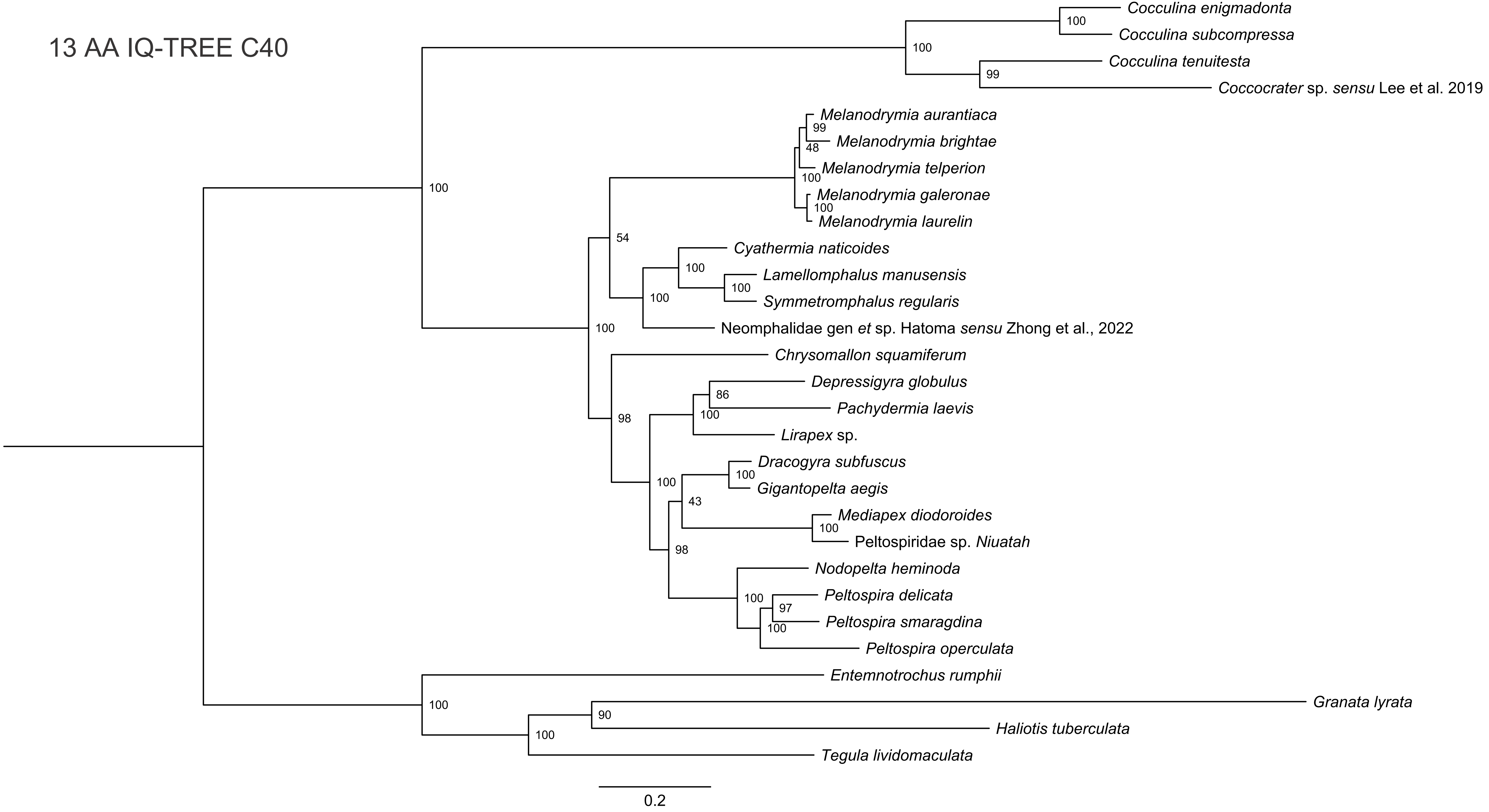

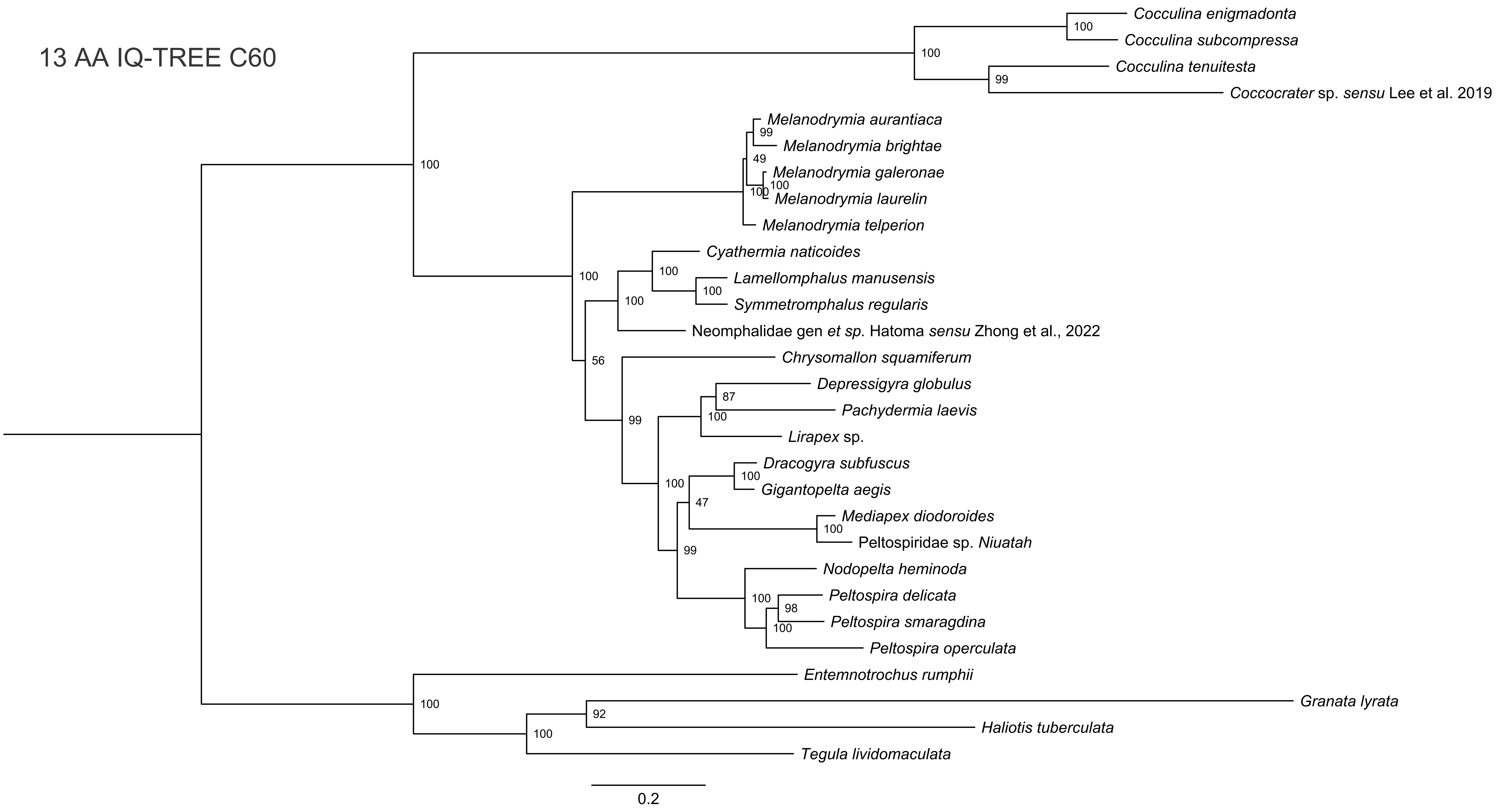

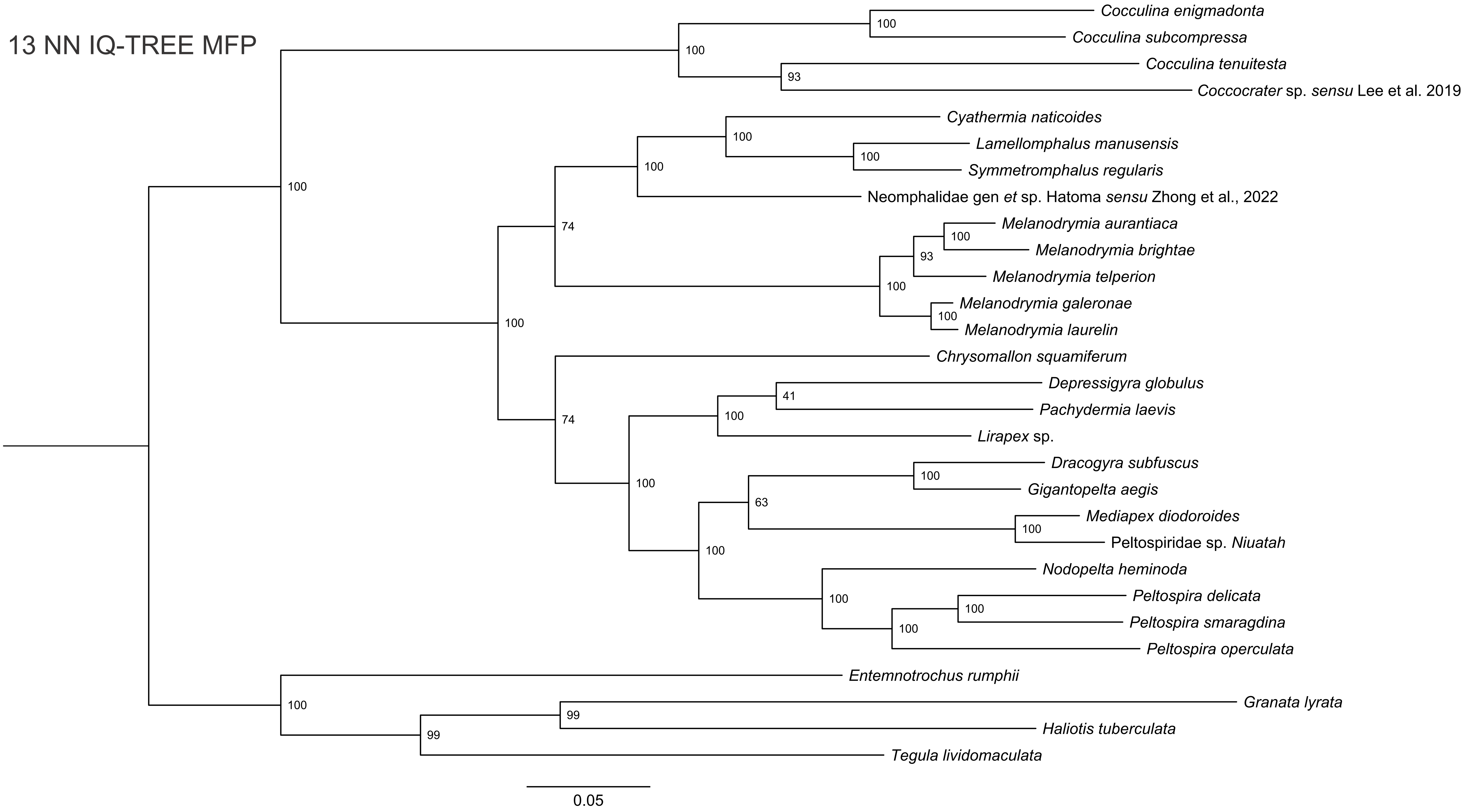

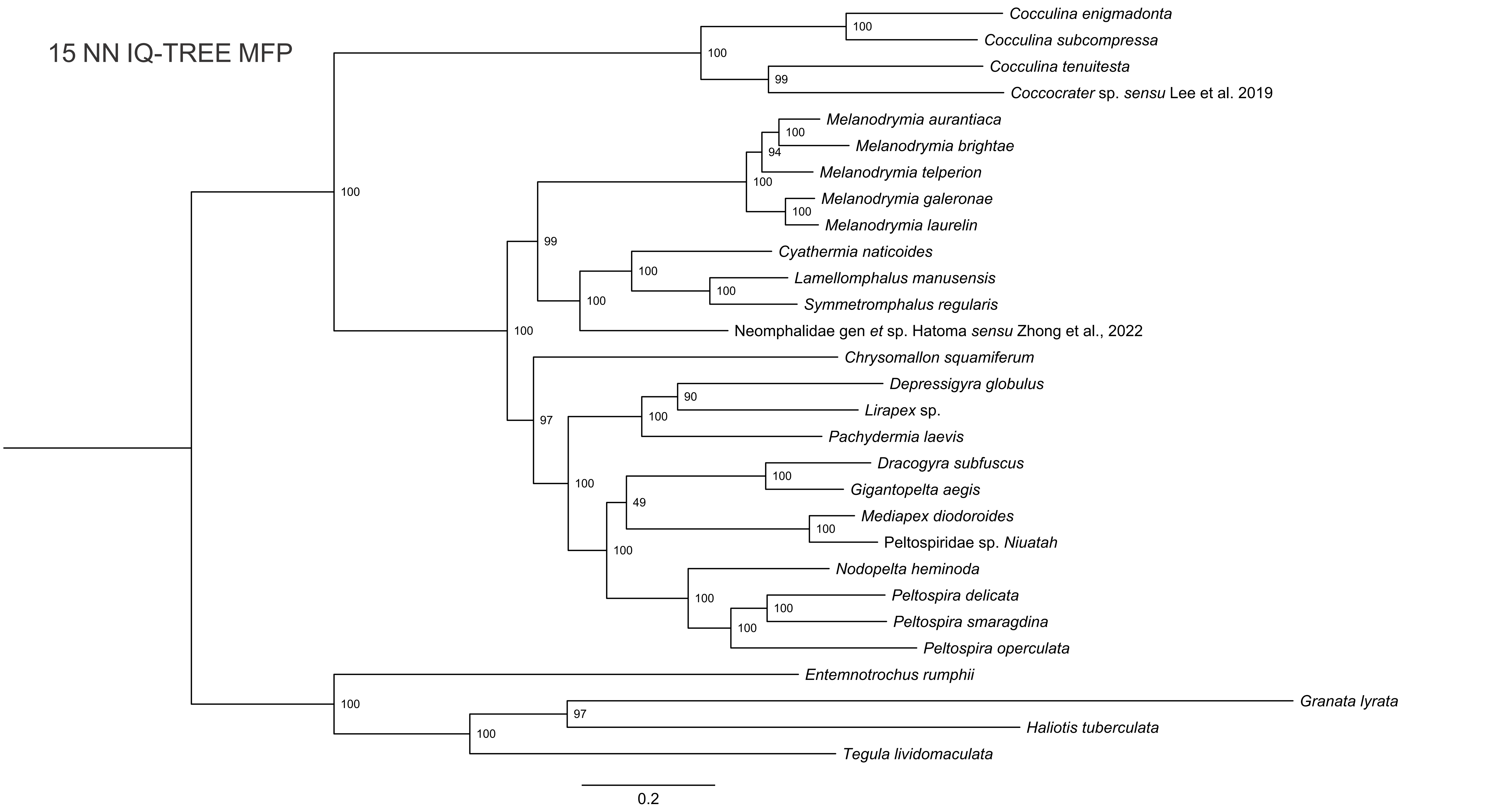

### Supplementary Fig. 7

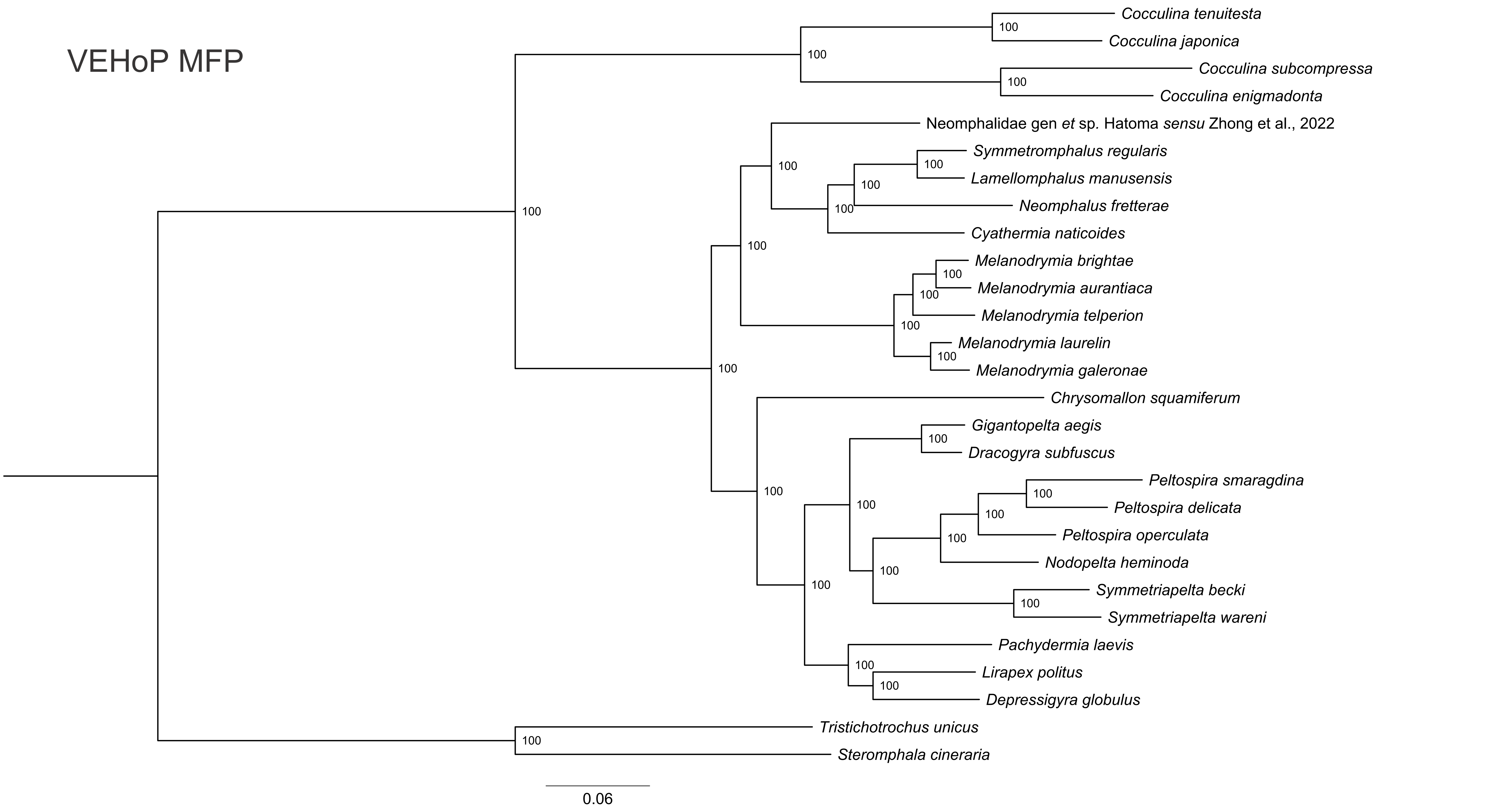

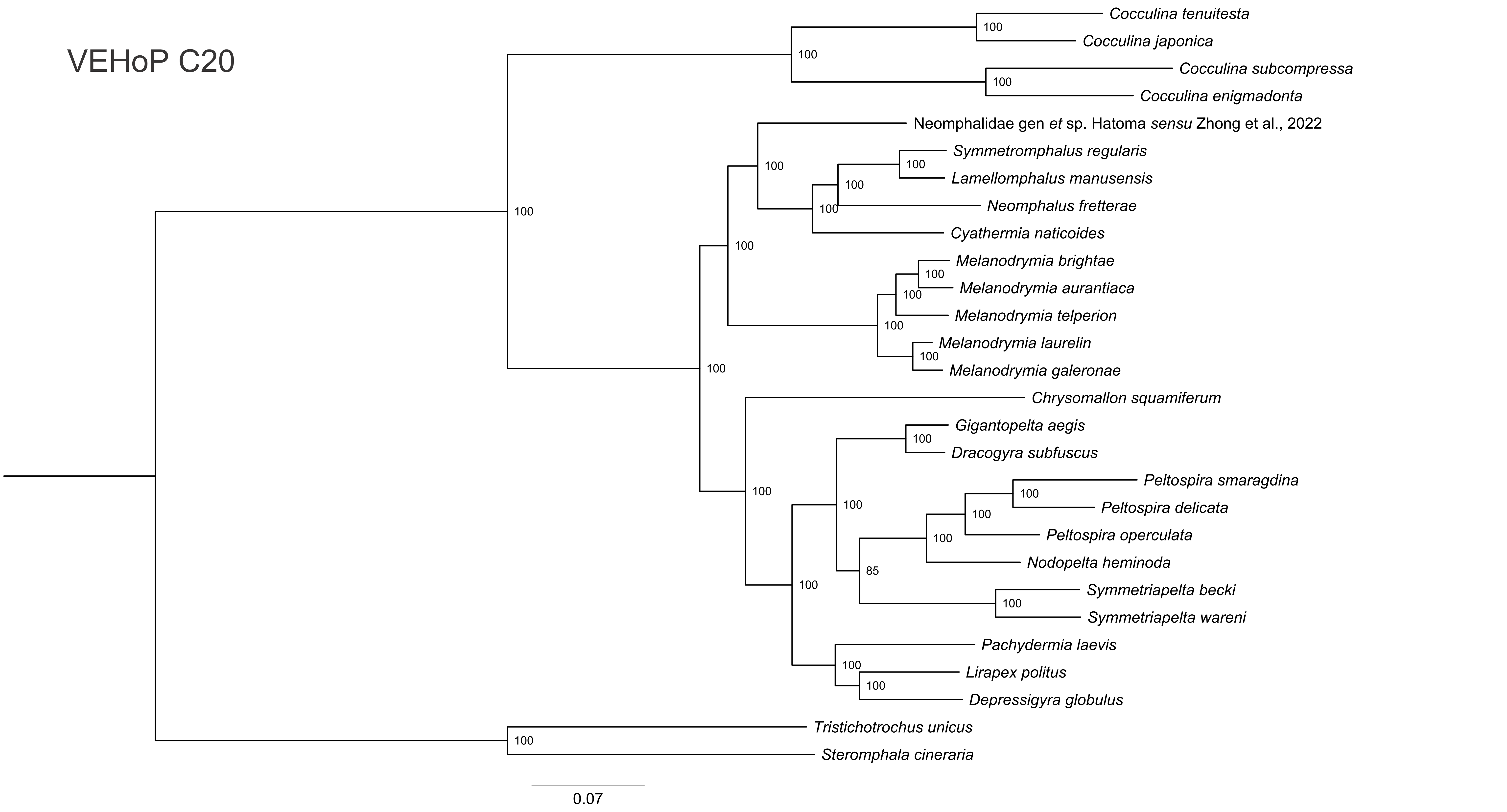

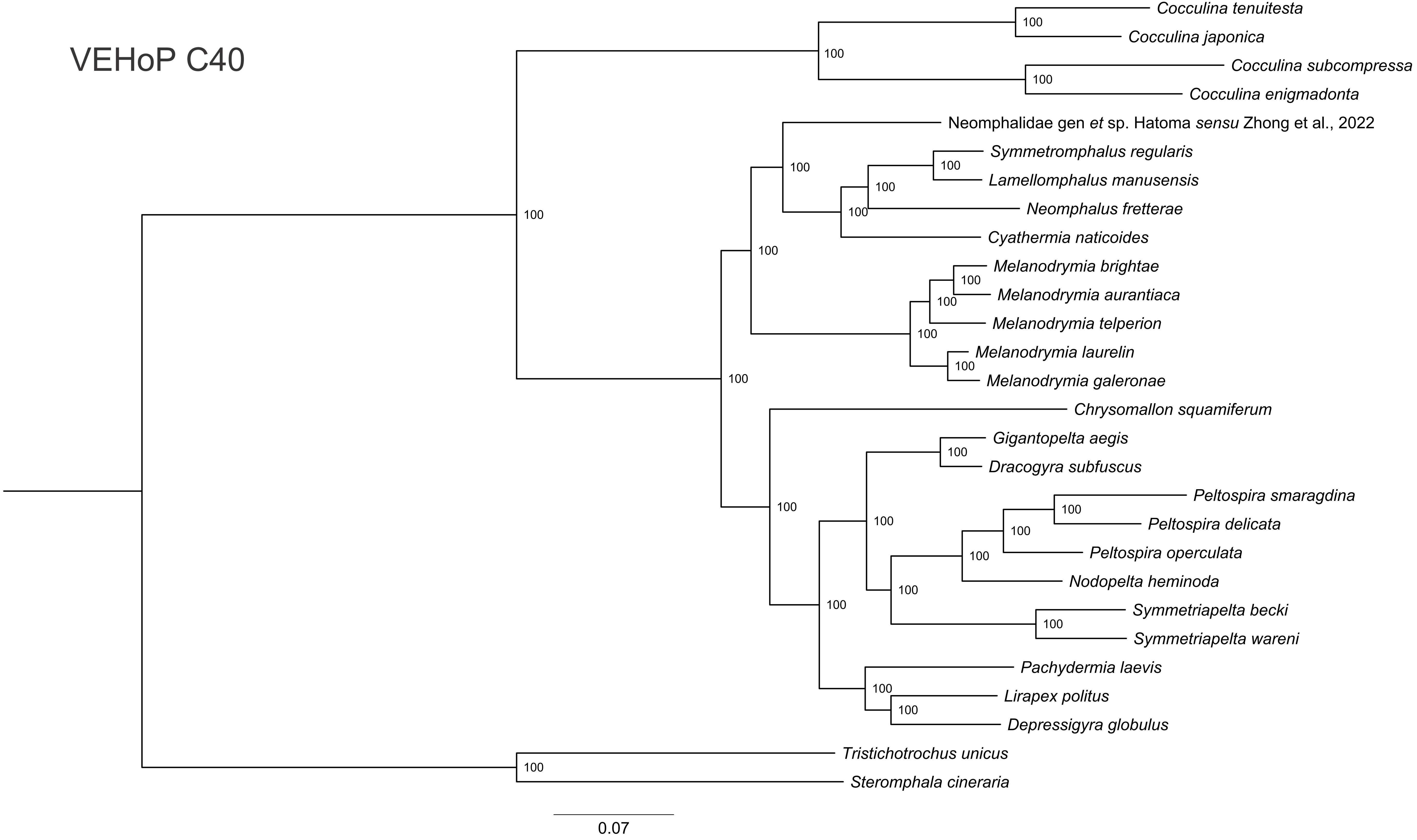

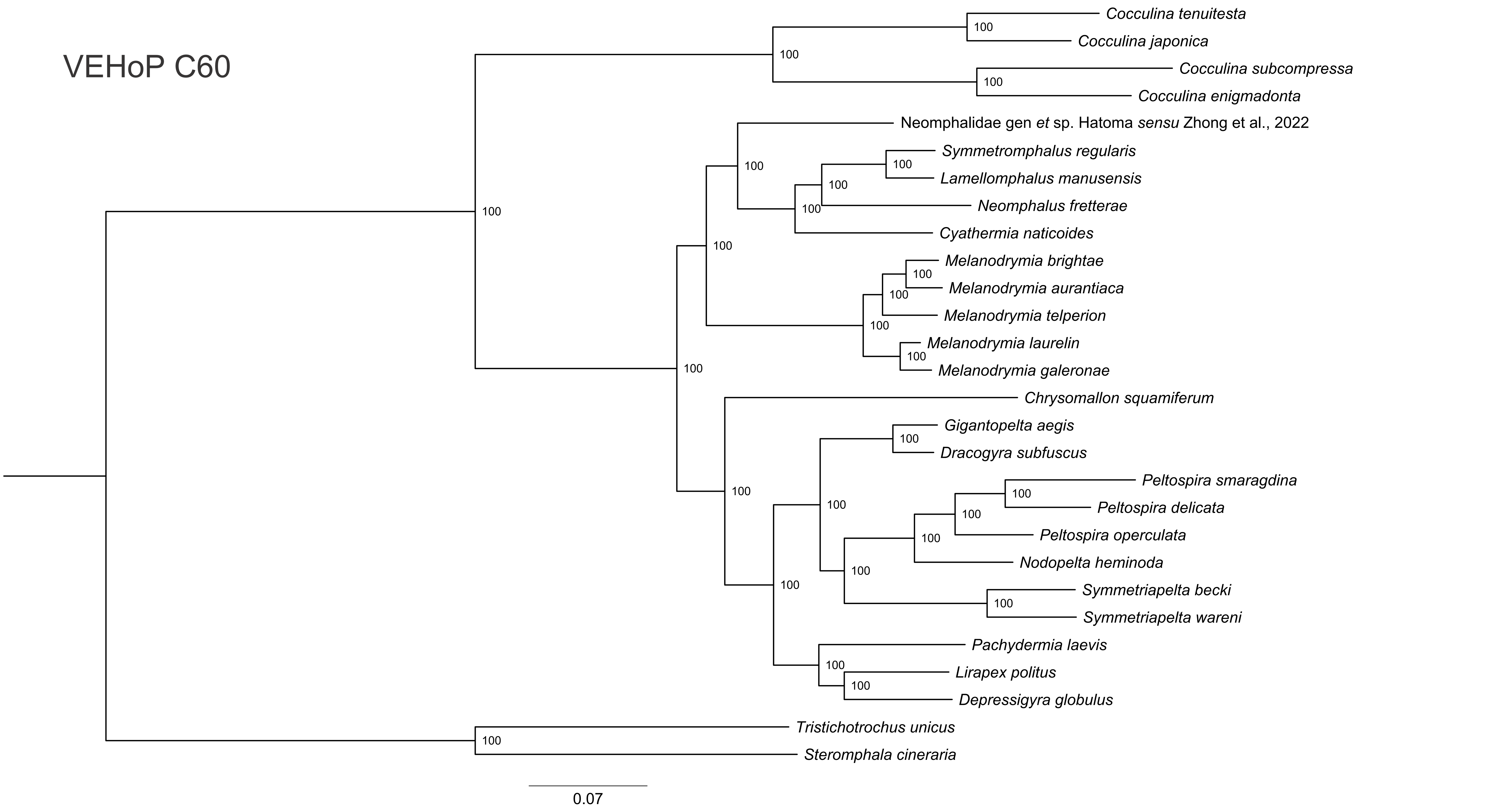

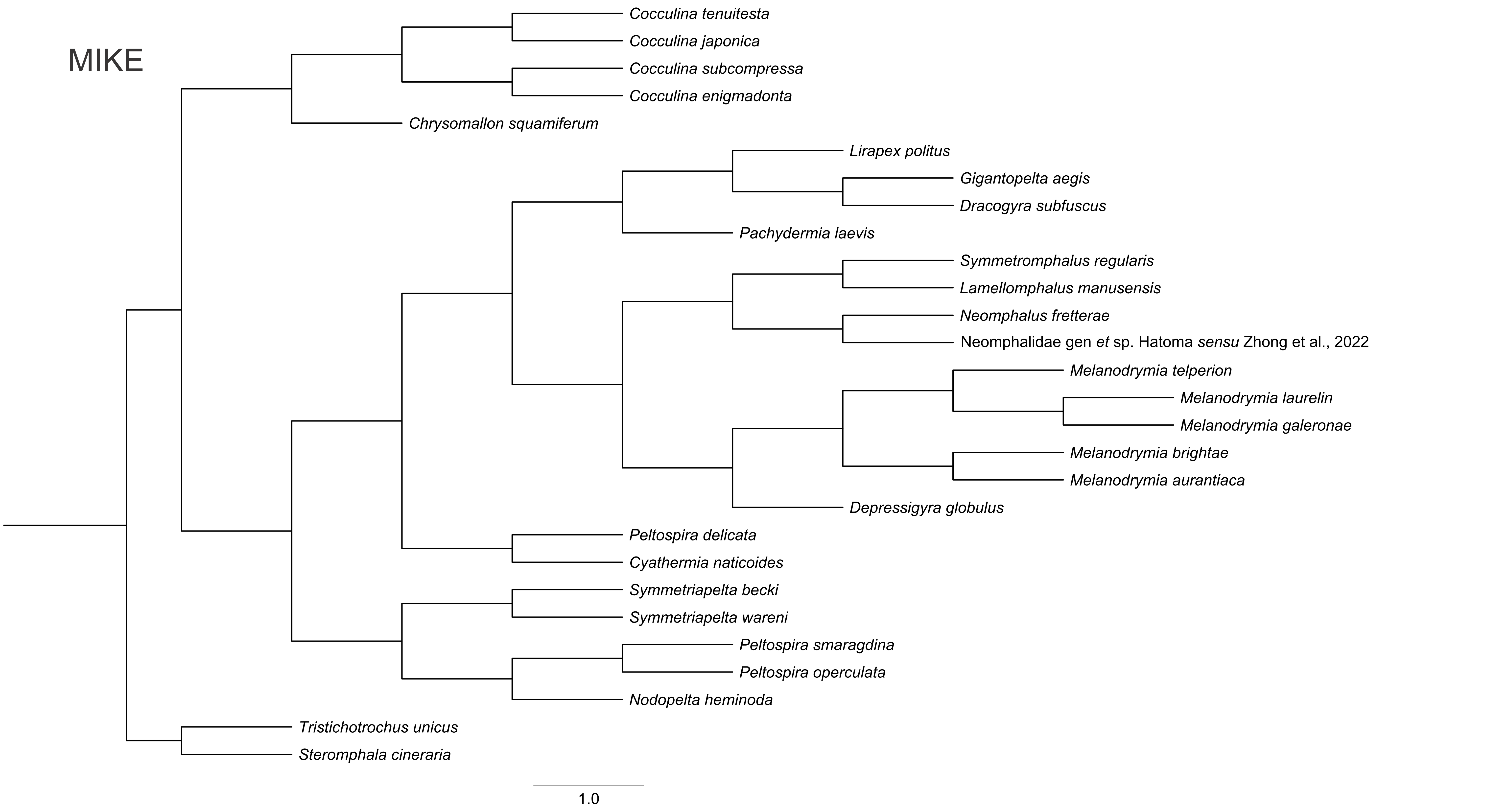

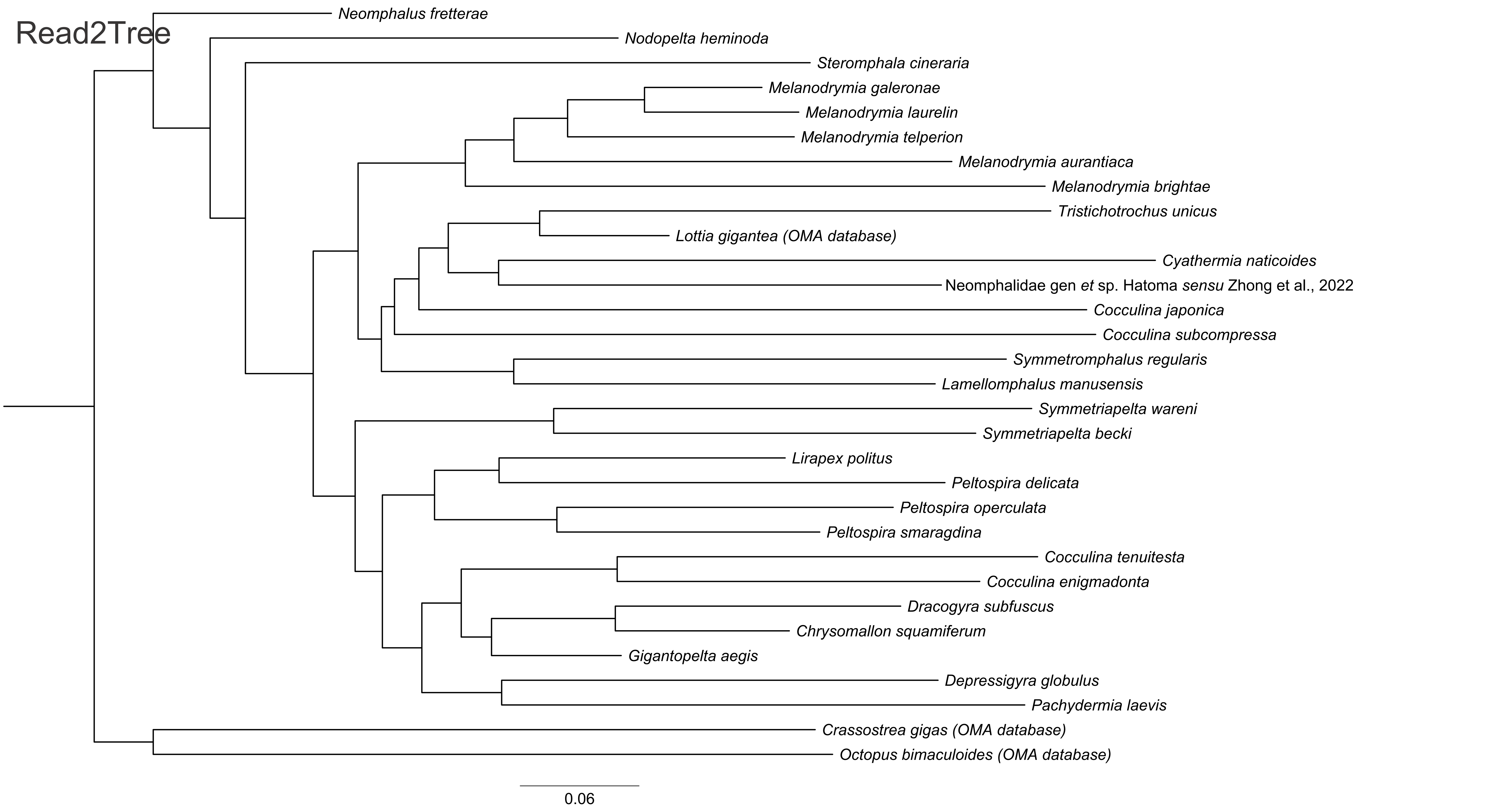
