## Supplementary Fig. 5 for "Reliable inference of phylogenomic relationship via assembly-based strategy accommodating raw reads and proteins"

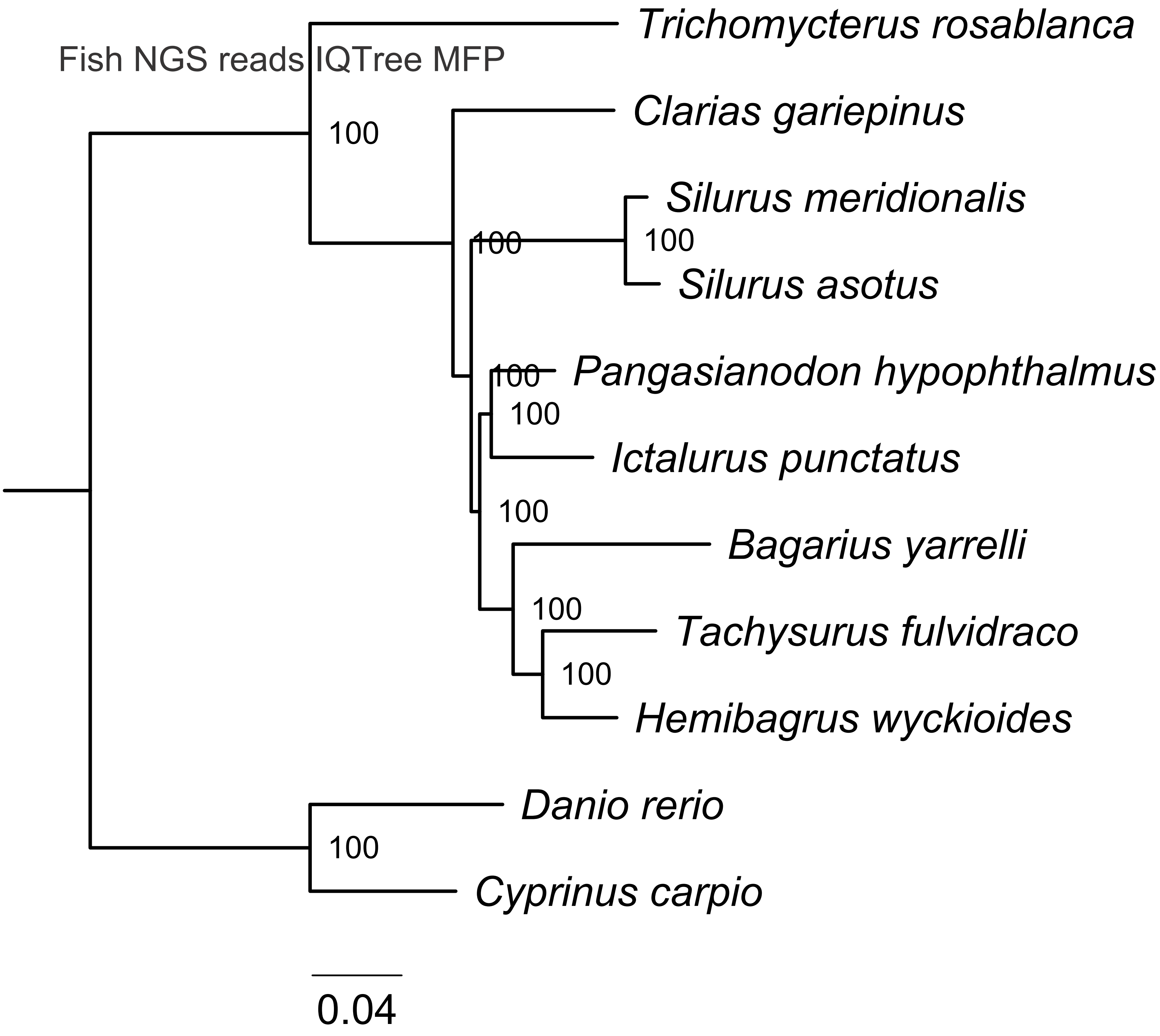
Fish NGS reads IQTree MFP
Trichomycterus rosablanca
Clarias gariepinus
Silurus meridionalis
Silurus asotus
Pangasianodon hypophthalmus
Ictalurus punctatus
Bagarius yarrelli
Tachysurus fulvidraco
Hemibagrus wyckioides
Danio rerio
Cyprinus carpio
100
100
100
100
100
100
100
100
100
100
0.04

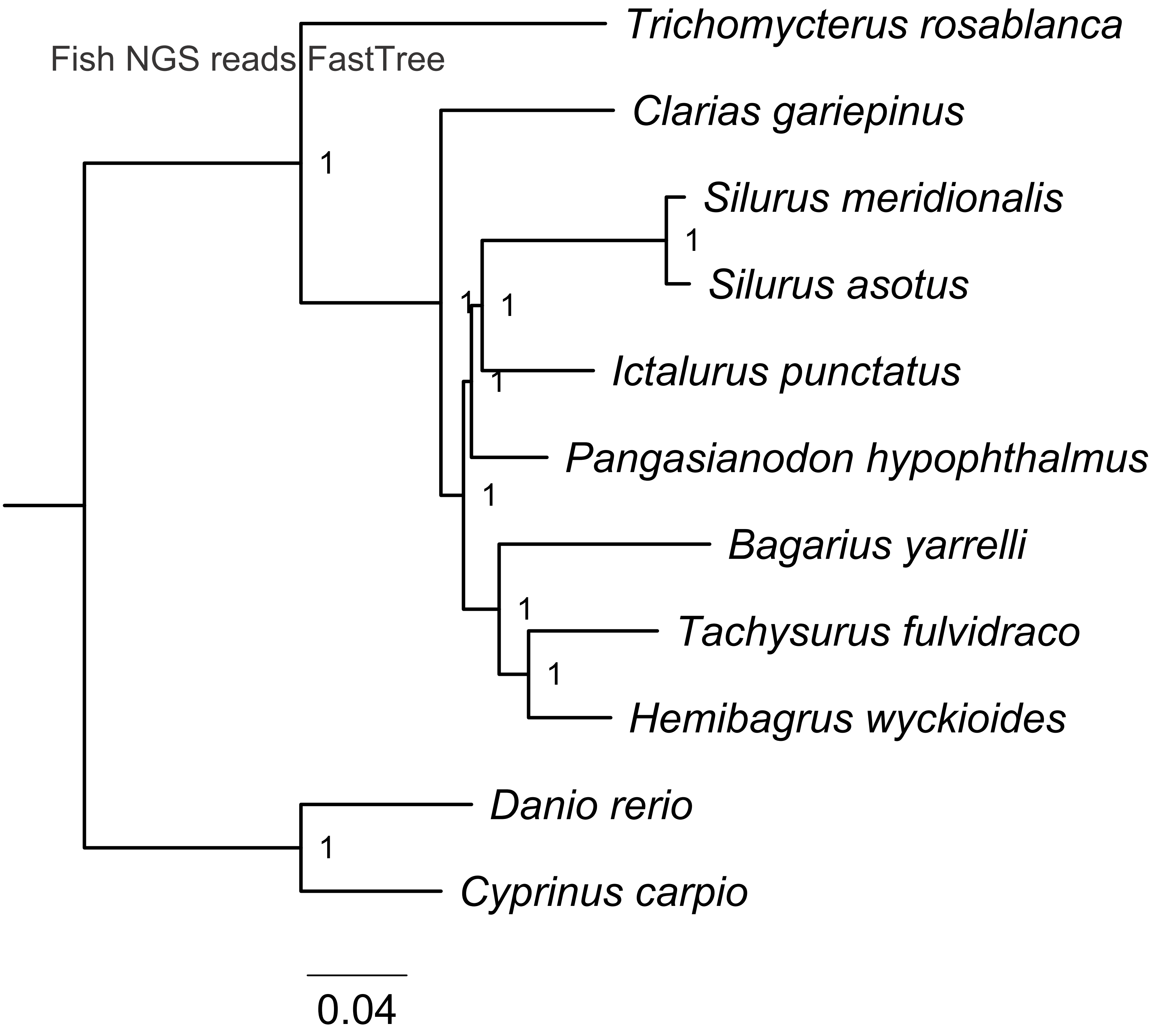
Fish NGS reads FastTree
Trichomycterus rosablanca
Clarias gariepinus
Silurus meridionalis
Silurus asotus
Ictalurus punctatus
Pangasianodon hypophthalmus
Bagarius yarrelli
Tachysurus fulvidraco
Hemibagrus wyckioides
Danio rerio
Cyprinus carpio
1
1
1
1
1
1
1
1
1
0.04

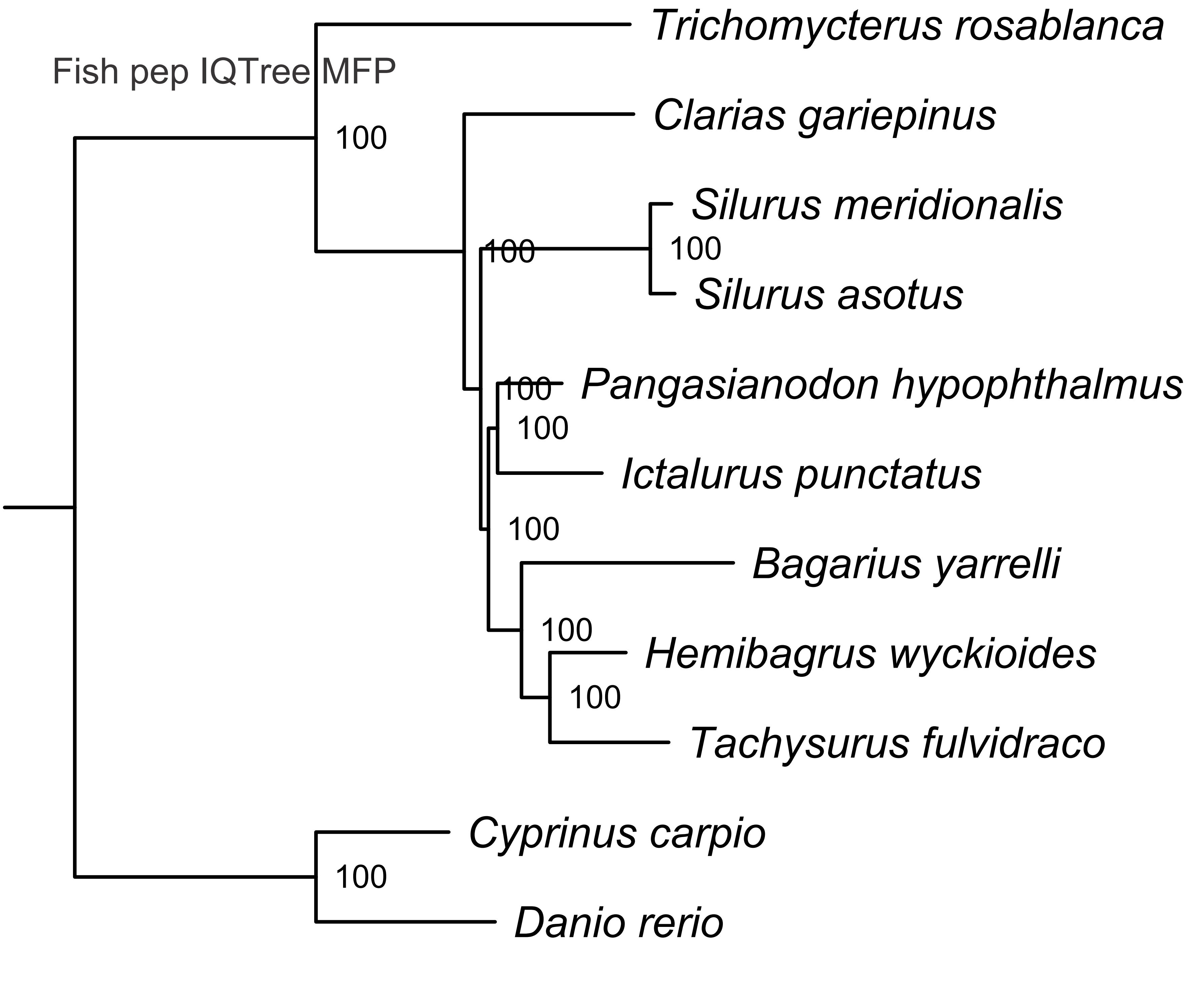

Fish pep IQTree MFP
Trichomycterus rosablanca
Clarias gariepinus
Silurus meridionalis
Silurus asotus
Pangasianodon hypophthalmus
Ictalurus punctatus
Bagarius yarrelli
Hemibagrus wyckioides
Tachysurus fulvidraco
Cyprinus carpio
Danio rerio
100
100
100
100
100
100
100
100
100
100
0.04

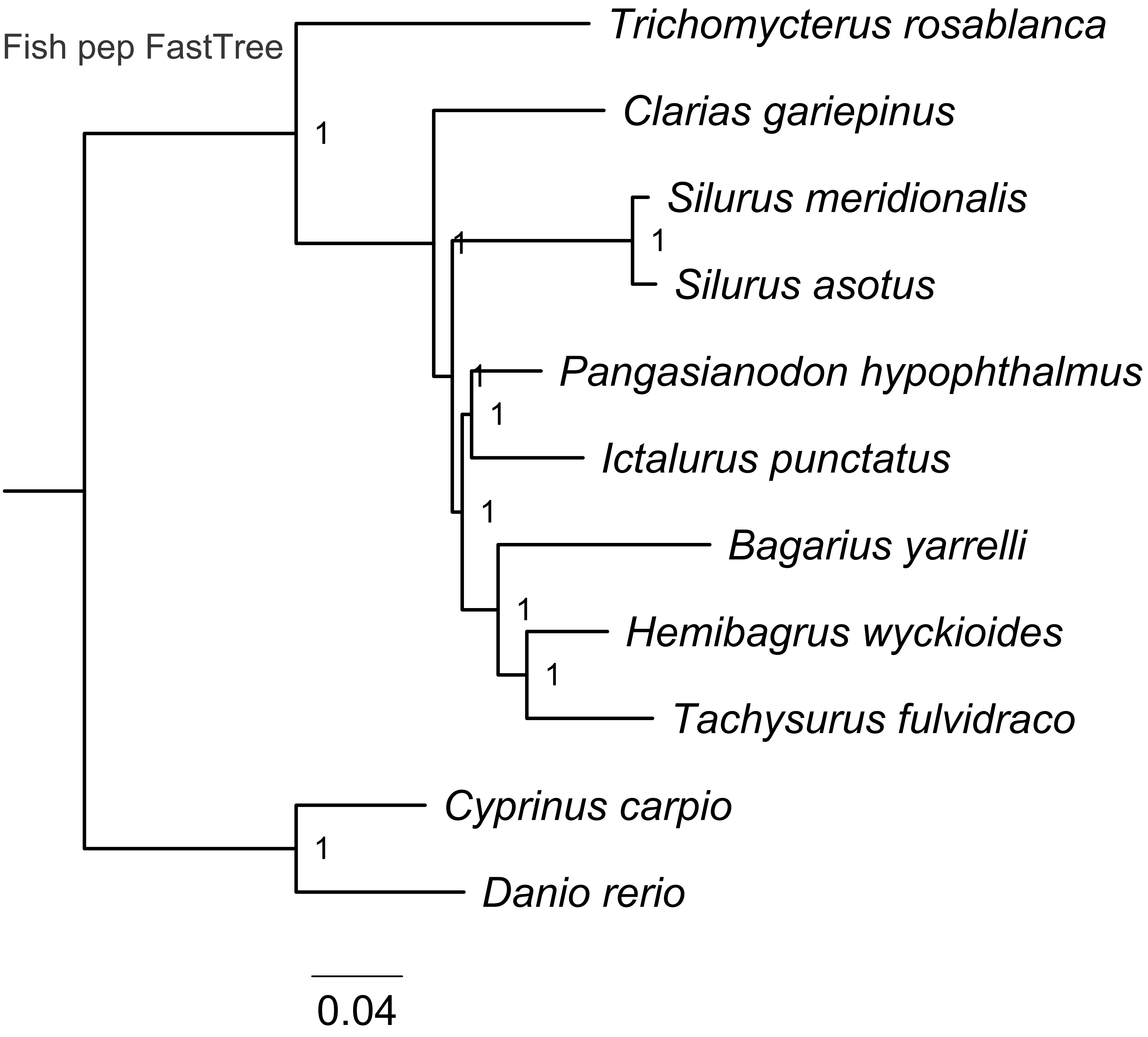
Fish pep FastTree
Trichomycterus rosablanca
Clarias gariepinus
Silurus meridionalis
Silurus asotus
Pangasianodon hypophthalmus
Ictalurus punctatus
Bagarius yarrelli
Hemibagrus wyckioides
Tachysurus fulvidraco
Cyprinus carpio
Danio rerio
1
0.04

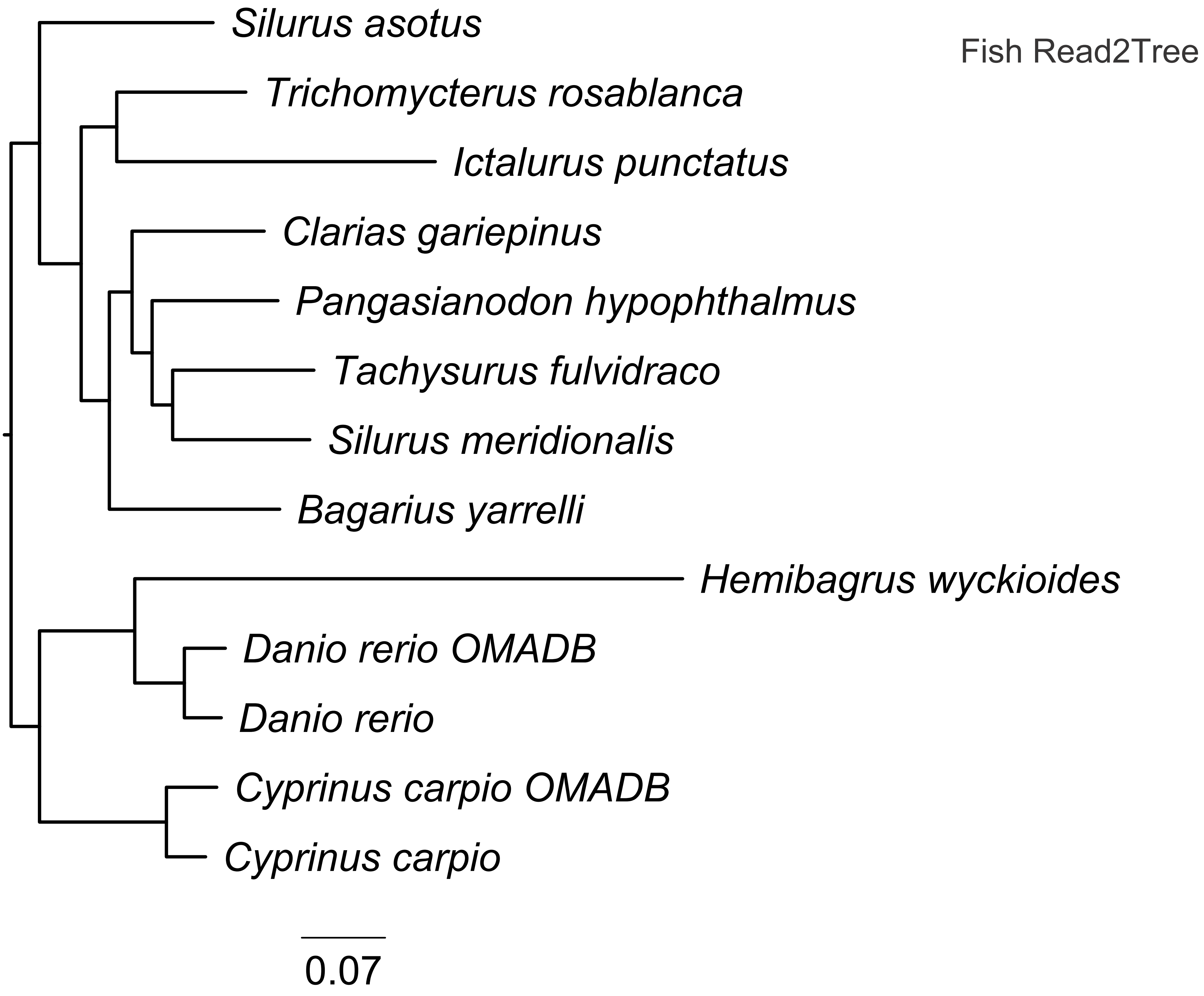
Silurus asotus
Fish Read2Tree
Trichomycterus rosablanca
Ictalurus punctatus
Clarias gariepinus
Pangasianodon hypophthalmus
Tachysurus fulvidraco
Silurus meridionalis
Bagarius yarrelli
Hemibagrus wyckioides
Danio rerio OMADB
Danio rerio
Cyprinus carpio OMADB
Cyprinus carpio
0.07

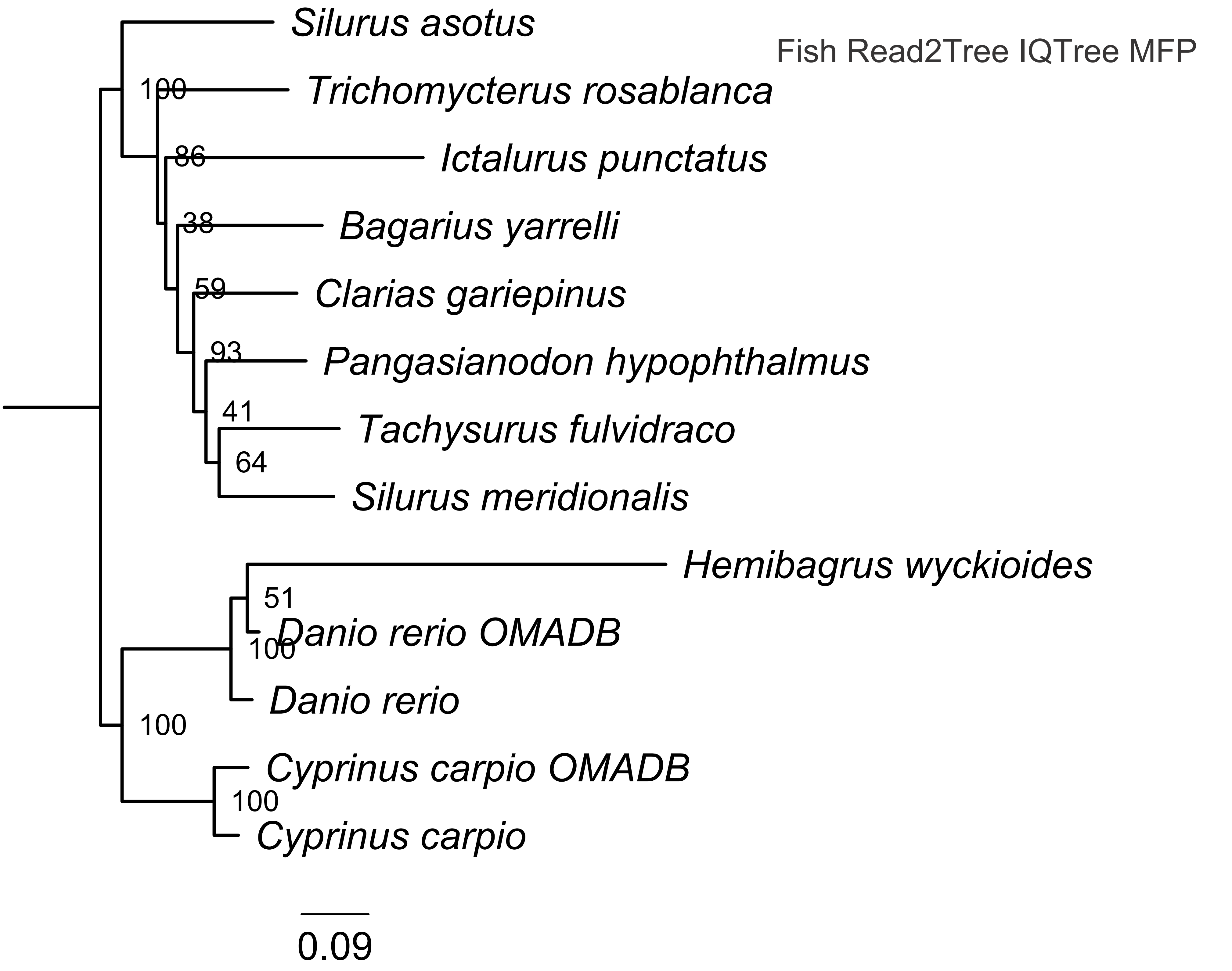

Fish Read2Tree IQTree MFP
Silurus asotus
Trichomycterus rosablanca
Ictalurus punctatus
Bagarius yarrelli
Clarias gariepinus
Pangasianodon hypophthalmus
Tachysurus fulvidraco
Silurus meridionalis
Hemibagrus wyckioides
Danio rerio OMADB
Danio rerio
Cyprinus carpio OMADB
Cyprinus carpio
100
86
38
59
93
41
64
51
100
100
100
0.09

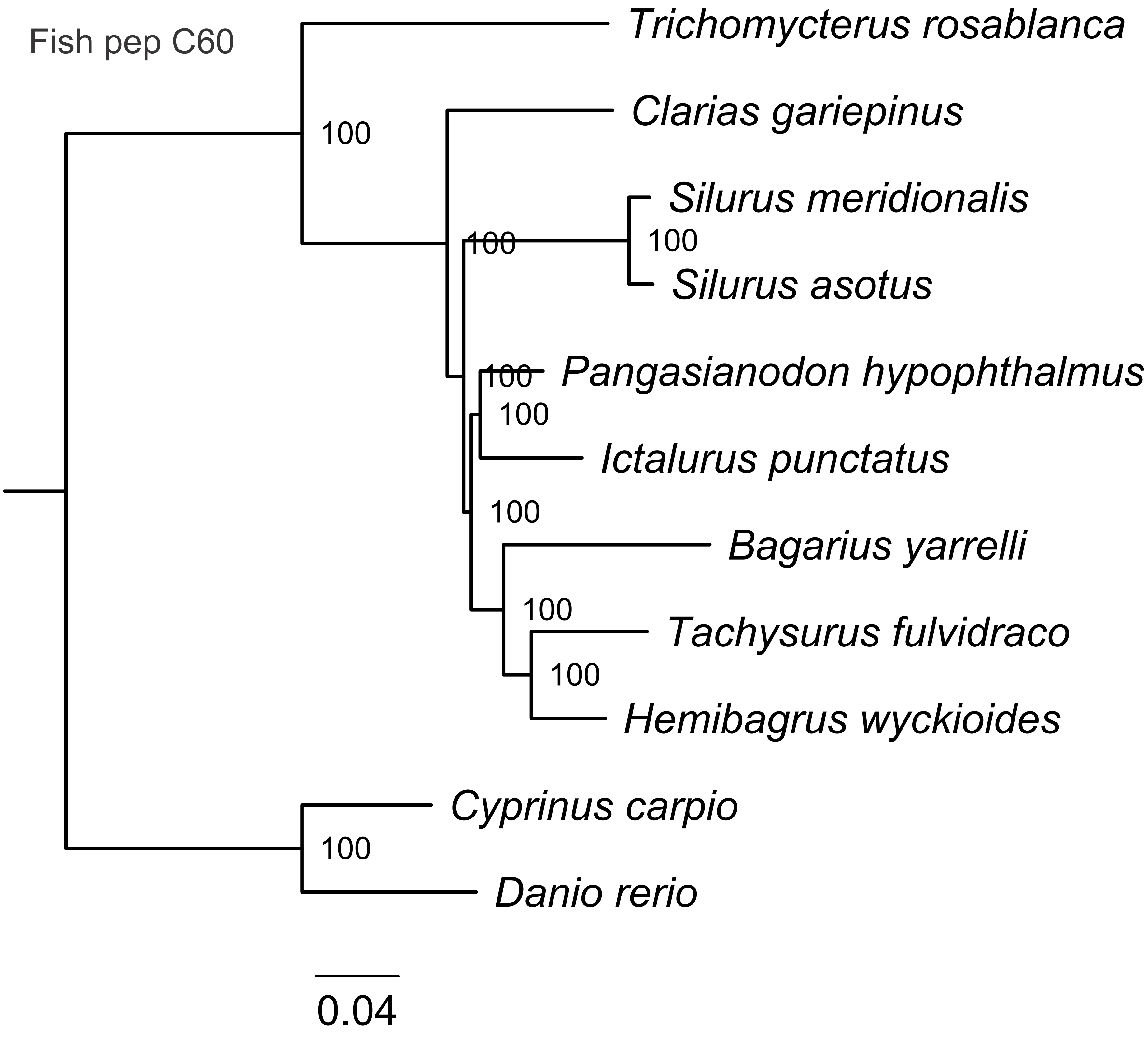
Fish pep C60
Trichomycterus rosablanca
Clarias gariepinus
Silurus meridionalis
Silurus asotus
Pangasianodon hypophthalmus
Ictalurus punctatus
Bagarius yarrelli
Tachysurus fulvidraco
Hemibagrus wyckioides
Cyprinus carpio
Danio rerio
100
100
100
100
100
100
100
100
100
0.04

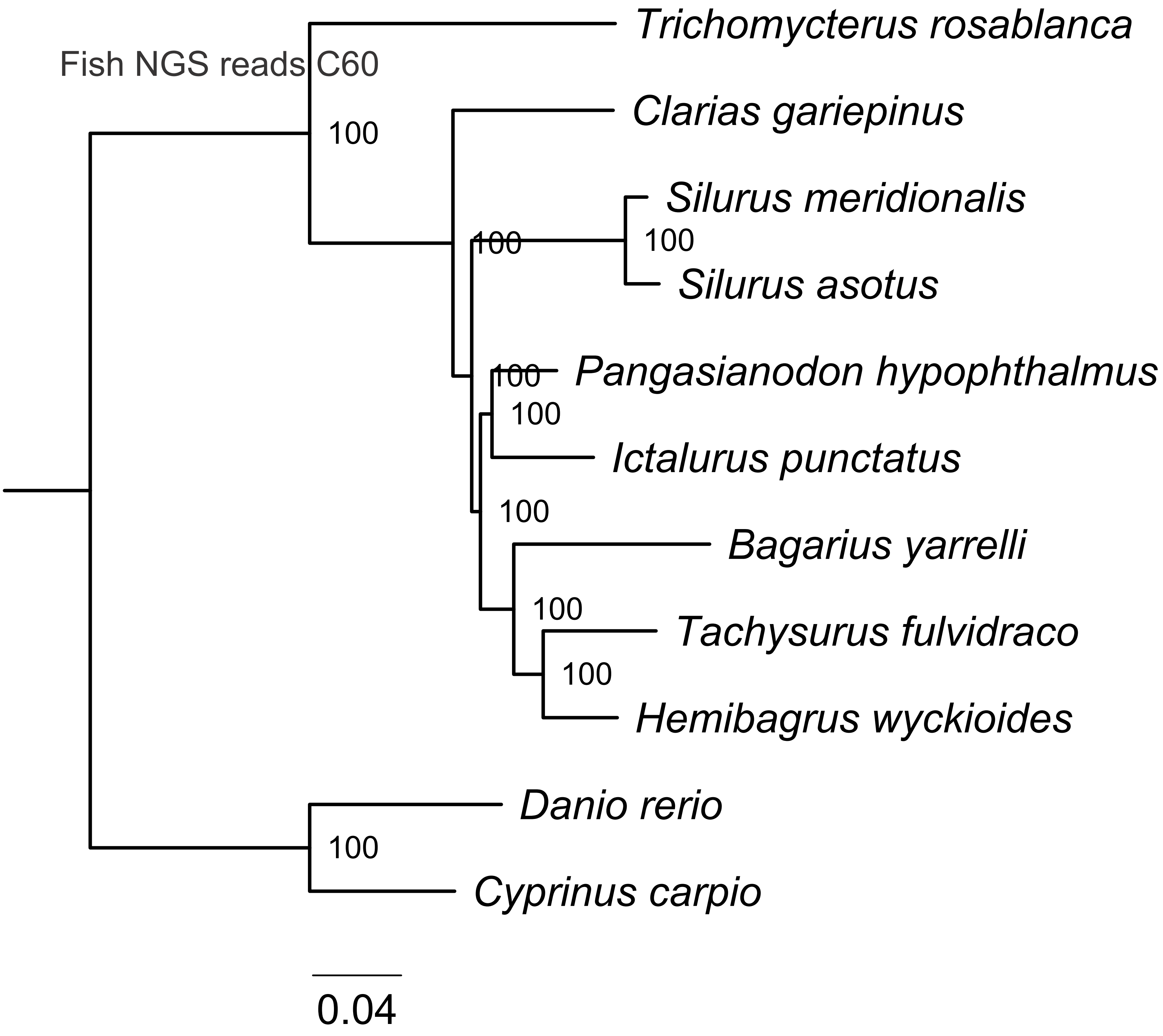

Fish NGS reads C60
Trichomycterus rosablanca
Clarias gariepinus
Silurus meridionalis
Silurus asotus
Pangasianodon hypophthalmus
Ictalurus punctatus
Bagarius yarrelli
Tachysurus fulvidraco
Hemibagrus wyckioides
Danio rerio
Cyprinus carpio
100
0.04

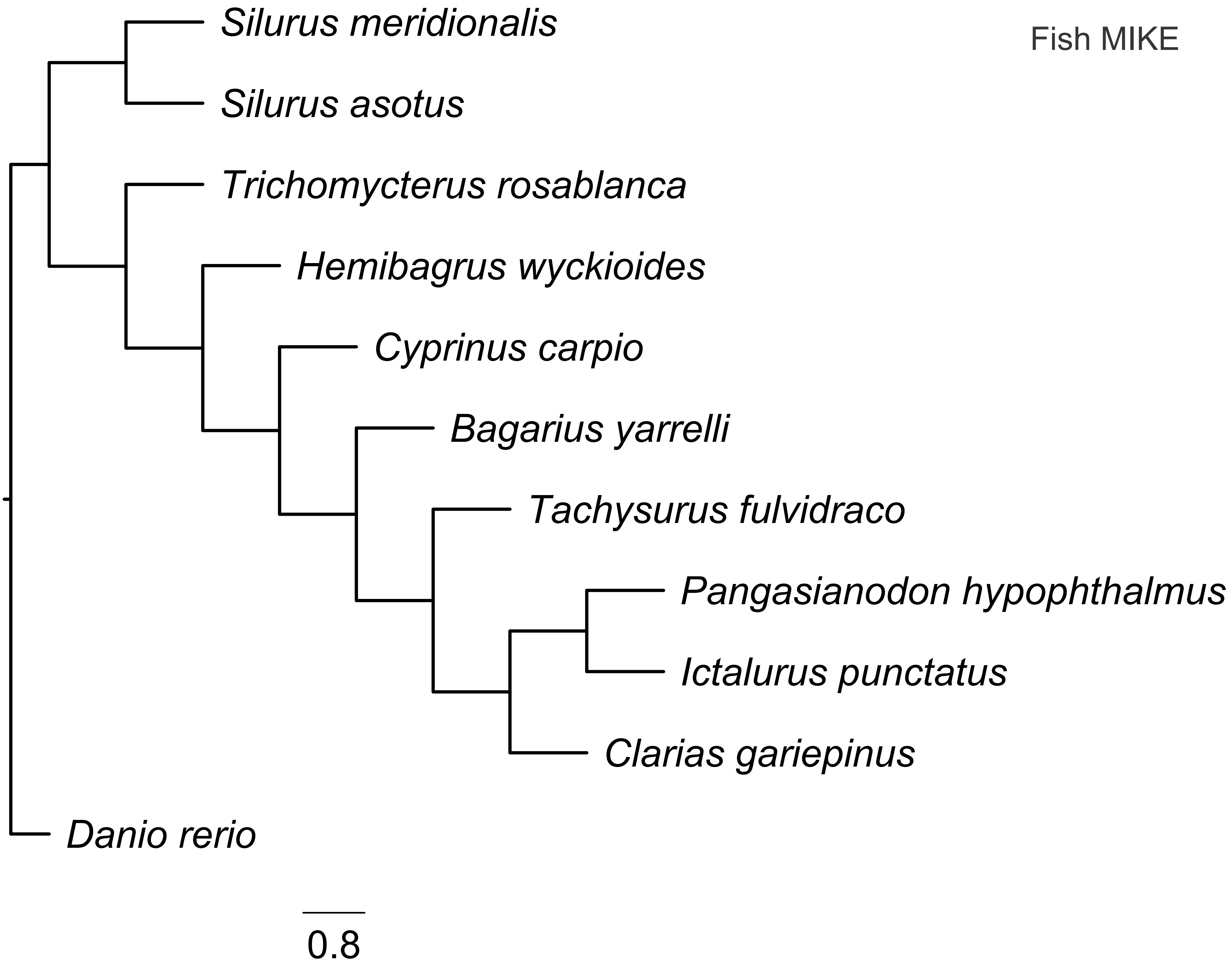
Fish MIKE
Silurus meridionalis
Silurus asotus
Trichomycterus rosablanca
Hemibagrus wyckioides
Cyprinus carpio
Bagarius yarrelli
Tachysurus fulvidraco
Pangasianodon hypophthalmus
Ictalurus punctatus
Clarias gariepinus
Danio rerio
0.8

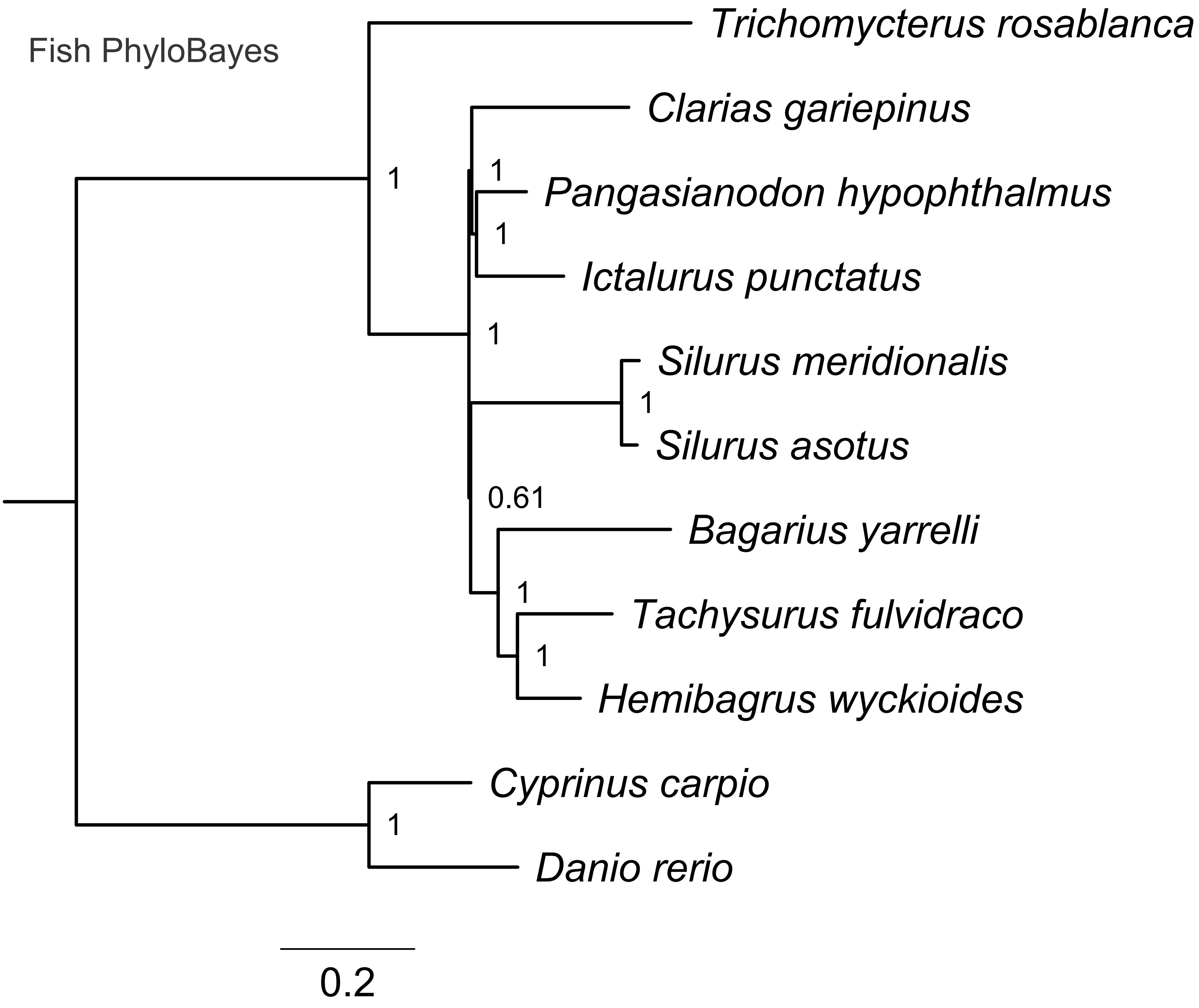
Fish PhyloBayes
Trichomycterus rosablanca
Clarias gariepinus
Pangasianodon hypophthalmus
Ictalurus punctatus
Silurus meridionalis
Silurus asotus
Bagarius yarrelli
Tachysurus fulvidraco
Hemibagrus wyckioides
Cyprinus carpio
Danio rerio
1
1
1
1
1
0.61
1
1
1
0.2

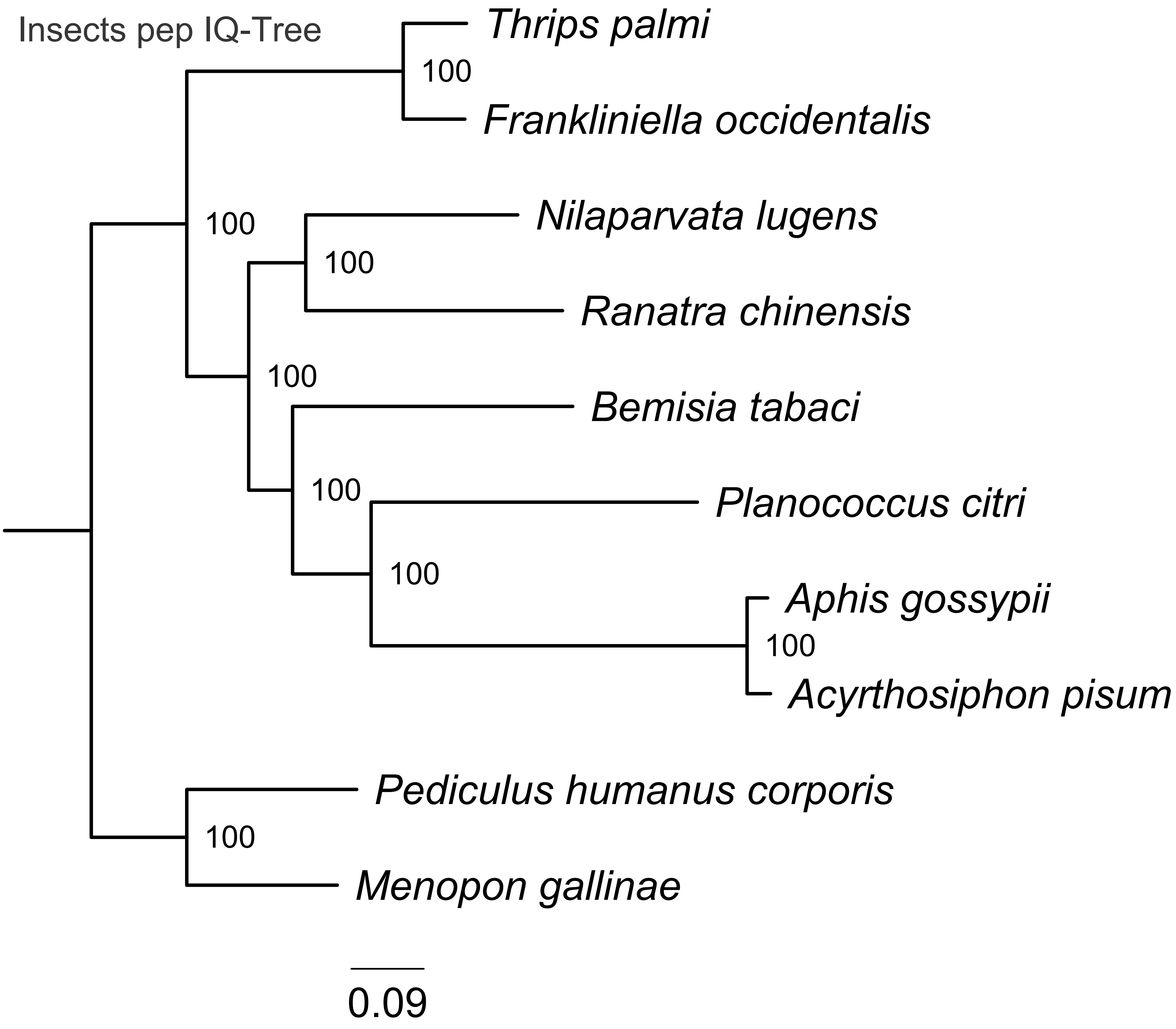

Insects pep IQ-Tree
Thrips palmi
100
Frankliniella occidentalis
100
Nilaparvata lugens
100
Ranatra chinensis
100
Bemisia tabaci
100
Planococcus citri
100
Aphis gossypii
100
Acyrthosiphon pisum
Pediculus humanus corporis
100
Menopon gallinae
0.09

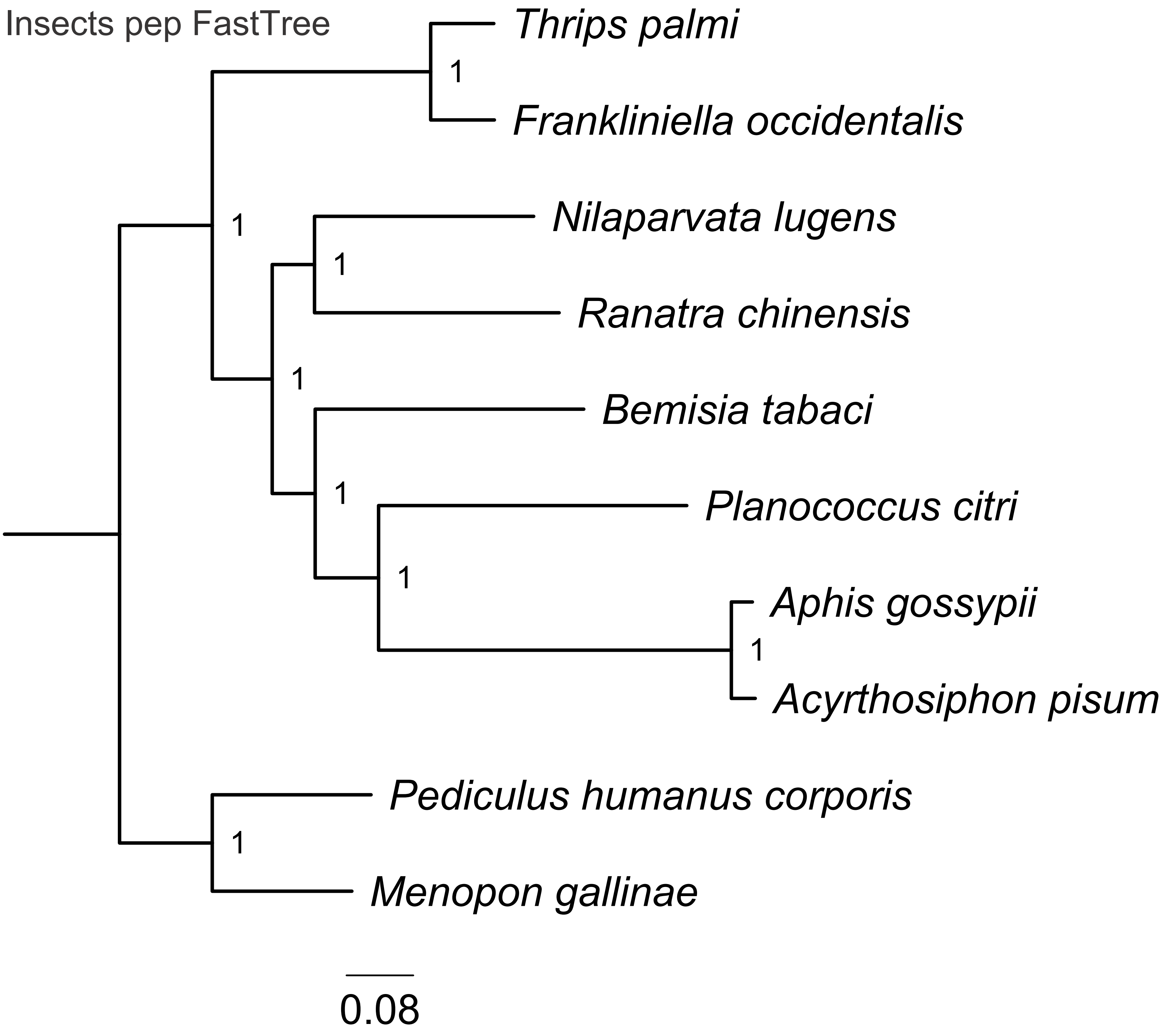

Insects pep FastTree
Thrips palmi
1
Frankliniella occidentalis
1
Nilaparvata lugens
1
Ranatra chinensis
1
Bemisia tabaci
1
Planococcus citri
1
Aphis gossypii
1
Acyrthosiphon pisum
Pediculus humanus corporis
1
Menopon gallinae
0.08

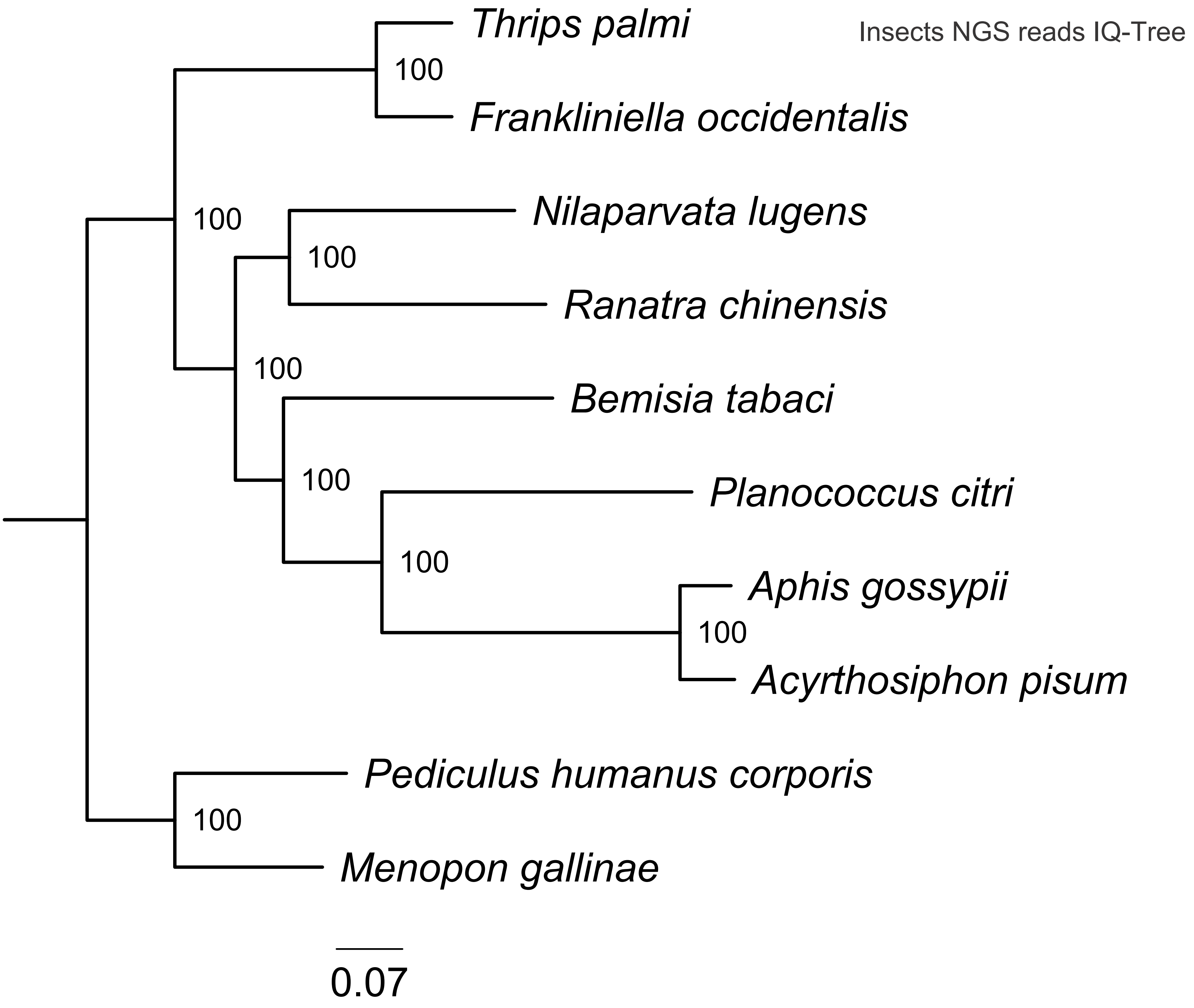
Insects NGS reads IQ-Tree
Thrips palmi
Frankliniella occidentalis
Nilaparvata lugens
Ranatra chinensis
Bemisia tabaci
Planococcus citri
Aphis gossypii
Acyrthosiphon pisum
Pediculus humanus corporis
Menopon gallinae
100
100
100
100
100
100
100
100
0.07

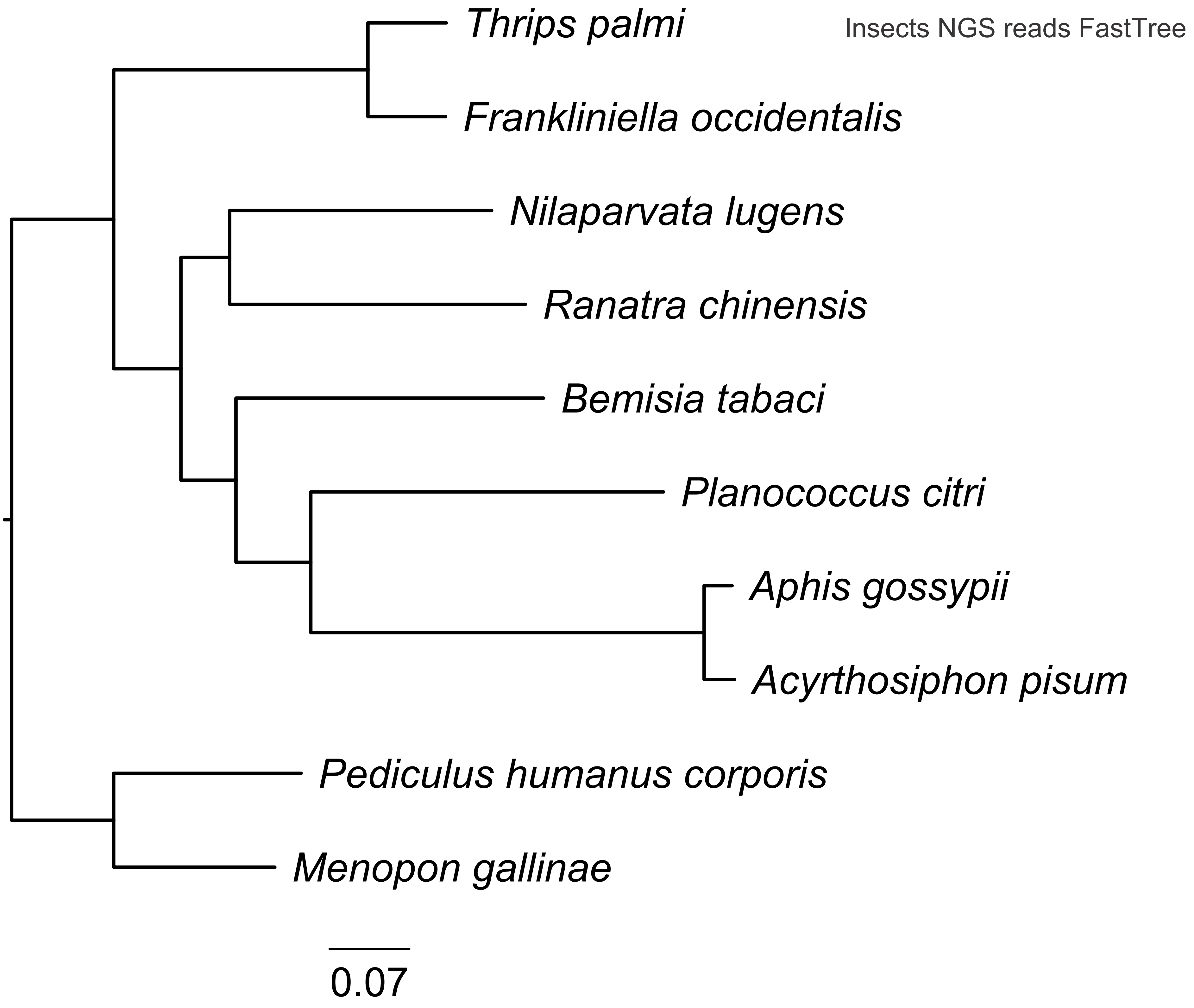
Insects NGS reads FastTree
Thrips palmi
Frankliniella occidentalis
Nilaparvata lugens
Ranatra chinensis
Bemisia tabaci
Planococcus citri
Aphis gossypii
Acyrthosiphon pisum
Pediculus humanus corporis
Menopon gallinae
0.07

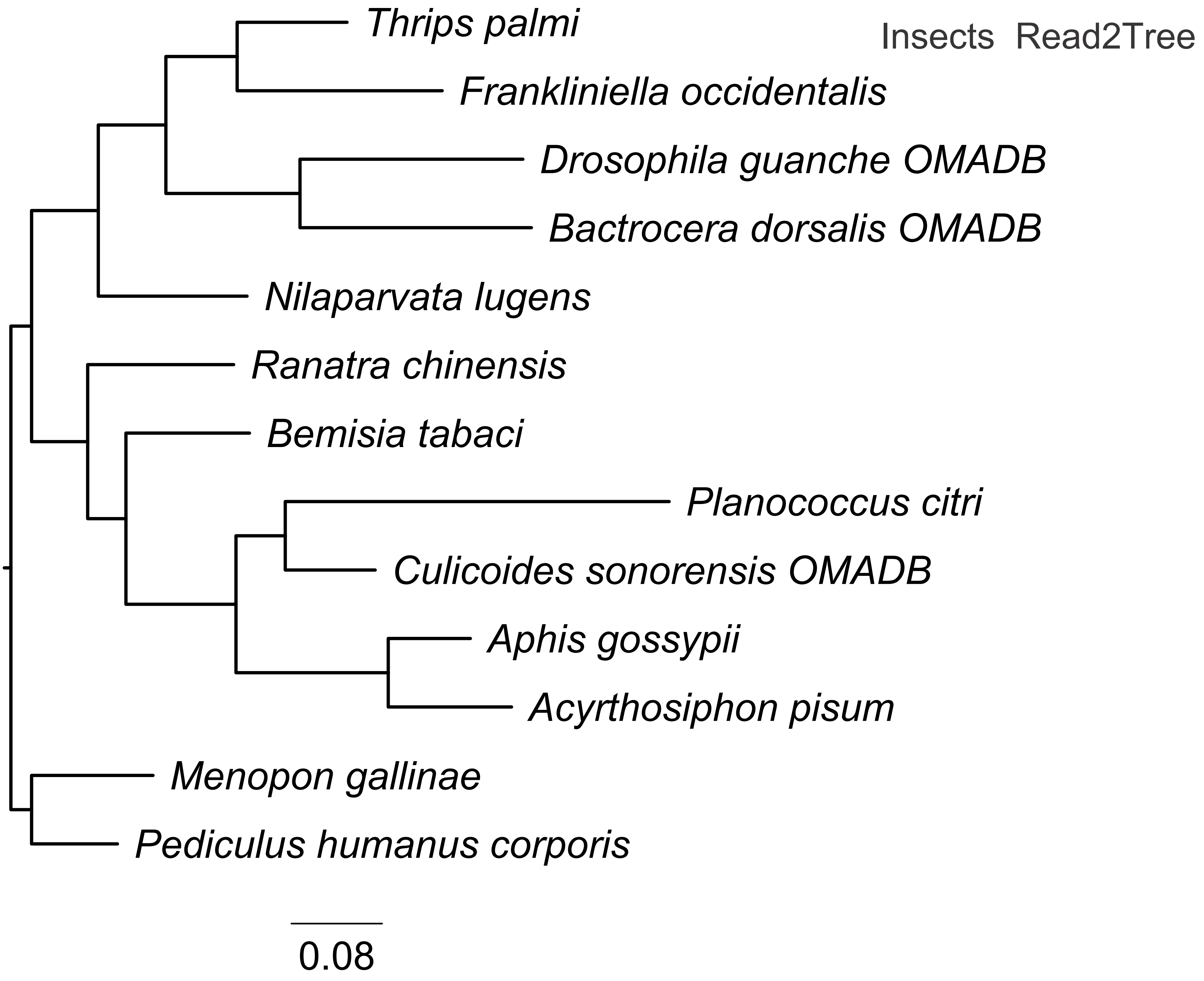
Insects Read2Tree
Thrips palmi
Frankliniella occidentalis
Drosophila guanche OMADB
Bactrocera dorsalis OMADB
Nilaparvata lugens
Ranatra chinensis
Bemisia tabaci
Planococcus citri
Culicoides sonorensis OMADB
Aphis gossypii
Acyrthosiphon pisum
Menopon gallinae
Pediculus humanus corporis
0.08

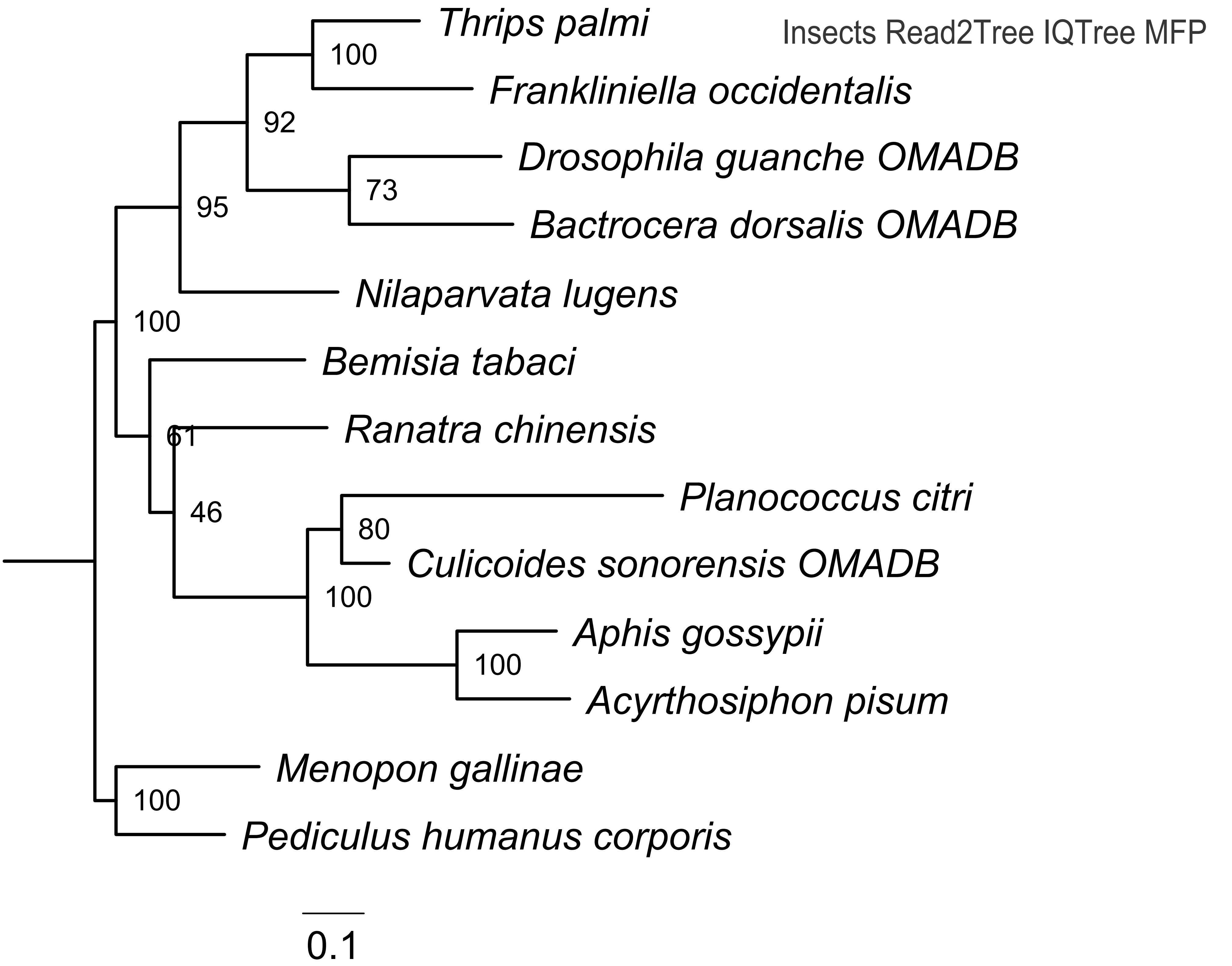

Insects Read2Tree IQTree MFP
Thrips palmi
Frankliniella occidentalis
Drosophila guanche OMADB
Bactrocera dorsalis OMADB
Nilaparvata lugens
Bemisia tabaci
Ranatra chinensis
Planococcus citri
Culicoides sonorensis OMADB
Aphis gossypii
Acyrthosiphon pisum
Menopon gallinae
Pediculus humanus corporis
100
92
73
95
100
61
46
80
100
100
100
0.1

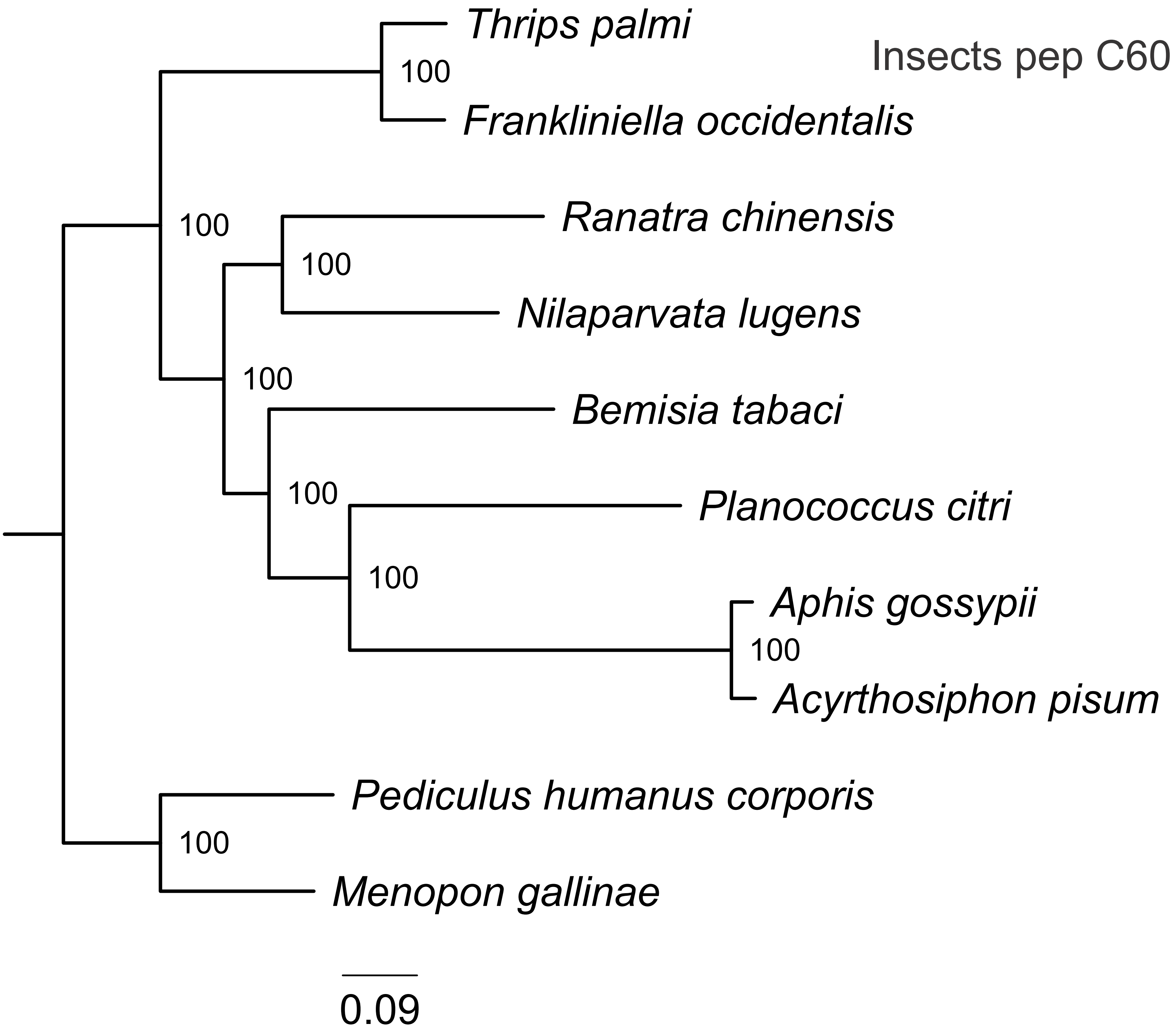

Insects pep C60
Thrips palmi
Frankliniella occidentalis
Ranatra chinensis
Nilaparvata lugens
Bemisia tabaci
Planococcus citri
Aphis gossypii
Acyrthosiphon pisum
Pediculus humanus corporis
Menopon gallinae
100
100
100
100
100
100
100
100
0.09

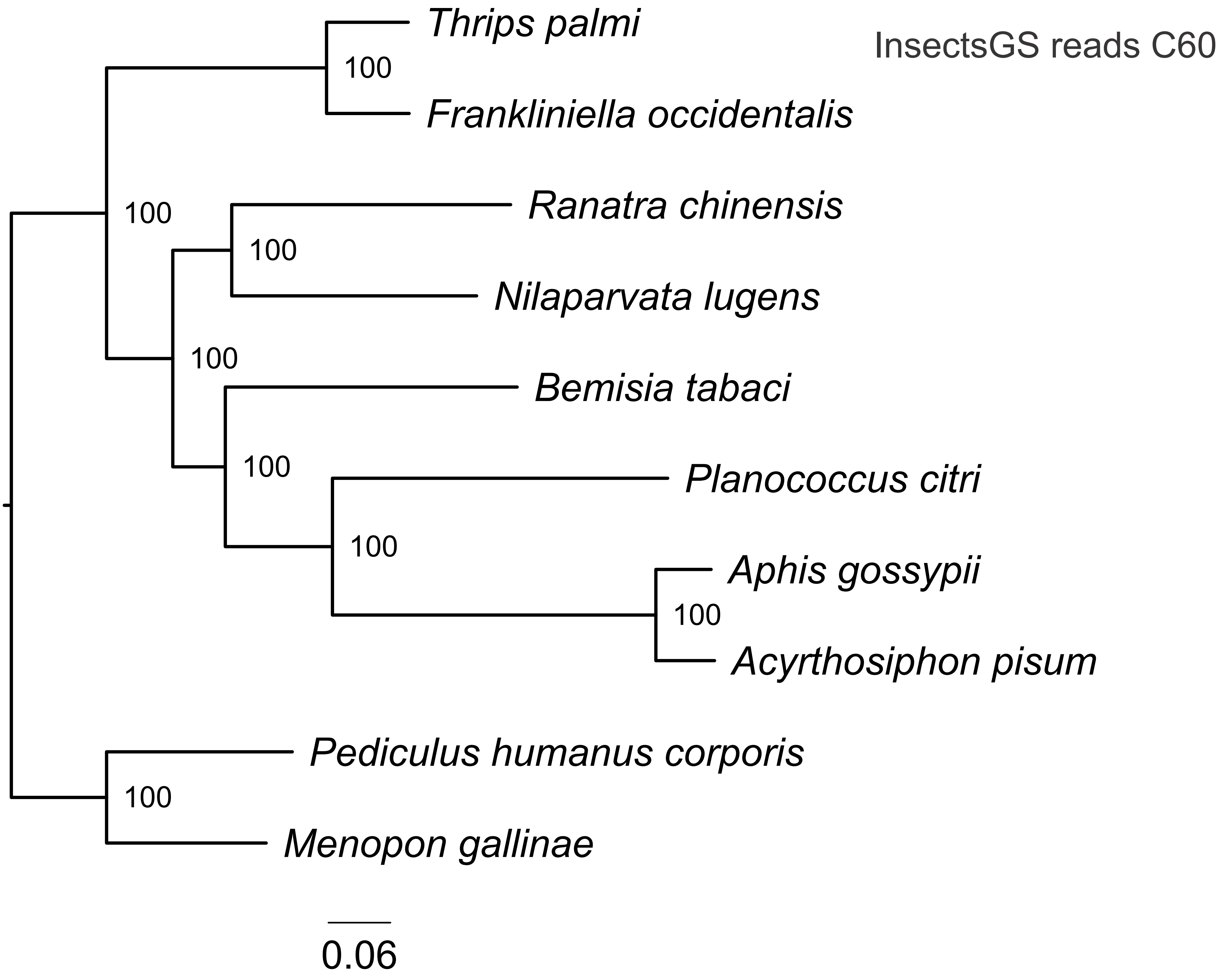
Thrips palmi
Frankliniella occidentalis
Ranatra chinensis
Nilaparvata lugens
Bemisia tabaci
Planococcus citri
Aphis gossypii
Acyrthosiphon pisum
Pediculus humanus corporis
Menopon gallinae
100
100
100
100
100
100
100
100
InsectsGS reads C60
0.06

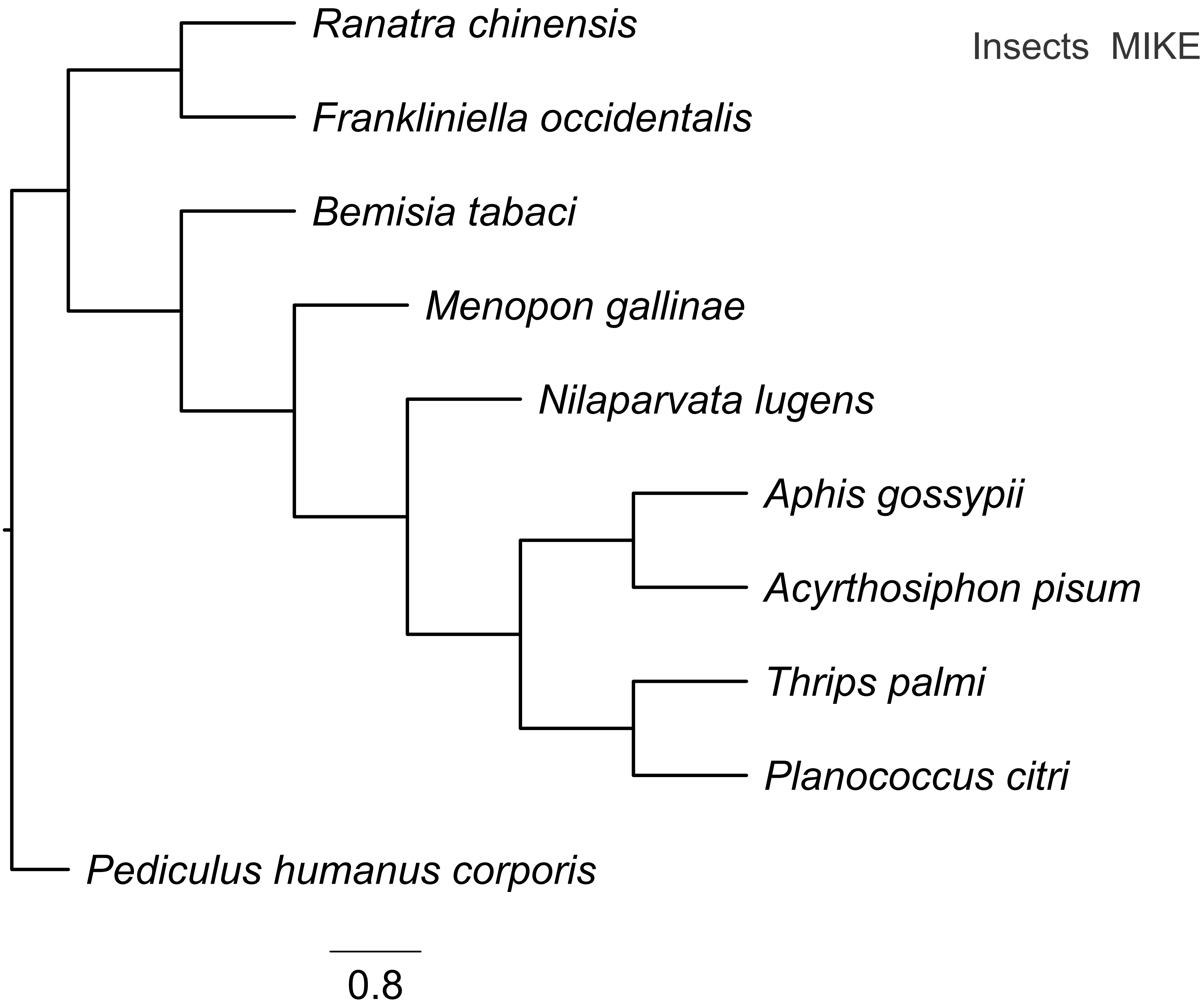
Insects MIKE
Ranatra chinensis
Frankliniella occidentalis
Bemisia tabaci
Menopon gallinae
Nilaparvata lugens
Aphis gossypii
Acyrthosiphon pisum
Thrips palmi
Planococcus citri
Pediculus humanus corporis
0.8

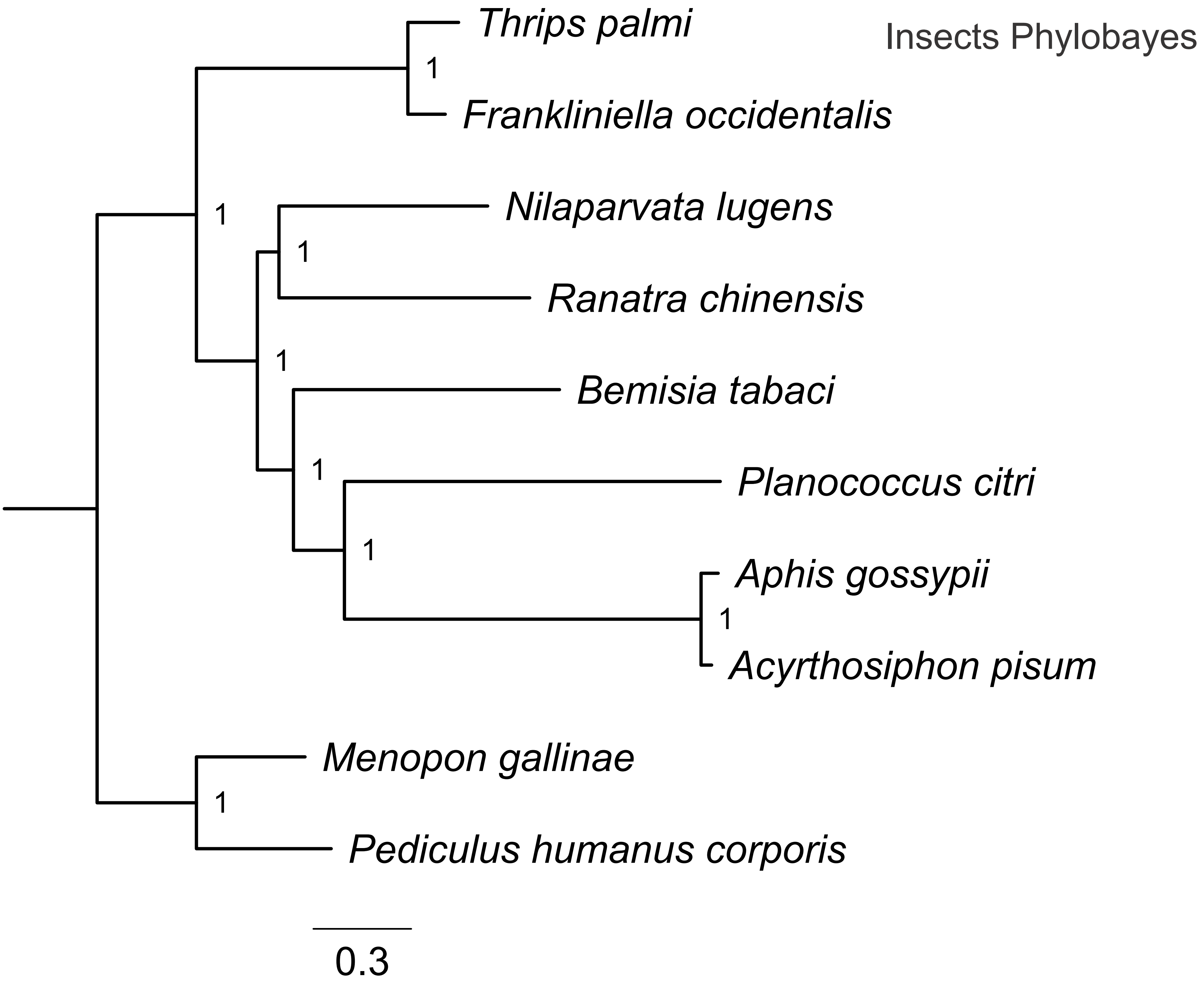

Insects Phylobayes
Thrips palmi
1
Frankliniella occidentalis
1
Nilaparvata lugens
1
Ranatra chinensis
1
Bemisia tabaci
1
Planococcus citri
1
Aphis gossypii
1
Acyrthosiphon pisum
Menopon gallinae
1
Pediculus humanus corporis
0.3
