## Supplementary Fig. 8 for "Reliable inference of phylogenomic relationship via assembly-based strategy accommodating raw reads and proteins"

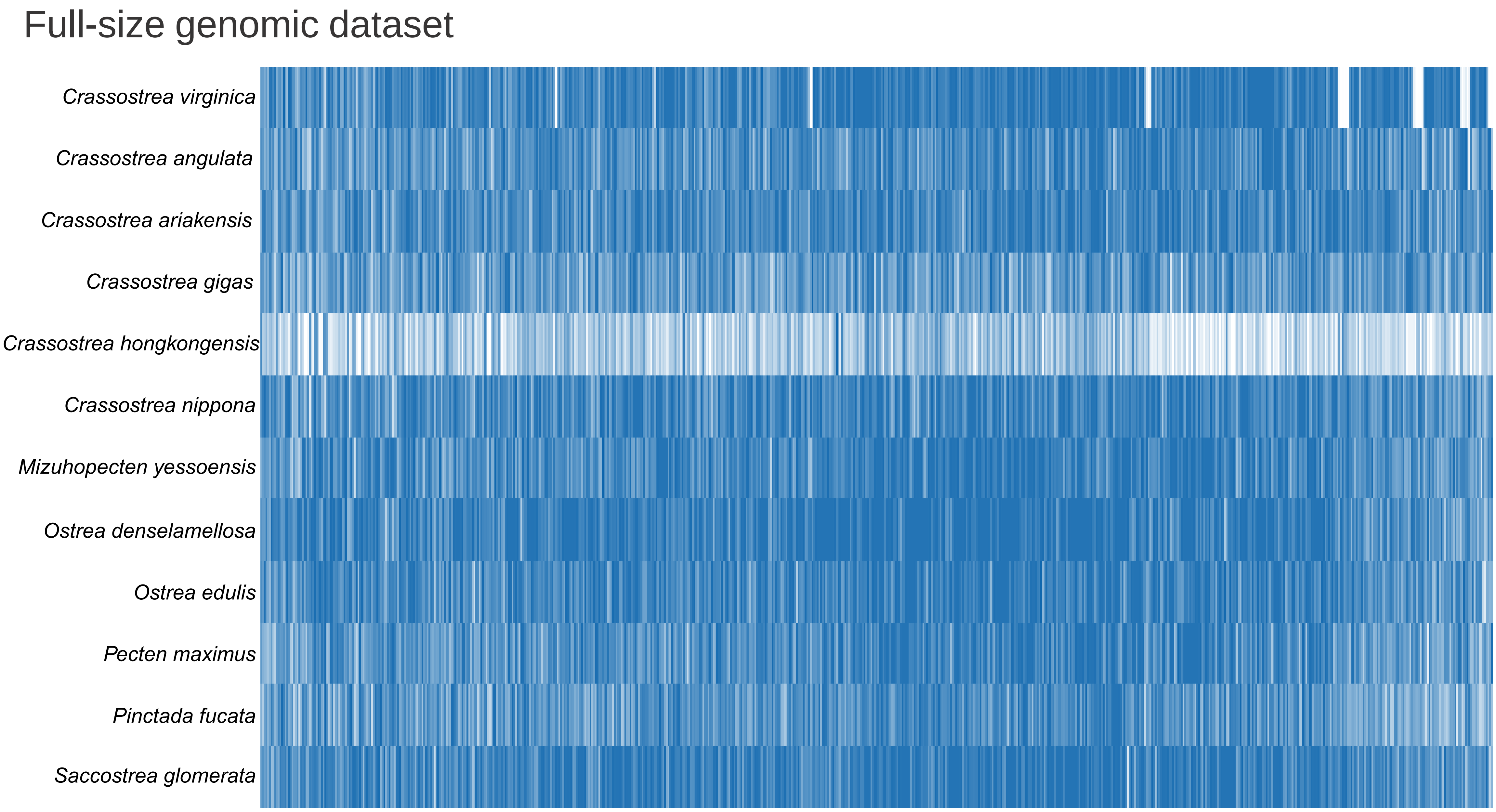
Full-size genomic dataset
Crassostrea virginica
Crassostrea angulata
Crassostrea ariakensis
Crassostrea gigas
Crassostrea hongkongensis
Crassostrea nippona
Mizuhopecten yessoensis
Ostrea denselamellosa
Ostrea edulis
Pecten maximus
Pinctada fucata
Saccostrea glomerata

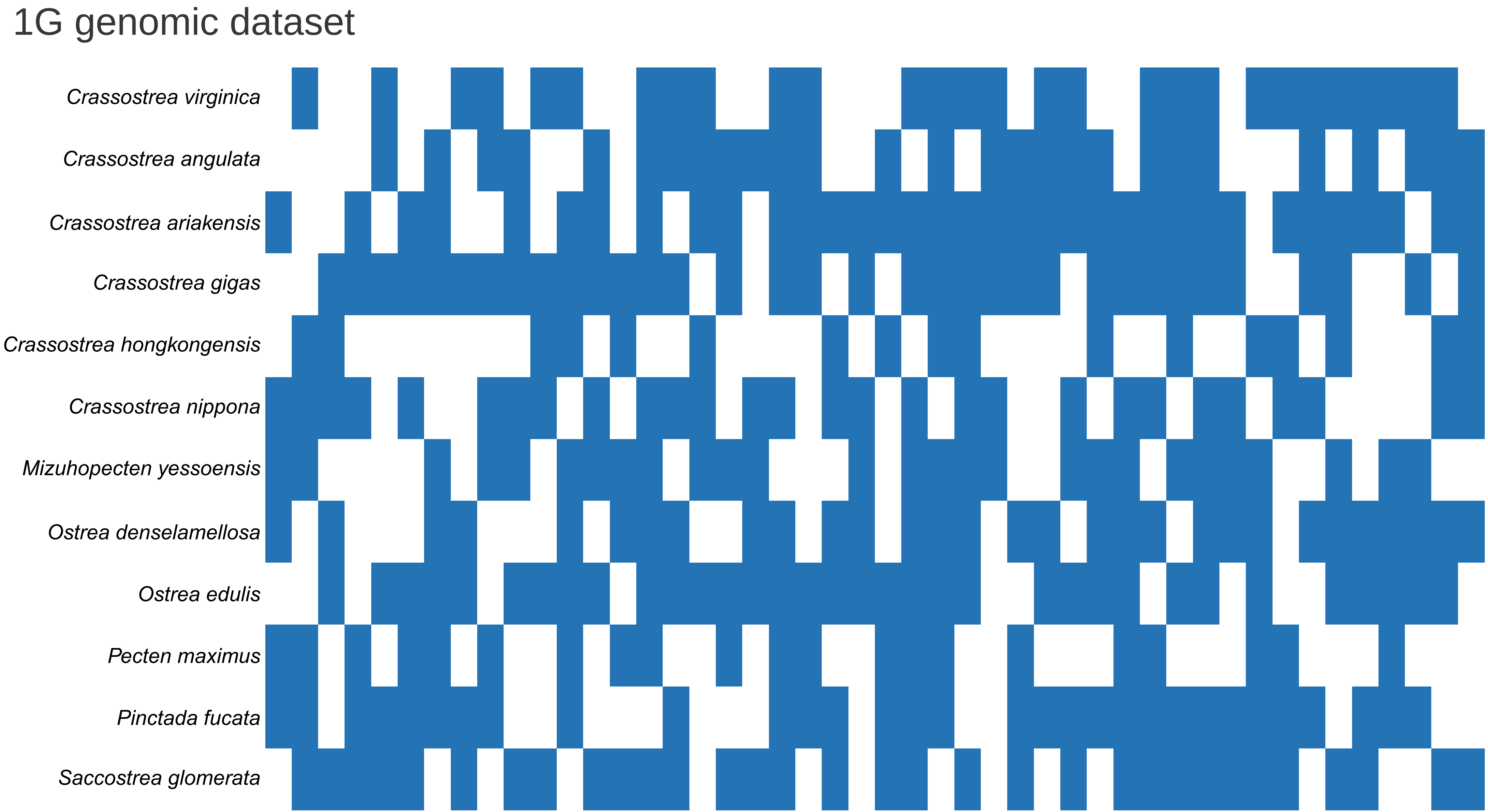

1G genomic dataset
Crassostrea virginica
Crassostrea angulata
Crassostrea ariakensis
Crassostrea gigas
Crassostrea hongkongensis
Crassostrea nippona
Mizuhopecten yessoensis
Ostrea denselamellosa
Ostrea edulis
Pecten maximus
Pinctada fucata
Saccostrea glomerata

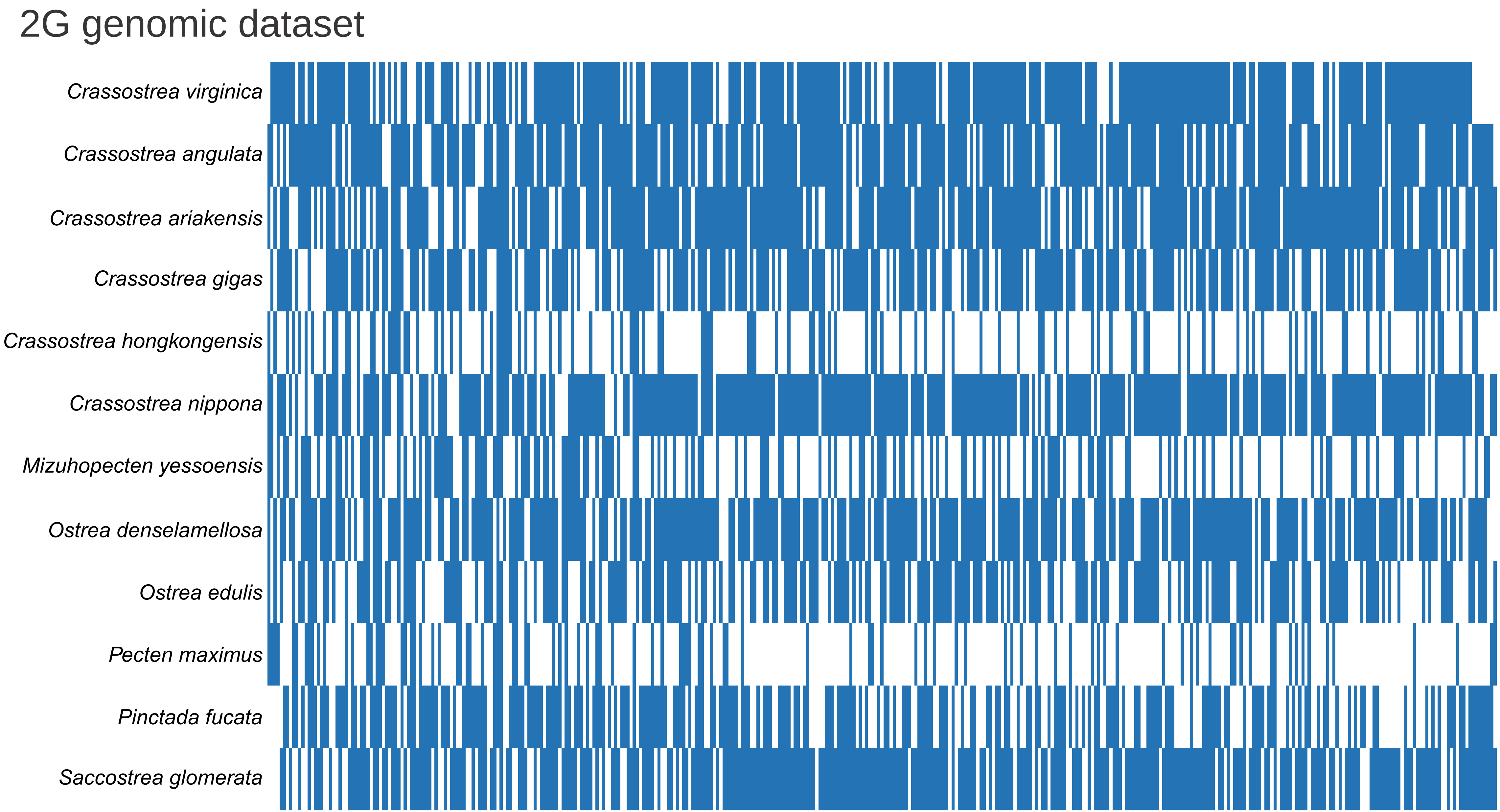
2G genomic dataset
Crassostrea virginica
Crassostrea angulata
Crassostrea ariakensis
Crassostrea gigas
Crassostrea hongkongensis
Crassostrea nippona
Mizuhopecten yessoensis
Ostrea denselamellosa
Ostrea edulis
Pecten maximus
Pinctada fucata
Saccostrea glomerata

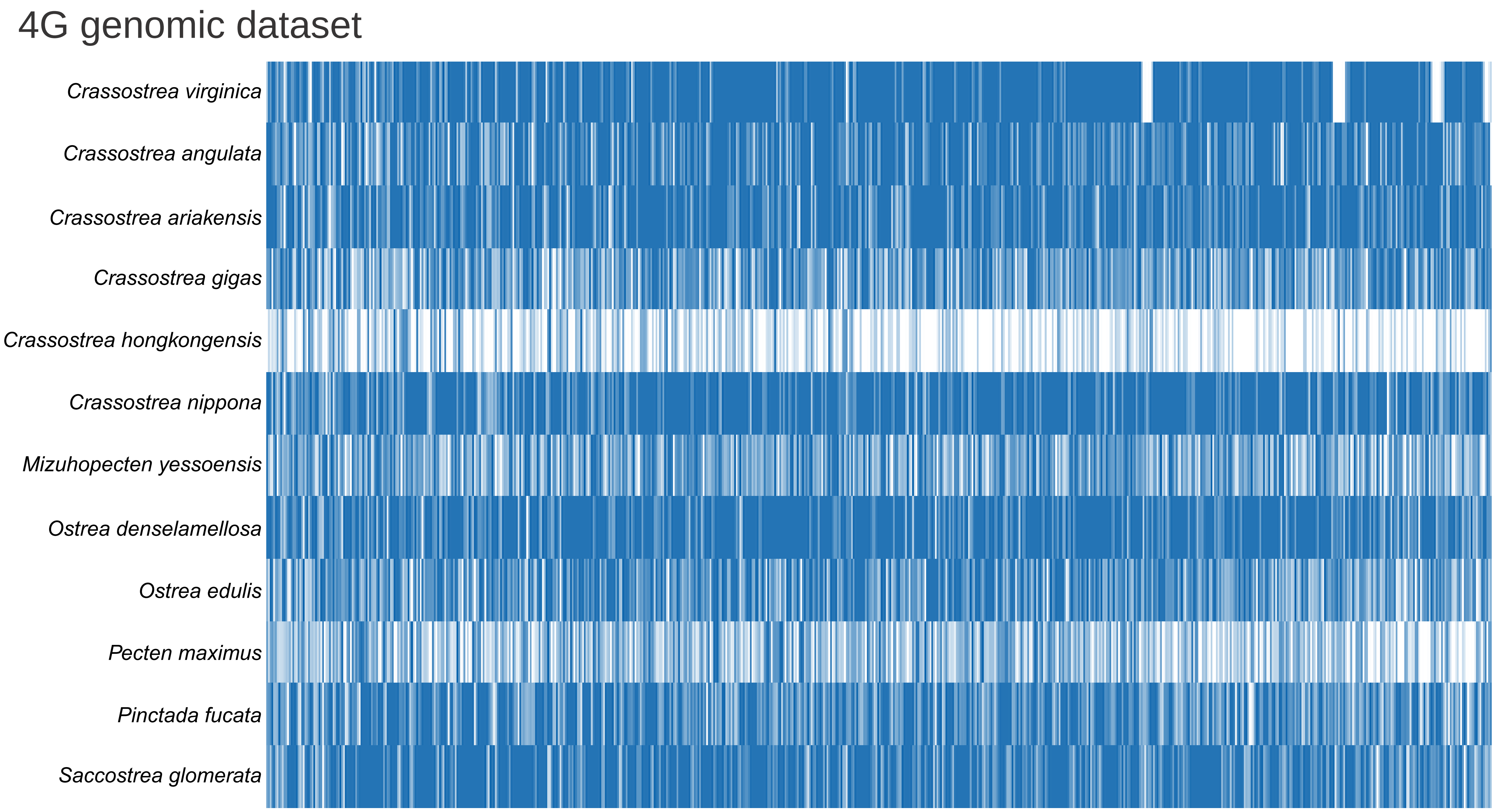
4G genomic dataset
Crassostrea virginica
Crassostrea angulata
Crassostrea ariakensis
Crassostrea gigas
Crassostrea hongkongensis
Crassostrea nippona
Mizuhopecten yessoensis
Ostrea denselamellosa
Ostrea edulis
Pecten maximus
Pinctada fucata
Saccostrea glomerata

### 6G genomic dataset

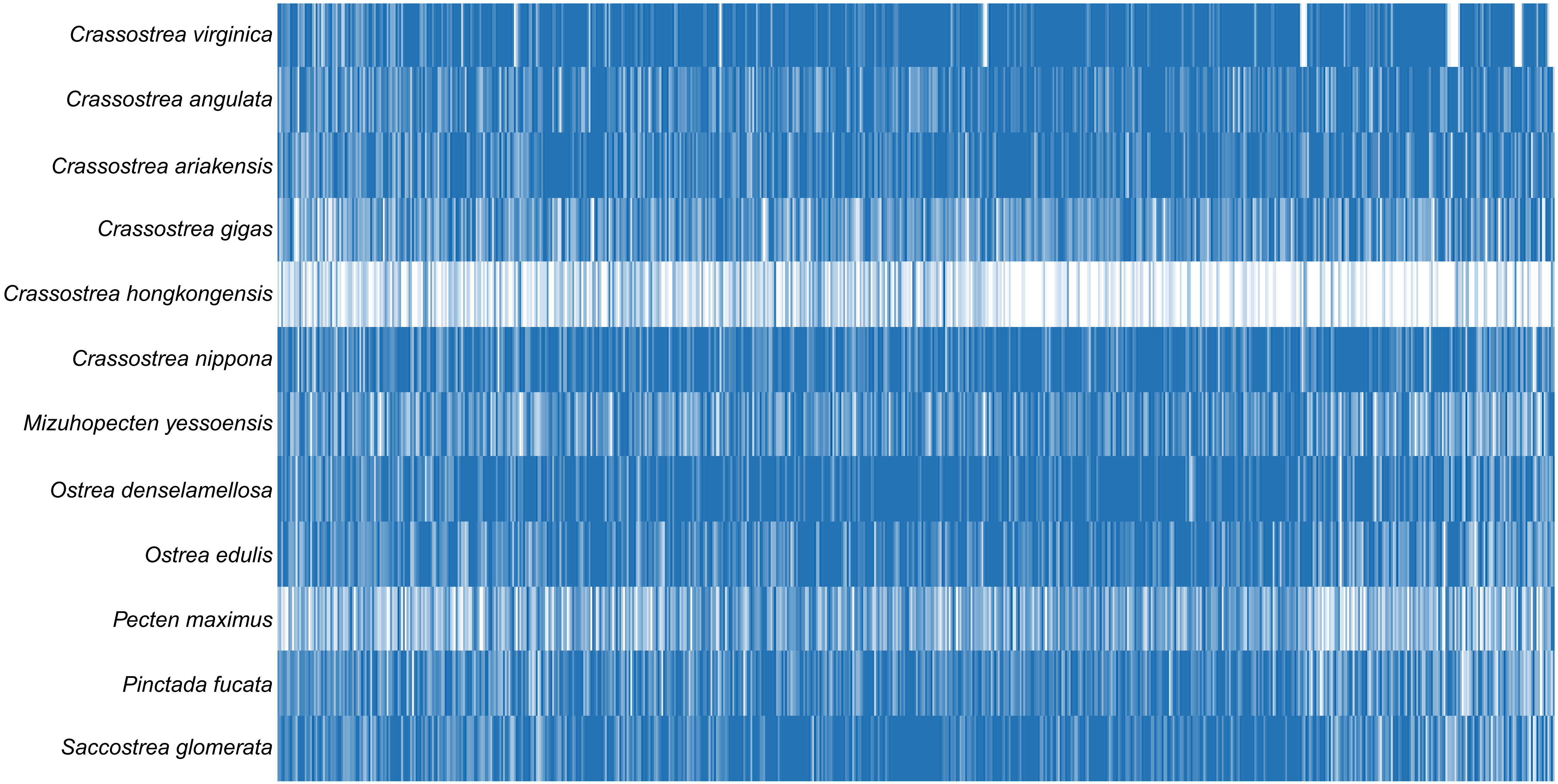

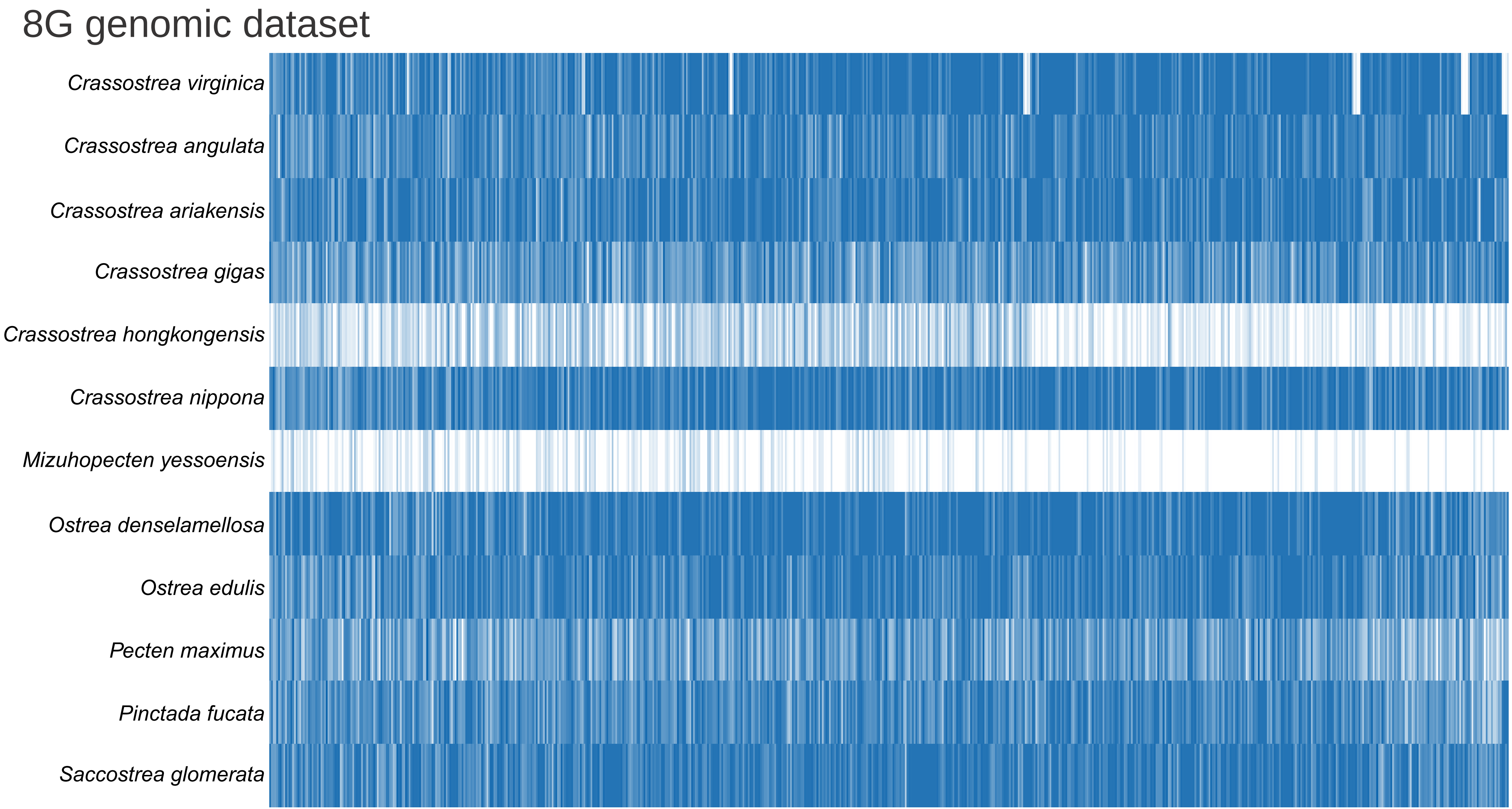
8G genomic dataset
Crassostrea virginica
Crassostrea angulata
Crassostrea ariakensis
Crassostrea gigas
Crassostrea hongkongensis
Crassostrea nippona
Mizuhopecten yessoensis
Ostrea denselamellosa
Ostrea edulis
Pecten maximus
Pinctada fucata
Saccostrea glomerata

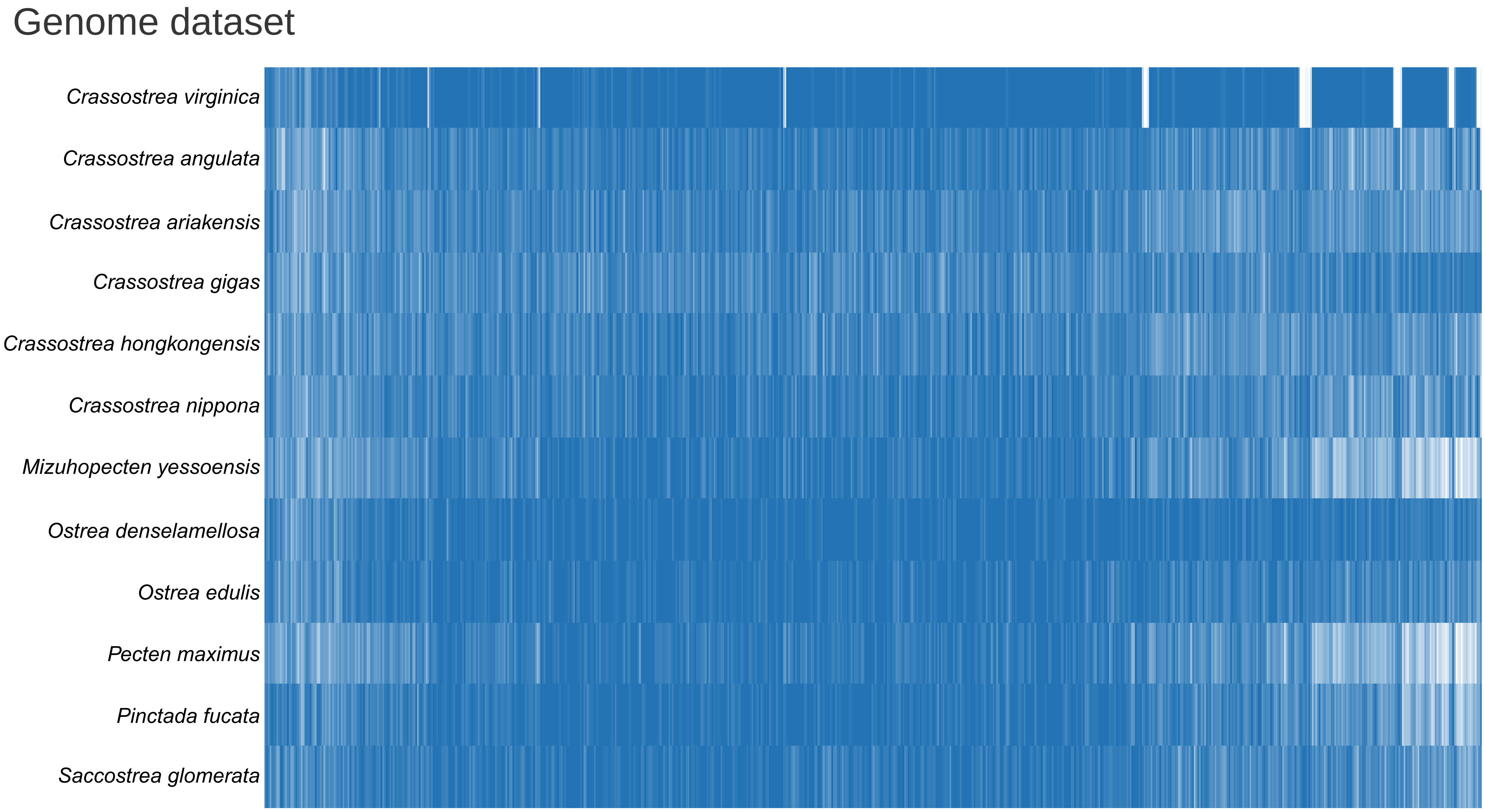
Genome dataset
Crassostrea virginica
Crassostrea angulata
Crassostrea ariakensis
Crassostrea gigas
Crassostrea hongkongensis
Crassostrea nippona
Mizuhopecten yessoensis
Ostrea denselamellosa
Ostrea edulis
Pecten maximus
Pinctada fucata
Saccostrea glomerata

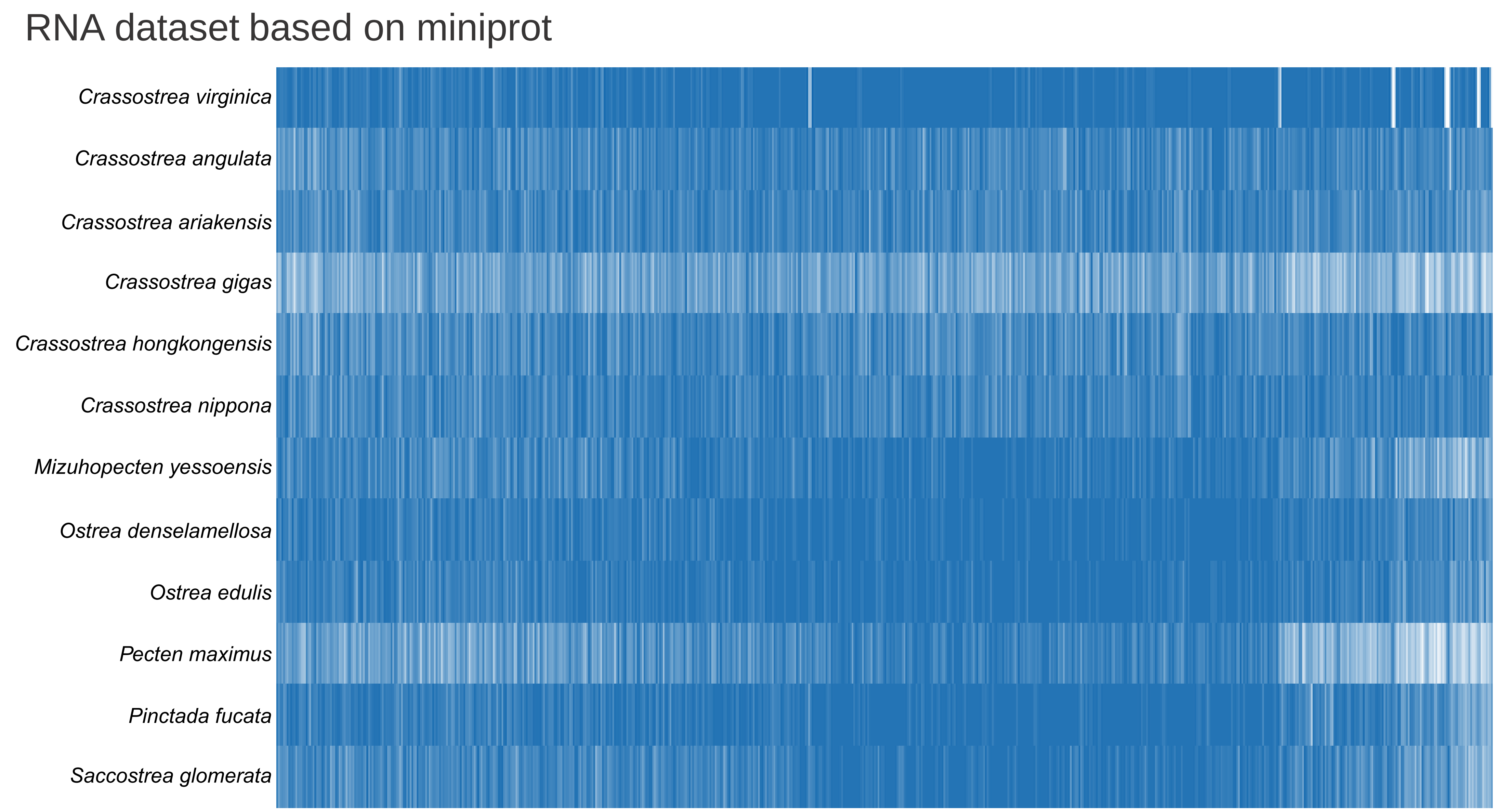
RNA dataset based on miniprot
Crassostrea virginica
Crassostrea angulata
Crassostrea ariakensis
Crassostrea gigas
Crassostrea hongkongensis
Crassostrea nippona
Mizuhopecten yessoensis
Ostrea denselamellosa
Ostrea edulis
Pecten maximus
Pinctada fucata
Saccostrea glomerata

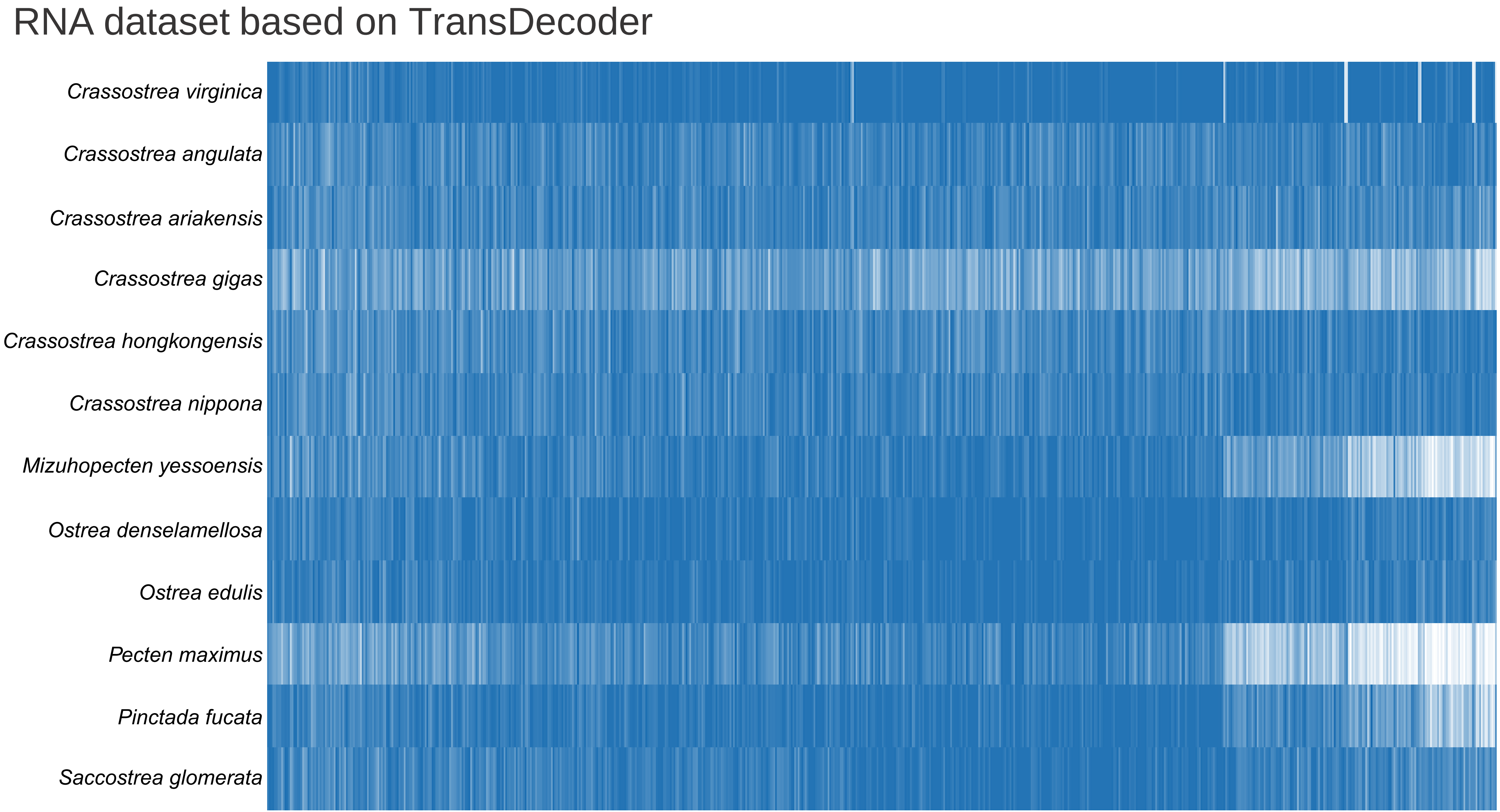
RNA dataset based on TransDecoder
Crassostrea virginica
Crassostrea angulata
Crassostrea ariakensis
Crassostrea gigas
Crassostrea hongkongensis
Crassostrea nippona
Mizuhopecten yessoensis
Ostrea denselamellosa
Ostrea edulis
Pecten maximus
Pinctada fucata
Saccostrea glomerata

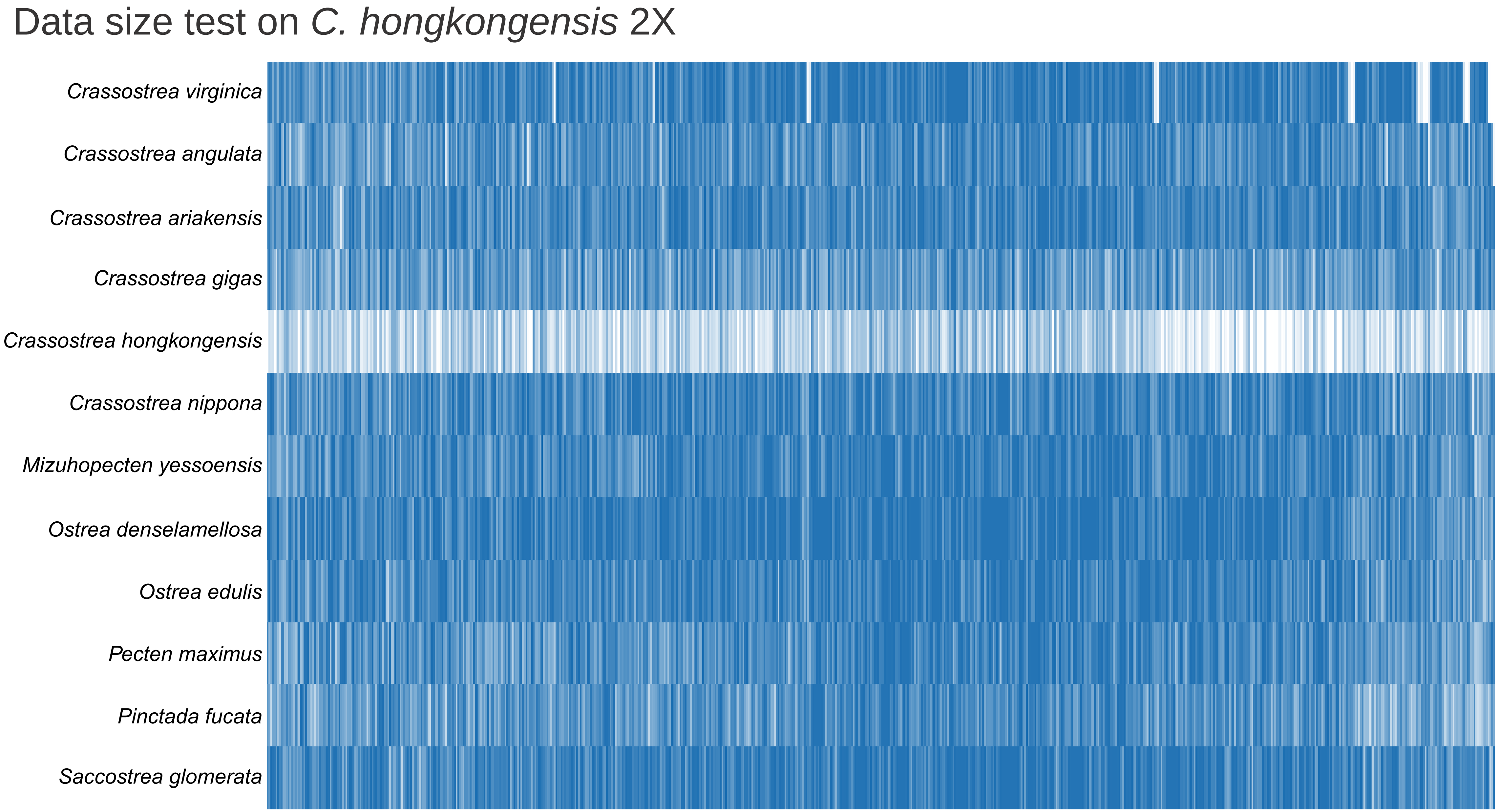
Data size test on C. hongkongensis 2X
Crassostrea virginica
Crassostrea angulata
Crassostrea ariakensis
Crassostrea gigas
Crassostrea hongkongensis
Crassostrea nippona
Mizuhopecten yessoensis
Ostrea denselamellosa
Ostrea edulis
Pecten maximus
Pinctada fucata
Saccostrea glomerata

Data size test on *C. hongkongensis* 4X

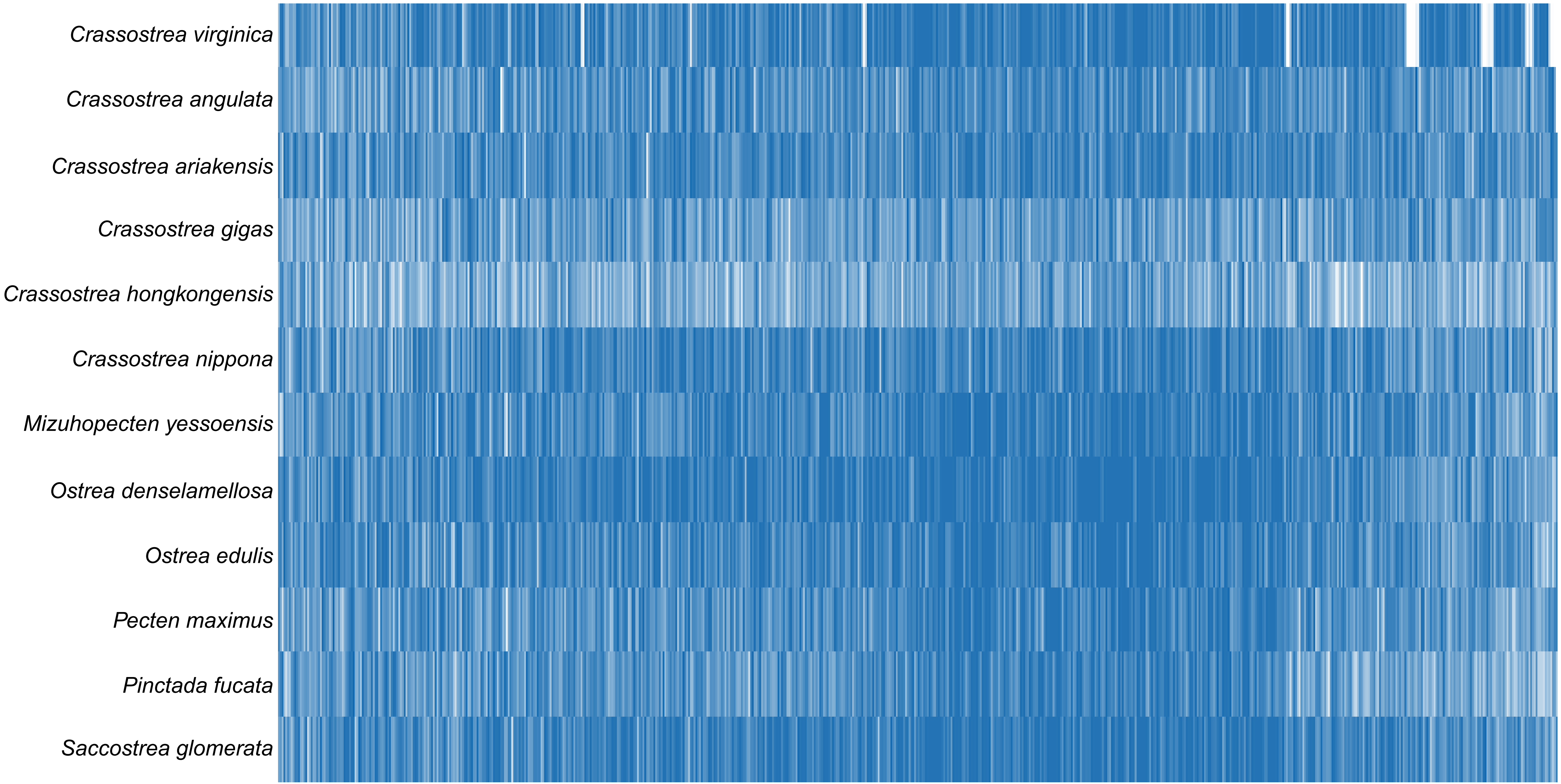

### Data size test on *C. hongkongensis* 6X

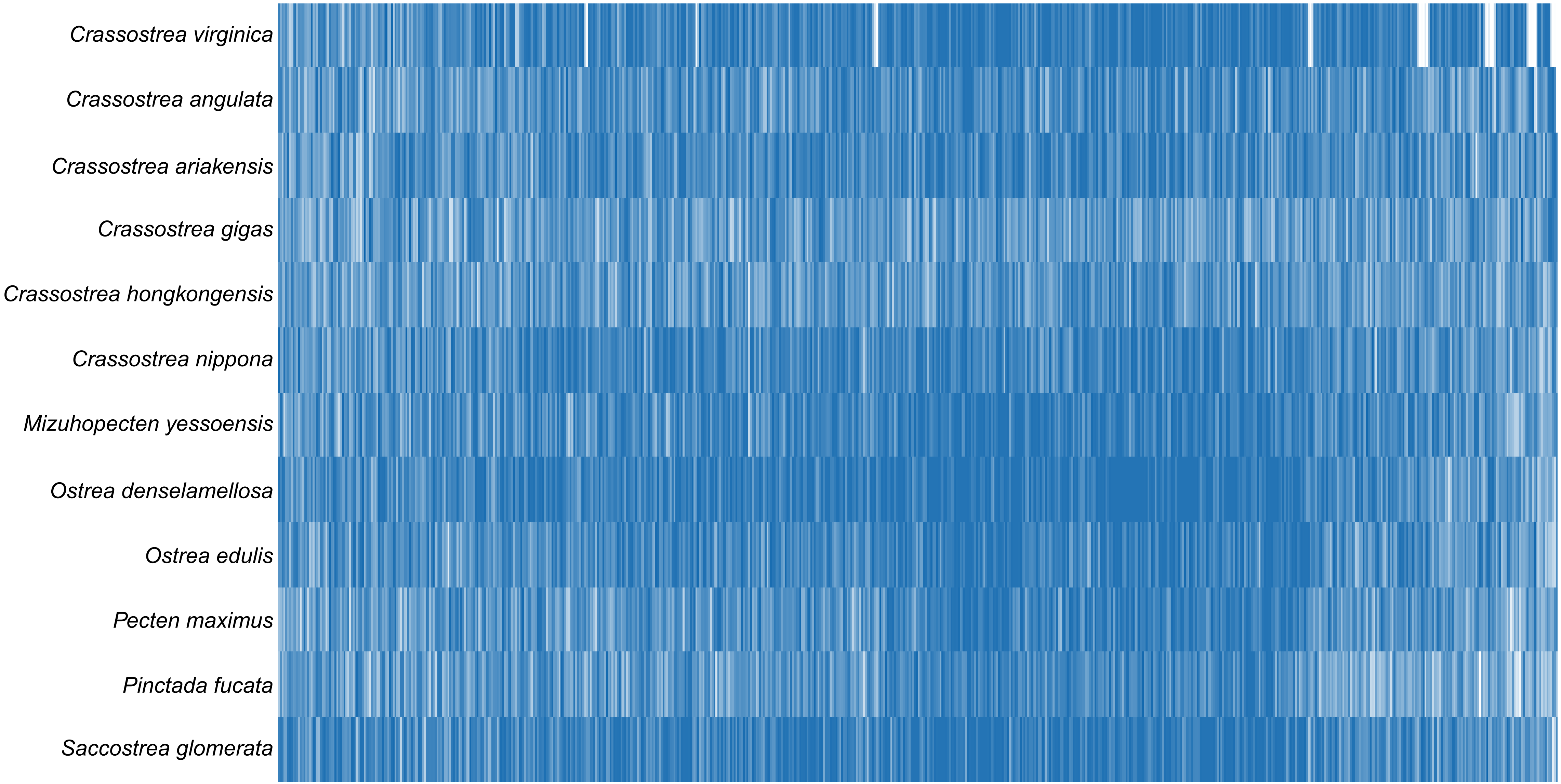

Data size test on *C. hongkongensis* 8X

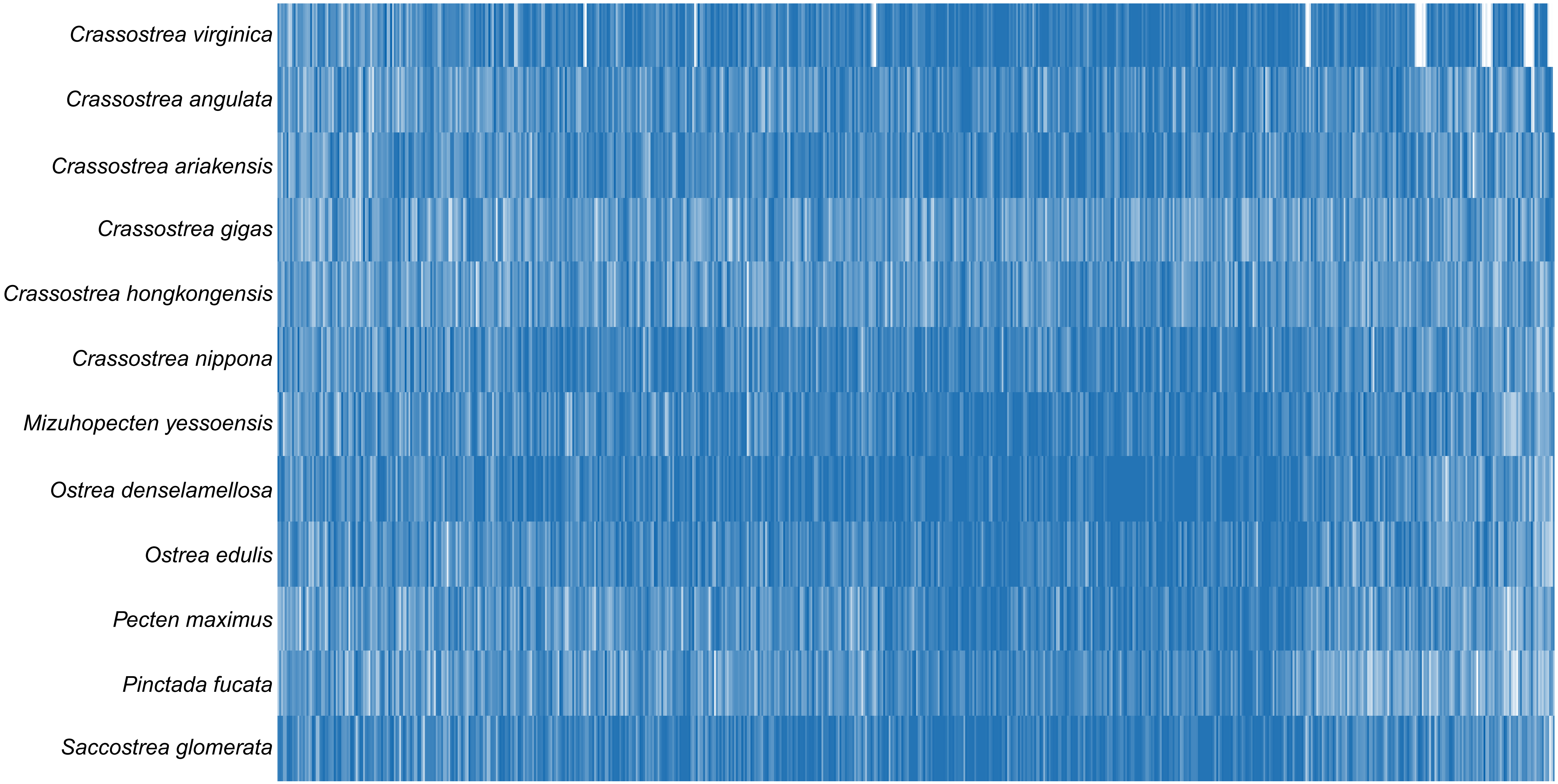

#### Neomphalida dataset

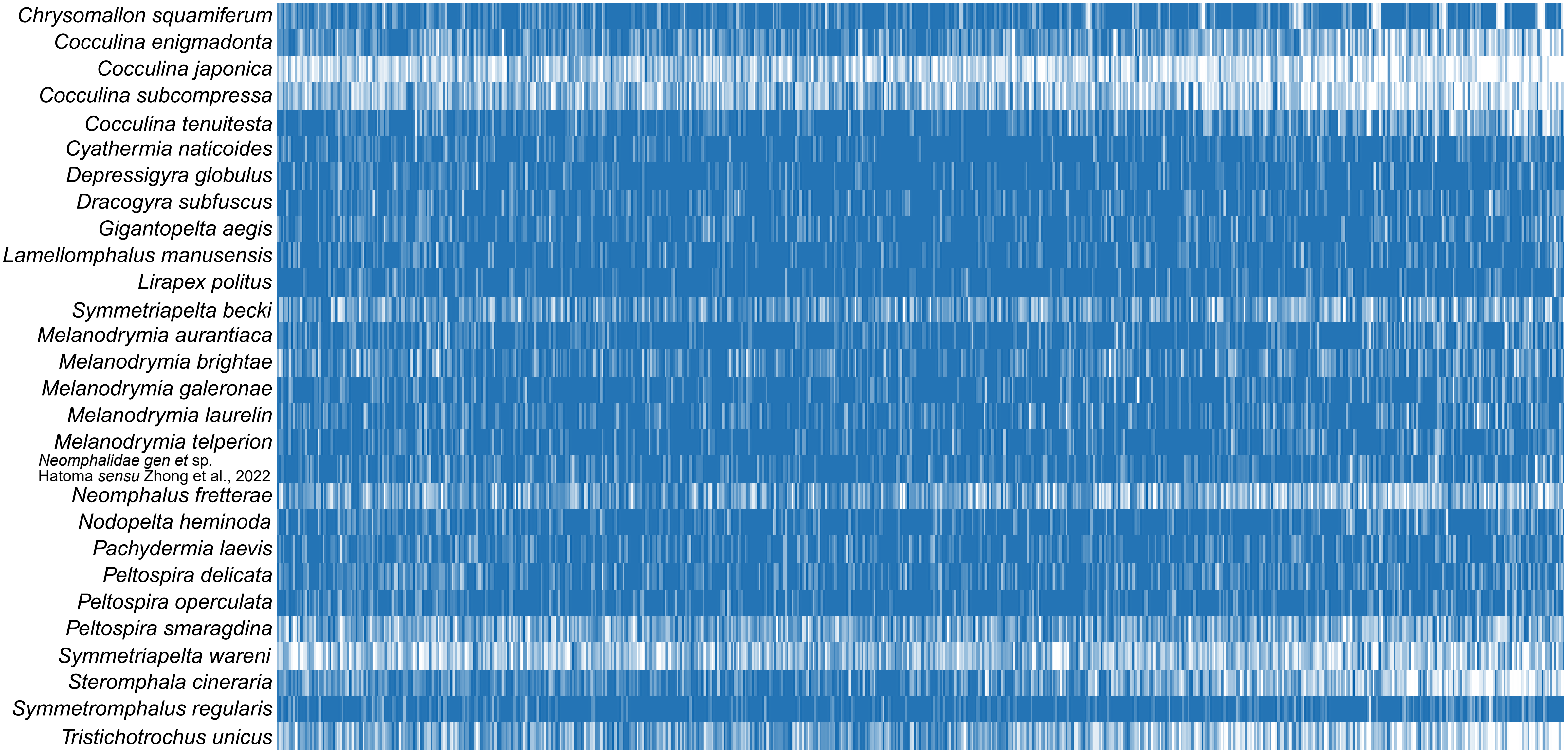
